# An Accurate Genetic Clock, *GENETICS*

**DOI:** 10.1101/026286

**Authors:** David H. Hamilton

**Affiliations:** Department of Mathematics, University of Maryland, College Park

**Author notes:** Department of Mathematics, University of Maryland, College Park, md 20742.

**Keywords:** Molecular clock, TMRCA, SNP, STR

## Abstract

Our method for “Time to most recent common ancestor” TMRCA of genetic trees for the first time deals with natural selection by apriori mathematics and not as a random factor. Bioprocesses such as “kin selection” generate a few overrepresented “singular lineages” while almost all other lineages terminate. This non-uniform branching gives greatly exaggerated TMRCA with current methods. Thus we introduce an inhomogenous stochastic process which will detect singular lineages by asymmetries, whose “reduction” then gives true TMRCA. This gives a new phylogenetic method for computing mutation rates, with results similar to “pedigree” (meiosis) data. Despite these low rates, reduction implies younger TMRCA, with smaller errors. We establish accuracy by a comparison across a wide range of time, indeed this is only y-clock giving consistent results for 500-15,000 ybp. In particular we show that the dominant European Y-haplotypes R1a1a & R1b1a2, expand from c4000BC, not reaching Anatolia before c3800BC. This contradicts previous clocks dating R1b1a2 to either the Neolithic Near East or Paleo-Europe. However our dates match R1a1a & R1b1a2 found in Yamnaya cemetaries of c3300BC by Nielsen et al (2015), Pääbo et al(2015), together proving R1a1a & R1b1a2 originates in the Russian Steppes.

## Introduction

Zuckerkandl & Pauling (1962) made the empirical observation that “time to most recent common ancestor” (TMRCA) of a genetic tree seemed proportional to genetic variance. Thus began the molecular clock, a recent account can be found in Wakeley(2013): a very short list of applications might be the divergence of the eukaryotes (Kimura and Ohta, 1973)… the origins of human immunodeficiency virus (Korber et al., 2000). The Y(chromosome) clock also gives an important tool of archeology since pioneering work by L.L. Cavalli-Sforza et al(1978) correlated genetic variances to age of human lineages. Although in broad agreement with the fossil record or ^14^C dating, very large confidence intervals meant that questions such as “peopling the Americas” or the origin of the dominant European Y-haplotype R1b1a2 could not be accurately resolved.

Moto Kimura’s theory of neutral mutations(1968) predicted a molecular clock with constant rate. This has been applied in many versions, we call KAPZ, see Nielsen(2005). Actual timing requires calibration of mutation rates, either by counting genetic differences over a lineage, i.e the phylogenetic method, or by direct observation of parent-offspring pairs, i.e. meiosis. For the human Y-chromosome Burgella and Navascue(2011) combined 80 meiosis studies, which for our 33 markers averages at about .0043 mutations per generation, while their alternative regression method averages at .0026. Chandler(2006) combines 20 meiosis studies to get an average of .0032. Unfortunately these rates when used in KAPZ do not correlate very well with known genealogy. So FTDNA used their huge data base to give an average of .00635. Indeed for the Y-clock in general “phylogenetic” rates are about 2 times larger than those from meiosis. While this gives plausible results for genealogy it makes ancient lineages seem far too young. Zhivotovsky (2000) conceived of a much smaller “long term evolutionary rate", computing on average .0007 from genetic lineages of remote Polynesian populations. Myres et al (2011) used this in a KAPZ giving 9000BC for the origin of R1b1a2, i.e. right after the Ice Age.

However a long term constant rate has proved untenable as estimating rates from different genetic lineages of known ages gave rates with significant discrepancies, see Nie and Kumar(2000). Its variability was attributed to changing generation times, metabolism etc. and of course natural selection, see Ohta(2002), Lynch(2010). Some sophisticated empirical approaches allowing variable rates have been developed, see Nielsen(2005). However confounding examples convinced Ayala(1997) to challenge the very idea of molecular clock hypothesis, reasoning that variable rates are the result of the unpredictable events of natural selection. Never the less, while future natural selection cannot be predicted, we show how singular events of the past can be detected.

Human populations have been classified by Y(chromosome) haplotypes defined by a SNP (single nucleotide polymorphism) mutation, a famous example being R1b1a2. Now one is looking at mutations of “short tandem repeat” (STR) at different **D**NA **Y**-chromosome **S**egments (DYS). In applications of the Y-clock researchers have stuck with constant mutation rates. However, comparing rates at the same DYS markers we see an average 50% variation over eight major Y-haplotypes, i.e the same phenomena seen for the very long term allele clock. Now as a huge variation in human generation times etc over recent time does not seem plausible reseachers have fallen back on the stochastic process itself to generate unlikely distributions. These Bayesian methods use Monte-Carlo simulations of all possible genealogical trees giving the present sample data, then find TMRCA by a maximum likehood estimate. An example of this for the Y-clock is the algorithm BATWING (Wilson et al, 2003). Now for BATWING the TMRCA is often greater than the KAPZ, e.g. for the Cinnioglu et al(2004) study of Anatolian R1b1a2 both methods were applied to the same data and mutation rates. The KAPZ gives TMRCA 9800BC compared with 18,000BC for BATWING. Balaresque et al (2010) used BATWING to give an origin for R1b1a2 in Neolithic Anatolia c.6000BC, however disputed by Busby (2012). Another Bayesian algorithm is BEAST, see Drummond and Rambaut(2007), which has been used for mitochondrial clock where finding the phylogenetic tree is easier. Unequal fertility can be represented by non-uniform branching but in Bayesian Models being less probable increases TMRCA.

Instead, we deal with natural selection by apriori mathematics and not as a random factor. Bioprocesses such as “kin selection” generate a few overrepresented “singular lineages” while almost all other lineages terminate. These singular lineages are extremely unlikely so cannot be predicted. Instead we introduce an inhomogenous stochastic process where we have constant mutation rates but variable sources representing nonuniform fertility, i.e. many virtual patriarchs giving a quasi-mixed population. To find these sources is an inverse problem, however inversion is not stable or even unique: the population could have been created yesterday! However we are saved by finding the present population is dominated by a relatively few “singular lineages". Our model has many extinct twigs with a few successful branches. In particular about 50% of the DYS markers show a singular side branch. This requires a new technique: singular branches can be detected by asymmetries in the present distribution (taking mutation bias into account), then *reduced* revealing the original TMRCA.

We account for the tendency of bioprocesses to exaggerate the apparent mutation rate by using *reduction of singular lineages* to give a new phylogenetic method for computing mutation rates. This gives an average of .0031 mutations per generation, right between the meiosis results of Burgella and Navascue, Chandler. Indeed our theoretical phylogenetic rates are similar to actual laboratory meiosis rates, see Table 8. Despite our low mutation rates, “reduction of singular lineages” implies younger TMRCA. Also our low SD verifies the Kimura theory that rates are constant, at least for the human Y-Chromosome for 500 - 15,000ybp. However the large difference between rates, up to a factor of 100, implies that in the very long term that these may vary, probably as the geometry of the genome evolves. Furthermore we shall see that the essential equality of our rates with those from meiosis implies *a priori*, a generation length of 30 years. This agrees with Fenner(2005) who found a 30 year generation from field work among Hunter Gatherers. All of this provides theoretical validation.

However theoretically plausible the true test is accuracy, indeed our original motivation was to develop a Y-clock with accuracy approaching ^14^C dating. This is verified by a comparison across a wide range of time, the first STR-clock giving consistent results for 500-15,000 ybp with accuracy about 11% (one SD). The usual sample error, uncertainty in mutation rates contributes to the total accuracy. However ~ 1/2 is inherent in the method itself, this being determined by large scale simulations of random trees with known TMRCA. It is easy to forgot to include this, e.g. rarely computed for KAPZ methods. Comparison of 7, 15, 29 markers leads us to expect about 5% (one SD) for 120 markers.

We show our dates correlate with European history. In particular the dominant European Y-haplotype R1b1a2, expands from c4000BC, not reaching Anatolia before c3800BC. This contradicts previous clocks dating R1b1a2 to either the Neolithic Near East or Paleo-Europe. However our dates match R1b1a2 found in Yamnaya cemetaries of c3300BC, see Ramus Nielsen et al(2015), Svante Pääbo et al(2015). Our times and their places almost certainly implies R1b1a2 originates in the Russian Steppes, disproving the “out of Anatolia hypothesis” and instead giving strong evidence for the Indo-European hypothesis of Marija Gimbutas(1956). A different example is the dominant Amerindian Y-haplotype, Q1. Bortolini et al(2003) used 8 markers to give a TMRCA =11600BC (SD3000). Using recent data of Battaglia et al (2013) we find the TMRCA =13300BC with a 90% chance of occurring before the Canadian Ice Corridor opens, i.e. the initial migration was along the coast.

Very recently Batini et al (2015) computed TMRCA using the algorithm BEAST, with mutation rates for MYS sites on the Y-chromosome found from Icelandic genealogy by Helgason et al (2015). This is the complete opposite of our method, using *~* 3 × 10^6^MSY sites compared with our 29 DYS, but samples sometimes as small as *N* = 8 while we have *N* ~ 1000. We compare all results in TABLE 10, see appendix. Their TMRCA are generally similar to ours, so together we support recent dates for R1b1a2 (and also verifying the Icelandic rates). For example Batini finds that R1b1a2 has TMRCA= 3550BC, compared with our 4000BC. But their sample of R1b1a2 (unlike ours) is M246*, i.e. major branches like L11, P312 are removed. So there is little branching and their results using BEAST and a KAPZ method RHIO are very similar. However when there is large scale branching our theory predicts their Bayesian method will give older results. This is seen for the SNP J2 where our TMRCA is about 5000 years younger. Our dates for J2 and G2a2 correlate with their expansion from the Neolithic Revolution. Another example is for J2b, which is presently concentrated is Eastern Europe. Now Nielsen et al, Reich et al used ADMIXTURE to find the initial Indo-European expansion into Central Europe during the Corded Ware Culture (3000-2000BC) was mainly of Siberian type DNA but with significant Middle Eastern component not found in the original Yamnaya Culture, probably from mixing in the Western Urkraine. This correlates with our finding for J2b, TMRCA =3286BC (cf. 10,000BC for Batini).

## Singular Lineages

A fundamental problem is that present populations have highly overrepresented branches we call *singular lineages*. A well known example is the SNP L21 which is a branch of R1b1a2. Individuals identified as L21 are often excluded from R1b1a2 analysis because they skew the results. Such a singular lineage causes the variance to be much greater, even though the original *TMRCA* remains unchanged, see figure 1.

**Figure 1.**
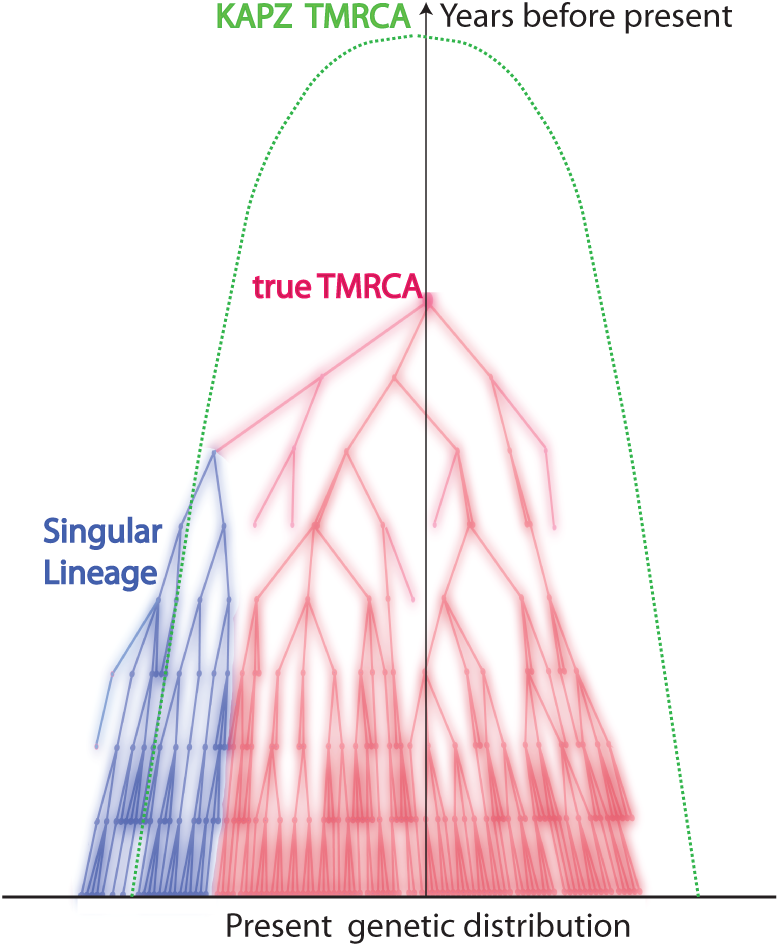
Singular Lineages in Random Trees

For Bayesian methods such lineages are very unlikely giving an even greater apparent *TMRCA*. However one cannot deal with singular branches by excluding them. For one thing, our method will show that 50% of markers show evidence of singular side branches, i.e. more than a SD from expected. For instance for a large sample of R1b1a2 we plotted the distribution about the mode for DYS markers GATA H4 and YCA IIa. For the later there is little difference between the theoretical distribution (red) and the actual data(blue), however for GATA H4 one sees the large discrepancy (see SI 1 for all 33 DYS distributions, in Supplement). The ideal distribution is not symmetric because we have also accounted for bias in the mutation process, see Wakeley(1994). These singular lineages are very (mathematically) unlikely to arise from the stochastic system which is the mathematical basis of KAPZ (or the equivalent Monte-Carlo process modeling Bayesian trees). We believe that the standard stochastic process is perturbed by “improbable” biological processes.

First, the Watson-Galton process introduced to explain the extinction of aristocratic lineages(see Kendall, 1975) implies lineages almost certainly die out. Conversely, natural selection causes some branches to flourish, e.g. the “kin selection” of W.D. Hamilton (1964), shows kin co-operation gives genetic advantages. Consider three examples with well developed DNA projects. Group A of the Hamiltons has approximately 100, 000 descended from a Walter Fitzgilbert c1330AD. Group A of the Macdonalds has about 700, 000 descendants from Somerfeld c1100AD, and Group A of the O’Niall has over 6 million descendants from Niall of the Seven Hostages, c300AD. These are elite groups with all the social advantages. One sees lines of chieftains, often polygamous. We emphasize kin selection because it seems dominant over natural selection for recent human lineages, certainly no one has claimed the O’Niall are genetically superior! Natural selection would cause similar branching over longer time scales. These singular lineages cannot be predicted by the stochastic model itself.

## Reduction of Singular Lineages

Although our method is for general molecular clocks to be specific we focus on the Y-clock. Consider **D**NA **Y**-chromosome **S**egments (DYS) counting the “short tandem repeat” (STR) number of nucleotides. One uses these DYS micro-satellites, marked by *j* = 1, …*n*, each individual *i*, 1 = 1, ¨*N*, has STR number *xi*,*j*. The Y-chromosome is unchanged except for mutations *xi*,*j → xi*,*j* ± 1 occurring at rate *μj* per generation.

Modelling singular lineages requires a new stochastic system. The standard system is: at time *t* the probability *P* that the DYS markers have given STR vector *X* is governed by a homogenous system of discrete diffusion equations

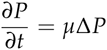

allowing variable mutation matrix *μ* = *μ* (*t*, *X*). Our model will have constant mutation rate matrix *μ* in a discrete inhomogenous equation

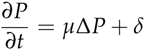

but with unknown sources *δ* = *δ* (*X*, *t*) modeling unequal fertility. Solving either system for unknown *μ* or *δ* is an inversion problem. But inversion is unstable for such systems, also there is no unique solution.

However in practise, up to a standard deviation, the in-homogenous system is a perturbation of the KAPZ system, *δ*(*X*, *t*) = 0, so that “singular branches” are clear enough to be be computed *a priori* without empirical considerations. These singular branches are then *reduced* revealing the original lineage.

We then compute a branching time *tj* for each DYS marker *j*. Now the nonuniform branching process causes the *tj* to be randomly distributed so their mean is not the TMRCA. Figure 3 for R1b1a2, shows the error spread for each *tj* over 29 DYS sites (note the outliers!). One can combine the distributions for *tj* give the “Branching Spectrum” (dotted curve). Now large errors and outliers means one cannot simply take the max *tj* to be the *TMRCA*. Instead stochastic simulations of the branching process, using robust statistics to avoid outliers, find the most likely *TMRCA*. The effect of reduction is dramatic, e.g. the TMRCA for R1b1a2 changes from 7150BC(KAPZ) to 4000BC after singular reduction, using the same markers and mutation rates, see Figure 4 for effect on the Branching Spectrum.

**Figure 2.**
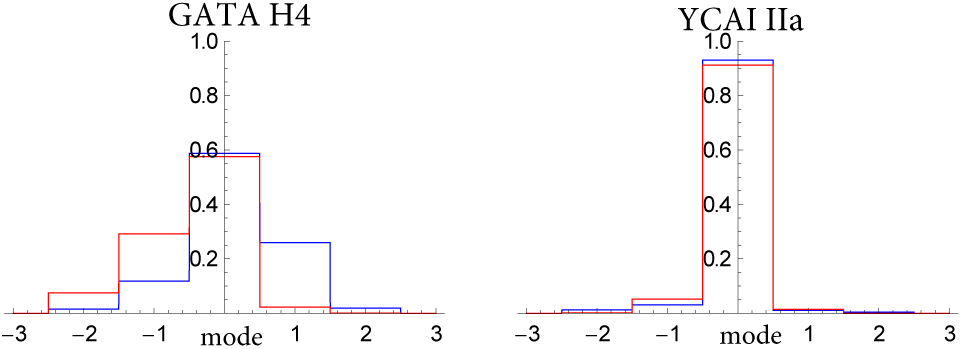
Biased distributions

**Figure 3.**
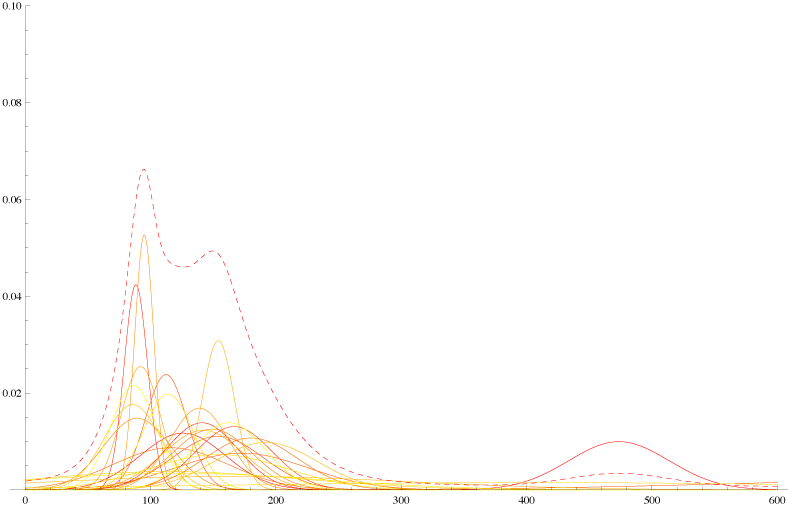
Distribution of branching times

**Figure 4.**
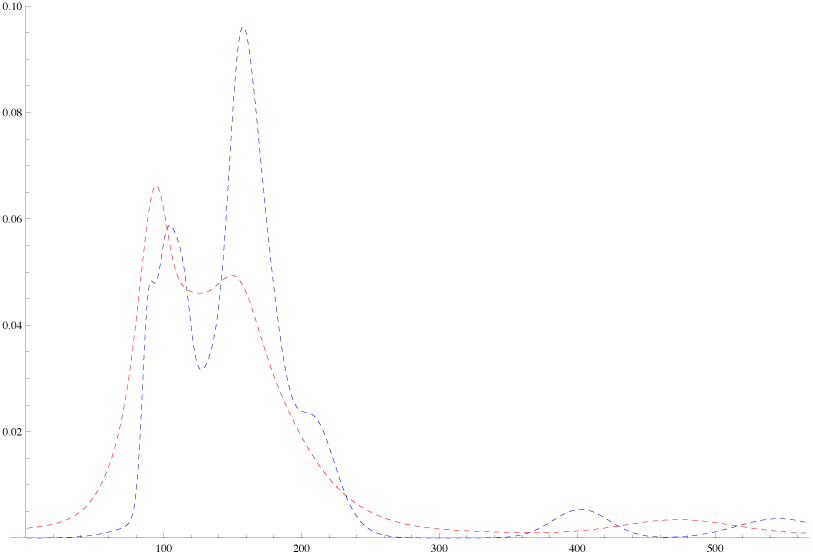
Branching Spectrum before reduction(blue) and after.

In SI 6 we display similar for eight different SNP. Some of these are like a tight normal distribution showing that the expansion occured quickly, e.g. L21. Others like J2 show the branching occurs over 100 generations.

## Accurate Mutation rates

By relying on asymmetries of the distribution to find singular lineages we have to be aware that the mutation process itself might not be symmetric, this is called bias (Wakeley, 1994). If ignored we might be just detecting these asymmetries. Changing the model, the probability of a mutation per generation is

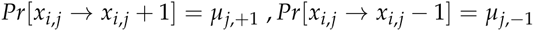

If this marker is free from singular lineages we observe that the ratio of the frequencies to the left and right of the mode is

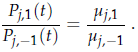

which is time independent. So using eight very large SNP projects we find enough markers free of singularities to compute these ratios and their standard deviations, see Table 9 in Appendix and SI for computations. In particular about 33% of DYS markers show asymmetric ratios are significant, i.e more than two SD from ratio 1. These asymmetric ratios play a very important role, for this ratio is all you need to detect a singular lineage and reduce it. Of course not knowing the exact asymmetric ratio means that bootstrap methods are used extensively for singular reduction, both to compute values and SD.

These methods also imply a new way of computing mutation rates. Previously, there were methods based on meiosis data or phylogenetic studies of family DNA projects (which gave quite different rates). We begin with 8 very large SNP projects from FTDNA using 37 markers, of course with unknown *TMRCA*. We first reduce singular lineages. Then taking asymmetry into account we find mutation rates are the fixed points of an iterative process. This takes about 3 iterates to converge. These mutation rates are normally distributed with mean and SD. Discarding markers with mutation *SD* > 33% leaves us with 29 markers. We find this advanced phylogenetic method gives mutation rates close to those obtained from meiosis and nearly 1/2 the values obtained from the usual phylogenetic method. Further validation comes from finding that the equivalence of our rates with meiosis implies *apriori* a human generation of c30 years.

## Results

Accuracy is verified by checking for consistency over all of European history beginning with Group A of three genealogical projects of medieval age:

**Table 1:** Medieval comparison

| Group | TMRCA | [SD] | Oldest patriarch |
| --- | --- | --- | --- |
| Hamilton | 1399AD | [180] | Fitzgilbert d1330AD |
| Macdonald | 1060AD | [270] | Somerfeld c1100 – 1150AD |
| O’Niall | 100AD | [260] | Niall 300AD/Conn100AD |

Archeological finds convinced Marija Gimbutas(1956) to attribute Proto Indo-European (PIE) to the Yamnaya Culture c3000-4000BC of the Russian Steppes. This is consistent with main-stream linguistic theory, some even wrote of linguistic DNA, see Anthony (2007). But actual genetics was ignored because current genetic clocks for R1b1a2 pointed to the Renfrew Hypothesis(1987) that PIE spread from Neolithic Anatolia, c 6000BC. Or Mesolithic or Paleolithic, depending on the genetic clock. However no-one checked if their clock worked over the whole range of time for different lineages.

The next table shows the expansion times of the dominant European Y-haplotypes R1b1a2 & R1a1a. This data is from FTDNA projects for region X only using individuals with named ancestor from X. These independent results agree within the standard deviation, with dates matching the Corded Ware Culture, a seminomadic people with wagons and horses who expanded west from the Urkraine c3500BC. This is consistent with the oldest R1b1a2, R1a1a skeletons being from the Yamnaya Culture, c 3300BC, see R. Nielsen et al (2015) S. Pääbo et al (2015).

**Table 2:**
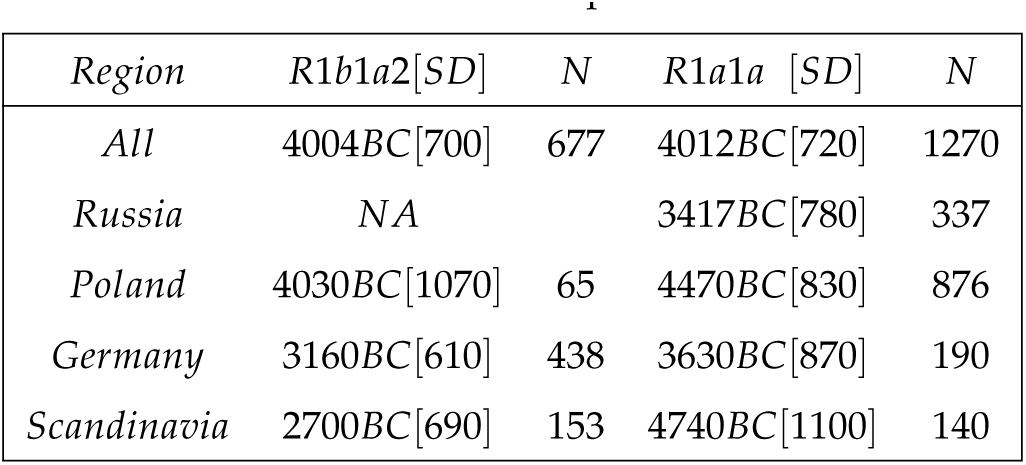
Indo-European SNP

The regional differences seem random suggesting rapid moment of whole peoples. Averaging overall (weighted by 1/Variance as the best linear estimate) gives R1b1a2 has TMRCA 4011BC(SD 580), R1a1a has TMRCA 3980BC(SD 380).

An interesting intermediate step occurs between the medieval and eneolithic. The mythical Irish Chronicles relate that the O’Niall descend directly from the first Gaelic High Kings, which tradition dated c1500 BC. The O’Niall have the unique mutation M222 which is a branch of the haplotype DF13 which is turn a branch of L21 and that of P312. The most likely TMRCA are

**Table 3:** Pre-Celtic SNP

| SNP | TMRCA | SD |
| --- | --- | --- |
| P312 | 2728BC | 376 |
| L21 | 2216BC | 294 |
| DF13 | 1867BC | 328 |

These are dates for for early to late Bell Beaker Culture of Western Europe, 2800-1800BC. Indeed the only known Bell Beaker genome was found to be P312 with ^14^*C* date 2300BC.

**Table 4:** Non FTDNA data

| *data* | *Haplotype* | *N* | *TMRCA* | *[SD]* |
| --- | --- | --- | --- | --- |
| *Underhill* | *R1a1a1* | 974 | 3000BC | [800] |
| *Rootsi* | *G2a* | 536 | 21000BC | [4200] |

Our method requires large data sets and many markers which means we have to rely on data from FTDNA, finding 29 useable markers out of standard 37 they use. In fact many researchers have used FTDNA data, e.g Balaresque at al(2010). We think our method of reduction with robust statistics solves any problems with this data. To test this we used our method with R1a1a1 data obtained from Underhill(2015) with *N* = 974 and 15 useable markers, and G2a from Roostsi(2012) with a different 15 markers, see Table 4. With 29 markers the TMRCA for R1a1a1 is somewhat after our date for R1a1a, while G2a predates ours for G2a2(Table 5 are Western European only, see Table 7 for Anatolian).

**Table 5 :** TMRCA ybp for 29, 15, 7 markers

| SNP | N | T29 | SD | T15 | SD | T7 | SD |
| --- | --- | --- | --- | --- | --- | --- | --- |
| G2a2 | 1221 | 7654 | 1007 | 10800 | 2340 | 6580 | 2021 |
| R1b1a2 | 660 | 6004 | 700 | 6800 | 1075 | 8264 | 2217 |
| R1a1a | 1270 | 6012 | 720 | 5270 | 840 | 5643 | 1665 |
| I1 | 2898 | 4109 | 460 | 5114 | 1010 | 5799 | 1775 |
| L21 | 1029 | 3927 | 415 | 4052 | 503 | 4252 | 882 |
| U106 | 1533 | 4883 | 534 | 3043 | 710 | 4243 | 947 |
| J2 | 1241 | 14490 | 2000 | 21060 | 3860 | 8814 | 2302 |
| P312 | 971 | 4449 | 490 | 5340 | 748 | 5145 | 1128 |
| Mean SD |  |  | 12 % |  | 18 % |  | 26 % |

Table 5 shows the results of extensive simulations using random subsets of our FTDNA data, for 29, 15(using Underhill markers) and 7 markers (Balaresque markers). In a fact our data for R1a1a for 15 markers is rather close to the Underhill data for R1a1a1.

However once you get down to 7 markers the confidence interval becomes too large for accuracy. Also it becomes difficult to deal with outliers. We might expect 5% accuracy with 120 markers.

An example with few markers is the R1b1a2 data of Balaresque et al(2010). Our method (this time with 7 useable markers) gave SD ~ 30%. Now Balaresque used BATWING to suggest a Neolithic origin in Anatolia. With the same Cinnioglu(2004) data our method gives for Turkish R1b1a2 anything from the Ice Age to the Iron Age as seen in

**Table 6:**
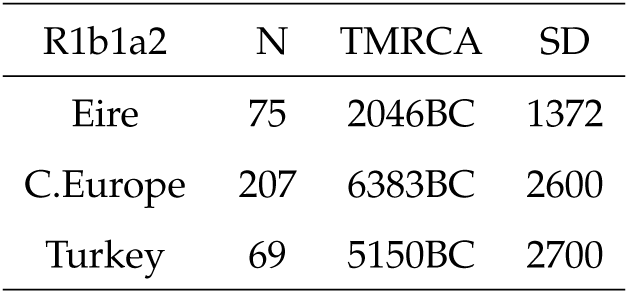
Balaresque Data

| R1b1a2 | N | TMRCA | SD |
| --- | --- | --- | --- |
| Eire | 75 | 2046BC | 1372 |
| C.Europe | 207 | 6383BC | 2600 |
| Turkey | 69 | 5150BC | 2700 |

Fortunately, once again, we find good data from FTDNA: the Armenian DNA project, see Table 7. By tradition the Armenians entered Anatolia from the Balkans c1000BC so they might not seem a good example of ancient Anatolian DNA. But some 100 generations of genetic diffusion has resulted in an Armenian distribution of Haplotypes J2, G2a2b, R1b1a2 closely matching that of all Anatolians, therefore representive of typical Anatolian DNA. We see that Anatolian R1b1a2 arrived after c3800BC, ruling out the Neolithic expansion c6000BC. When dealing with regional haplotypes, e.g. R1b1a2 in Anatolia, the *TMRCA* is only a upper bound for the arrival times, for the genetic spread may be carried by movements of whole peoples from some other region. This means one has to be careful interpreting regional data, e.g. the TMRCA for the R1b1a2(USA) is c4000BC but nobody thinks it arrived then.

**Table 7:** Anatolian SNP

| *Armenian* | *N* | *TMRCA* | *[SD]* |
| --- | --- | --- | --- |
| *R1b1a2* | 99 | 3780BC | [1000] |
| *G2a2b* | 46 | 10000BC | [2500] |
| *J2* | 97 | 12600BC | [2500] |

Observe that our *TMRCA* for Armenian G2a2b (formerly G2a3) and J2 correlates with the first Neolithic farmers in the Middle East. From Table 5 we see J2, G2a2b for all of Western Europe (non-Armenian data). Our dates show J2 was expanding at the end of the Ice Age. Modern J2 is still concentrated in the fertile crescent, but also in disconnected regions across the Mediterranean. The old genetic model predicted a continuous wave of Neolithic farmers settling Europe (L.L. Cavalli-Sforza et al, 2004). But you cannot have a continuous maritime settlement: it must be *leap-frog*. Also repeated resettlement from the Eastern Mediterranean has mixed ancient J2 populations, and our method gives the oldest date. On the other hand G2a2b shows exactly the dates expected from a continuous wave of Neolithic farmers across Central Europe. Our dates are consistent with recent findings that the majority of early Neolithic skeletons found in Western Europe are G2a2, c 5000BC, whereas the oldest R1b1a2 found in Western Europe is Bellbeaker c2300BC.

The problem of the peopling of the Americas has received as much attention. Y-chromosome analysis of (subarctic) Amerindians shows haplotype Q1 is dominant. Bortolini et al(2003) applied KAPZ with 8 markers to a sample of *N* = 24 to give a TMRCA for Q1 of c11600BC, (3000SD). More recent studies, e.g Reich et al(2012), have been careful not to estimate a TMRCA. We use 22 marker data from Battaglia et al(2013) who like Riech saw only one Q1 migration, but gave no dates. Ours are

**Table 8:** Battaglia Amerindian data

| SNP | N | TMRCA | SD |
| --- | --- | --- | --- |
| all Q1 | 437 | 12600BC | 2000 |
| Q1-L54 | 73 | 13300BC | 2800 |
| Q1-M3 | 335 | 10500BC | 1770 |

This is supported by 29 marker FTDNA data from their New Mexico project:

**Table 9:** New Mexico FTDNA

| SNP | N | TMRCA | SD |
| --- | --- | --- | --- |
| NM Q1 | 110 | 16200BC | 3000 |
| NM M3 | 71 | 11030BC | 2300 |

Thus we are 90% certain that the entry of Q1 into the Americas occurred before the Canadian Ice-Corridor was passable, c9500BC, see Mandryk, C.A.S. et al (2001). The Q1-M3 subgroup is found through most of the Americas, in fact is a majority of Amerindian Q1. Its later date is very interesting as it is about the time the Ice-Corridor opens. While it is probable that the subbranch L54 arrived with Q1 we are 90% certain that M3 did not arrive with the original migration. (We got 6000BC for the haplotype C3 which probably arrived with the later Na-Dene migration).

## Discussion

Archeology, evolutionary biology, not to mention epidemiology, forensics and genealogy are just some of the applications of molecular clocks. Unfortunately current clocks have been found to give only “ballpark” estimates. Our method gives accurate time, at least for the human y-chromosome verified over the period 500 15, 000*ybp*. Our main ideas hold for general molecular clocks, however our Y-clock has features specific to STR mutations. It would be important to develop reduction of singular lineages for other clocks.

Some geneticists thought natural selection makes mutation rates too variable to be useful. The problem is confusion between the actual biochemistry giving mutations and superimposed processes like kin selection producing apparently greater rates. Notice that the SD for our mutation rates is about 14%, previously many authors claimed about 10% yet had large differences with other studies.

Many applications to genetics, forensics, genealogy require the TMRCA between just two individuals, or between two species, a classic method was given by Walsh (2001). While we are accurate for “big data", for this “two-body problem” one cannot determine what singular lineages the branching has been through. Just using our new asymmetric mutation rates will not work. So it would be important to find an accurate method.

Pääbo et al(2015), Nielsen et al (2015) observed all 6 skeletons from Yamnaya sites, c 3300BC by ^14^*C* dating, are either R1a1b2 and R1a1a. This involves very difficult genetic analysis of fossils which may not always be available. Also such analysis cannot date the origin of R1a1b2 and R1a1a. Our *TMRCA* shows both these haplotypes expanding at essentially the same time c4000BC. This and our later date for Anatolia, combined with Pääbo et al, implies that R1b1a2 and R1a1a must have originated in the Yamnaya Culture.

In checking accuracy we ran into the question of the origins of Indo-European. Although there are genes for language there is certainly none specific to Indo-European language. Thus inferences have to be indirect. Marija Gimbutas(1956) saw patterns in symbolism and burial rituals suggesting the Yamnaya Culture was the cradle of Indo-European. Also their physiology was robustly Europeanoid unlike the gracile skeletons of Neolithic Europe, but this could be nutrition and not genetic. From the above we conclude that the spread of this robust type into Western Europe in the late Neolithic marked an influx of Steppe nomads. Now if R1b1a2 had been shown to spread from Anatolia c6000BC it would have been taken as strong evidence for “out of Anatolia” because of the association of R1b1a2, R1a1 with Indo-European languages. But our accuracy check showed that it was G2, J2 that spread with the Neolithic Expansion from Anatolia. Now these have been associated with Caucasian languages or Semitic, but never with Indo-European.

## Methods

This work is biomathematical theory validified by data from published sources. We give full mathematical details in the following appendix. Also see Supplementary Information (SI) for detailed algorithms and MATHEMATICA worksheets, examples and some data. To verify the theory and compute mutation rates we use diverse data, from FTDNA y-haplotype projects for G2a2b, R1b1a2, R1a, I1, L21, U106, J2, P312. Also we used regional projects for Germany, Scandinavia, Poland and Russia for their R1b1a2, R1a1a data. The Armenian DNA project was important for its R1b1a2, J2 and G2a2b data. We also used DNA projects M222 (O’Niall), Macdonald (Group A which is R1a1a), Hamilton (group A which is I1). This was compared with non FTDNA data from Balaresque, Underhill, Rootsi and Battaglia et al.

## Appendix

### Biomathematical theory

Our method requires a return to basic principles.

### Fundamental Solutions

The Y-chromosome has DYS marked by *j* = 1, …*n*, where one can count the STR number *x*_*j*_. Consider the probability *Pj*,*k* (at time *t* generations) that at marker *j* we have *x*_*j*_ = *k*. This satisfies the homogenous stochastic system

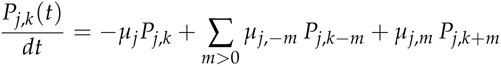

This homogenous system gives a uniform expansion from a single patriarch.

The system is essentially the model of Wehrhahn(1975) who had *μ*_*j*, −1_ = *μ*_*j*,1_. We introduce asymmetric mutations with total rate

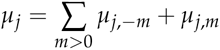

About 50% of DYS markers show asymmetric mutations, i.e. *μ*_*j*, −1_ ≠ *μ*_*j*,1_.

We use the generator function with complex variable *z*

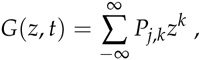

(see Durrett, 202), and normalized initial condition *x*_*j*_ = 0 or *P*_*j*,0_(0) = 1:

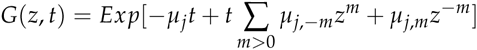

Then *G* can be expanded in powers of *z* to give *P*_*j*,*k*_ (*t*). Now for the simplest asymmetric case, with only one step mutations, we have 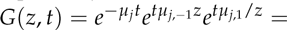

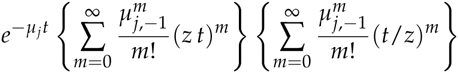

so using the Hyperbolic Bessel Function of Order *k* ≥ 0, see Olver (1964)

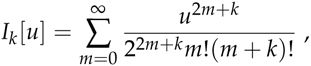

we see that the homogenous system has fundamental solution

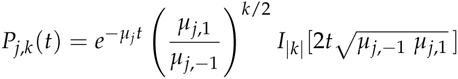

From the fundamental solution we find, independently of time

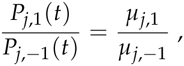

which we call the *asymmetric ratio*. It will be repeatedly used.

Of course the actual initial value is not *x*_*j*_ = 0 but was usually taken to be the mode *m*_*j*_ which was assumed to be the value for original patriarch. From the present distribution of data we use the frequency

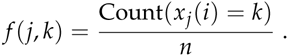

One problem with the KAPZ formula is that higher frequencies *f* (*j*, *k*), *k* = 2, 3… are overrepresented in the actual data. This is because the probability of a spontaneous two step mutation is much higher then the product of two one step mutations. So instead we use the frequency to solve the transcendental equation for the unknown *t*

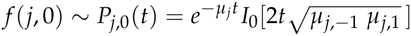

This nonlinear equation is easily solved via mathematical software such as MATHEMATICA (I used version 9 running on a boosted 2014 iMac which has accurate hyperbolic Bessel functions. Earlier versions on older iMacs gave inaccuracies so one had to compile one’s own functions). Using this formula resolves some other problems with the KAPZ method even for a uniform flow, e.g. *μ*_*j*, − 1_ ≠ *μ*_*j*,1_ gives an extra quadratic term which if ignored causes large errors.

### Heterogeneous diffusion equation

However the main problem is singularities in the stochastic process. For a uniform stochastic process, 1 − *P*_*j*,0_(*t*) ~ 1 − *f* (*j*, 0) is the probability of some mutation. So the expected variance is *f* (*j*, 0)(1 − *f* (*j*, 0)). Thus if the actual data variance *Vj* > > *f* (*j*, 0)(1 − *f* (*j*, 0)) we are not uniform. Now a sublineage of very high fertility increases variance, giving apparently greater *TMRCA* although it is unchanged. One finds similar results for Bayesian methods.

The correct approach to nonuniformity assumes at times *ti* (generations ago) a certain proportion 0 ≤ *ρi* ≤ 1 of the present population originated from a “virtual patriarch” with an initial STR value *mi*. The resulting system :

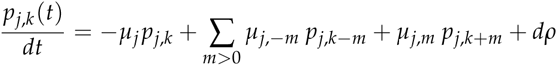

i.e. *dρ* are atoms of weight *ρi* with STR value *m*_*i*_ occurring at time *t*_*i*_. As the system is linear and isotropic the solution is a combination of fundamental solutions *P* of the homogenous system. Thus the present distribution *f* (*j*, *k*) is

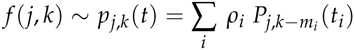

This allows us to consider populations mixed by having singular lineages from overfertile patriarchs, or by actual immigration from the outside. For example Nielsen, R. and J. Wakeley (2001) considered mixed populations with two sources. However in our general case has no obvious limit on the number of sources. The inverse problem seeks to find singularities from present data. Unfortunately, in general, inversion is ill posed for such systems like the heat equation. This instability produces poor accuracy. Furthermore there is no unique solution, e.g.the present distribution could have been created yesterday.

However we find that ~ 50% of the DYS markers show no significant difference from the uniform expansion of a single patriarch, i.e. the data variance *V*_*j*_ is close to the expected variance *f* (*j*, 0)(1 − *f* (*j*, 0)). The other markers show at most one significant side branch, i.e. there is an original branch starting at time *t*_*j*,0_ with STR *m*_0_ and a second one with STR *m*_1_ = *m*_0_ ± 1 at time *t*_*j*,1_ < *t*_*j*,0_ with significant 0 < *ρ*_1_ < *ρ*_0_. Yhus we have mane able 1-2 sources per marker.

### Reduction of Singular Lineages

We locate these singular lineages by looking for asymmetries in the distribution. For a uniform flow from a single patriarch the frequency of STR value *k* is given by *f* (*j*, *k*) ~ *P*_*j,k*_ (*t*). The asymmetric ratio:

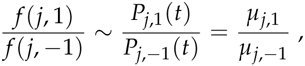

is completely independent of time *t*. Therefore if say

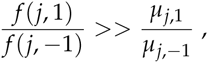

we have a singular lineage at *k* = +1. Thus the excess at *k* = +1 is

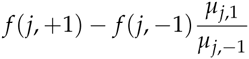

To first order approximation then frequency *f* (*j*, +2) is due to this singularity at *j* = +1 which therefore gave a contribution

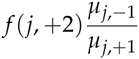

to *k* = 0. Thus removing the effect of the singularity at *k* = +1 leads to new frequencies

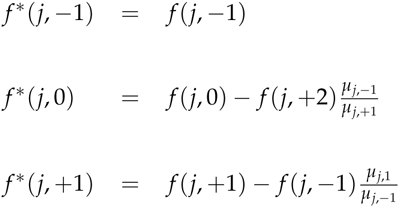

These of course are no longer normalized so we rescale to obtain the renormalized frequency *F*(*j*, *k*), e.g.

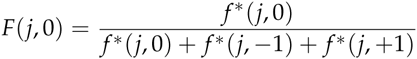

which will be used to compute the expansion time for marker *j*. There are similar formulae if the singularity was at *k* = −1.

However there is sampling error both in the frequencies and the *μ*_*j*,1_, *μ*_*j*, −1_. So we bootstrap taking into account these uncertainties, running the computation thousands of times. Generally we find the branch singularity is always one of *k* = 0, +1, −1 with no SD. In a few cases the singularity may seem to wander between *k* = 0, +1, −1. So in the case of a wandering singularity we obtain a distribution over *k* = 0, +1, −1 with a mean and SD. In these cases we find the singularity is relatively small and does not make much difference to the final result. However to have a stable method we do not throw out these wandering singularities but in the algorithm use the mean to average between *k* = 0 and *k* = ±1, e.g. if the mean is *k* = 0 then we use the original unreduced frequency.

Notice that we assume at most one side branch. In theory there could be many and solving for these produce even better approximations to the present data. In fact you could get perfect matching but find the atoms were created yesterday! The thing is that while many markers show significant deviation from a uniform flow from a single patriarch, after we have carried out reduction for one possible side branch we find no significant difference from a uniform flow, i.e. the difference is within the SD. This is of course an approximation, the next level beyond Zuckerkandl and Pauling, but given the noise in the data perhaps the best we can do. Later we further reduce the effect of outliers by using robust statistics.

Reducing the singular lineages increases the frequency *f* (*j*, 0) of the mode and decreases the computed *TMRCA*. But as the method of reducing singularities does not respect higher frequencies *f* (*j*, *k*) it follows the KAPZ formula cannot be used and instead we use the probability of no mutations, i.e. solve

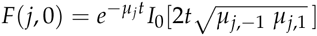

This is done for each DYS marker *j*, giving expansion times *t*_1_, …*t*_*N*_ for each marker, with computed CI. (An extra fixed source of error is the uncertainty in the mutation rates which we deal with later). We find the reduction of singularities makes striking difference to the *tj* of the effected markers, often a reduction of *~* 50% for *TMRCA*.

Now the existence of side branches implies that the main branch could itself have been the side branch for an earlier branch that did not survive. Thus we do not expect the expansion times *t*_1_, …*t*_*n*_ for each marker to be essentially equal., i.e they are not within the SD of each other. Indeed we see that the distribution of the times *tj* for different markers are almost certainly not randomly arranged about a single *TRMCA T* but distributed from *T* to the present. This is seen whether you use reduction or not, or our mutation rates or not. (For a given population one could scale mutation rates to get equal *t*_*j*_, but then applying these adhoc mutation rates to other populations does not yield the same values). The spread out distribution of surviving branches is another verification of our theory of many extinctions, few survivors. The distribution of the times *t*_*j*_ for different markers we call the branching distribution, which is now discussed.

### The Branching Distribution

The times *t*_j_ for different markers are sorted from the youngest to the oldest, forming a sequence 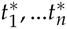. The generation of these branches is by an unknown probability distribution *dτ*0 over [0, *T*]. We model *dτ*0 by assuming a surviving lineage is generated at random with probability *β*Δ*t* in time period [*t*, *t* + Δ*t*], multiplied by the probability that the branching hasn’t already occurred. The constant *β* averages fertility and extinction rates, the chance of a new lineage surviving. As *β* → ∞ we get current theory where all lineages originate from a single patriarch at time *T*. Simulations with the data show that *β* varies in the range 1 to ∞. We make no a priori estimate of *β*, unlike Bayesian methods where an overall fertility rate is a predetermined parameter. Instead our stochastic simulation will find the most likely *β*, *T* in each case. Assuming independence, then the generation of branches follows the well known exponential distribution:

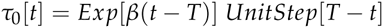

Notice this implies a finite probability that some markers have essentially zero mutations. This is actually seen in examples. Both the Hamilton Gp A and Macdonald Gp A have number of individuals *N* > 100. For the time scale of > 700 years we do not expect there is more than one marker out of 33 which shows absolutely no mutations from the mode. In fact in both cases there are 8 markers where all *N* individuals have exactly the same STR value.

We also tested this theory in simulations where we obtained random branching times based on the density for various values of *T*, *β*. In SI 2 we show results for “pseudo R1b1a2", i.e with TMRCA *T* = 200 generations. This looks like the actual spectrum for R1b1a2 for 1/*β ~* 20 generations, i.e. a lop sided bell curve showing fast expansion. For 1/*β ~* 100 the curve looks like our curve for J2, i.e spread out indicating much branching.

Estimating the parameter *T* for an exponential distribution is a well known problem of statistics. Kendall proved the best estimate for *T* would be max *t*_*j*_. Unfortunately there is also considerable error *λ*_*j*_% for the mutation rates *μ*_*j*_. Later we give a method for reducing this error, even so we find the SD in the range 10% 30% which gives corresponding range in error for each *t*_*j*_. We understand that the *t*_*j*_ are being generated by the distribution *dτ*0 but superimposed on this is a further uncertainty due to mutation rates etc. In particular the largest *t*_*j*_ may be wildly inaccurate. Also we found that simply taking the average consistently underestimates the *TMRCA* by a wide margin.

Assuming the mutation rates have normal distribution with mean *μ*_*j*_ and variance 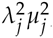, the *t*_*j*_ have SD *t*_*j*_ *λ*_*j*_. Thus the actual data for 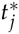 has probability density function for *s* > 0

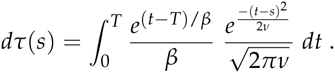

The variance *v* depends on two sources. First from the uncertainty in mutation rates, for each of the *n* markers we get variance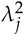, giving total

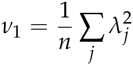

However a small sample also has inherent error from sampling. We are measuring the probability that there is a mutation. This is binomial with probability *H*_*j*_ (*t*) given by

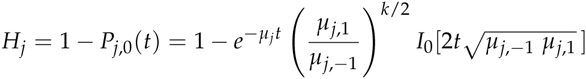

Hence for sample size *N* there is variance *H*_*j*_ (1 − *H*_*j*_)/*N*, so the variance in time due to this is scaled by the derivative giving:

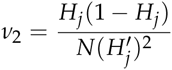

The function 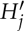 has actually to be computed as an inverse function depending on *H*_*j*_. Therefore the total variance averaged over all *n* markers is *v* = *v*_1_ + *v*_2_. Although for large samples (*N* > 1000) the second term is insignificant it does effect the results once you get to *N* = 100. In our algorithm the branching distribution is used to generate large numbers of random branching times so as to bootstrap error estimates. In turns out much faster to compile the distribution function as a table which can be repeatedly called on.

### Estimating TMRCA by Robust Statistics

Inaccurate large values of 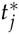 are mitigated by using “robust” statistics with quintiles instead of means/variances. Using FTDNA data we began with 37 markers. However the 4 markers of DYS464 are unordered and cannot be used. Also we find that markers DYS 19/394, 385b, 459b, CDYb have errors > 33% in mutation rates so are not used. (These are some of the most popular ones in the literature!). We find if we just use the 15 markers with errors < 15% the final SD is about 13% compared with 12% using 29. So usually we have *n* = 29 markers and take “quintiles” 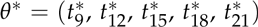. This means that tail end data is not discarded but kept as the information there are 8 values of 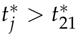, which effectively deals with outliers. Bootstrap methods give the confidence interval CI for each quintile.

Thus we wish to find the best estimate of *T* given *θ\** (and CI). This well known statistical problem was investigated by Stochastic Simulations (SS). We also tried Maximum Likehood Methods which gave similar results but with larger CI. Monte-Carlo Methods are used to produce very large numbers (~ 10^7^) of *T*, *β* with corresponding Distribution. These randomly generate ordered times (*s*_1_ …*s*_29_) for which we take the quintiles *θ* = (*s*_9_, *s*_12_, *s*_15_, *s*_18_, *s*_21_). We filter by requiring that *θ* close to the data *θ\**, i.e. ||*θ* − θ|| < rSD*. We tried *r* = 3, 2, 1, 0.5 obtaining almost identical final results, subsequently use *r* = 2. This gives a stochastic neighborhood of *θ\** typically containing *>* 10^5^ sets of data but with *T* is known for each *θ* ∊ 𝒰 Thus we can construct a quasilinear estimator:

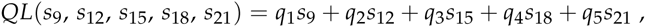

and use least squares over *𝒰* to find constants (*q*_1_, *q*_2_, *q*_3_, *q*_4_, *q*_5_) minimizing

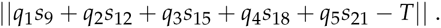

The (*q*_1_, *q*_2_, *q*_3_, *q*_4_, *q*_5_) are computed in MATHEMATICA We test this by applying the QL to all of *𝒰*, unsurprisingly

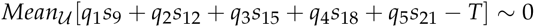

What is important is that we find the uncertainty in the SS itself. Actually this depends on the data and is calculated in each case but for our examples we find

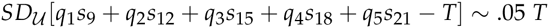

Finally the quasilinear estimator is applied to the experimental data

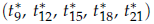

to obtain our best estimate of *T*. Application of *QL* computes the SD for our data, giving part of the overall SD. This must be combined with the SD coming from the uncertainty in the SS. Overall we find that our method has SD ~ 12%, this includes variances from our data, mutation rates and uncertainty in the SS. Also we tested with 7, 13 “quintiles” instead of 5 obtaining almost identical TMRCA and a slight improvement in accuracy to < 11%. We also tested with 15 and 7 markers. Here one must use “quintiles” 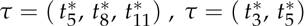, respectively with all the loss of accuracy that implies. See Table 5 for comparisons using 29, 15, 7 markers on same data.

### Accurate Mutation rates

Any genetic clock depends on reasonably accurate mutation rates. The meiosis method looks for mutations in father-son studies. However typical rates of *μ* = .002 would require nearly 50,000 pairs to get an SD of 10%. The phylogenetic approach studies well developed DNA/genealogy data. So inverting the KAPZ formula would yield accurate rates. However, *singular lineages* makes this problematic.

To compute our rates we apply our theory to the large DNA projects for the SNP M222, L21, P312, U106, R1b1a2, I1, R1a1a. This avoids dealing with populations such as family DNA projects which are self selecting, i.e only those with the correct surname which neglects distant branches. Also we have very large samples, our average *N* > 1000. Greater accuracy should come from more generations and individuals. The problem is that we do not know their *TMRCA*.

### Asymmetric Mutation

However before computing mutation rates we must consider asymmetric mutations, i.e. the left and right mutation rates *μ*_*j*, −1_ ≠ *μ*_*j*,1_. For a uniform stochastic process we again use the asymmetric ratio

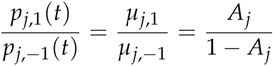

to define the *asymmetric constant A_j_ ∈* [0, 1] for marker *j*. For example *Aj* = 0.5 is complete symmetry. Of course singularities will effect this ratio, however these only occur < 50% of markers. Thus for each marker, SNP we compute this ratio. We find the SD for each SNP is relatively small while the difference between SNP can be large. However for each marker, using 8 SNP enables outliers to be easily removed leaving allowing us to use simple linear regression: i.e. average of the *A*_*j*_ over the remaining SNP groups. We see that asymmetry is a real effect: 50% of the *A*_*j*_ are more than two SD from symmetry *A*_*j*_ = 0.5.

Observe this is significant. The total second moment is

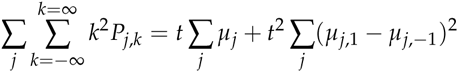

So using all our 33 DYS markers with our *μ*_*j*_, we compute constants

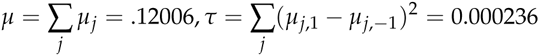

The classical KAPZ formula gives variance *V* = *μt* compared to the corrected formula *μt* + *τt*^2^. The uncorrected KAPZ gives an overestimate > 400% for > 200 generations. This effect can be nullified by using the mean instead of the mode, variance instead of the second moment, however failing to do so gives a large error. Furthermore other methods which assume symmetric mutations will also be inaccurate. Having estimates on the asymmetry is essential to our method because we find singular lineages by looking for asymmetry in the data. Any such anomaly needs to be significantly greater than the natural asymmetry.

### Mutation Rates as a fixed Point

Next we compute mutation rates using 8 very large SNP groups. First, using the asymmetric constants we find singular lineages and reduce their effect. We take account of the error in the *A*_*j*_ by a bootstrap technique, which gives the variance for each frequency *f* (*j*, 0).

For each SNP *k* if DYS markers *j* started their mutations at the same time TMRCA *Tk* we could calculate mutation rates *μ*_*j*_ via

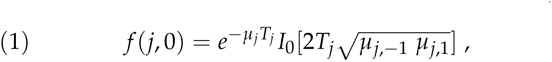

or rather average the 8 different *μ*_*j*_ we would obtain. However because of branching caused by extinction of lineages the different markers do not originate at the same time but at different times *t*_*j*_. In any case we don’t know the TMRCA *T*_*k*_.

Instead we use the log modal, i.e. defining 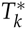 to be the log mean over the middle set of *t*_*j*_. So for a fixed marker *j* the normalized times 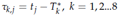 should straddle 0, i.e have a 50% chance of being positive. However the wrong choose of *μ*_*j*_ would give a bias, e.g. if *μ*_*j*_ is too big then the *t*_*j*_ are too small, so the *τ*_*k*,*j*_ will tend to be negative. In fact this is what we see if the mutation rates *μ*_*j*_ = .002 were chosen. In the SI figure shows the *τk*,*j*, *k* = 1, ¨8 bunched around a nonzero point. Thus we try to find *μ*_*j*_ so that the *τk*,*j*, *k* = 1, 2, ¨8 straddle zero. However the *τ*_*k*,*j*_, *k* = 1, 2, ..8 depend nonlinearly on the rates *μ*_*j*_, as does the log modal 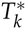, *k* = 1,..8. This nonlinear regression problem is solved by an iterative scheme which starts with any reasonable set of mutation rates and corrects them. We find any reasonable first choice iterates to the same final answer. So choose *μ*_*j*_ = .002 to begin. Suppose at some stage we have apparent mutation rates *μ*_*j*_. Then, for each SNP, and each marker we solve equation (1) to obtain the apparent *t*_*j*_. For each SNP *k* = 1, ..8 we compute the mean log modal 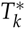. At the next we get new rates 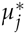 by solving the transcendental equation

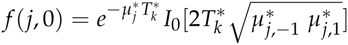

Averaging 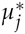, *k* = 1, ..8 we get our next set of *μ*_*j*_ of mutation rates. This method would be effected by a marker showing a singular lineage. Fortunately these are few in number and by comparison between the different SNP we remove the outliers. We then repeat the process, computing 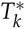again with the new rates, and another set of mutation rates.

One problem is that the iterates could tend to decrease to zero or increase to ∞, as we are only calculating relative rates. To prevent this we renormalize after each iteration so the total ∑ *μ*_*j*_ is constant. We found the iterative scheme converges in just 3 − 4 steps to a fixed set of mutation rates, unique up to a constant factor. The CI is computed by bootstrap parametrized by the uncertainties in data and the asymmetric constants.

### The generation factor γ

This method does not give absolute mutation rates but *relative* mutation rates *μ*_*j*_ *γ*, where *γ* is universal time scale constant. To find *γ* we apply our method to compute the *T* = *TMRCA* of three famous DNA projects and choose *γ* so the scaled *T*/*γ* best fits the historical record. We choose the DNA projects for the O’Niall(M222), Gp A of Macdonald (R1a1a) and Gp A of the Hamiltons (I1). These are large groups with characteristic DNA and fairly accurate times of origin. Of course finding one constant *γ* from three projects is inherently more accurate than using one project to find 33 different mutation rates. Actually assuming a generation of 30*years* these three projects yield *γ* = 1 with about 5% error, i.e. there is no actual need for this correction. This is a constant error (like uncalibrated ^14^*C* dating).

Thus *γ* is related to the length of a generation. Early researchers use 25*yrs* for *t >* 500*ybp* and 27*yrs* for *t* < 500*ybp*. Balaresque and al (2010) used 30*yrs* based on Fenner(2005) who sees a 30*yr* generation, from extensive field work with modern hunter-gatherers. Our theory allows any nominal generation as it really doesn’t matter, being included in the *γ* factor which we compute in years not generations. However to give actual mutation rates we need an actual generation so we take 30 years. This appears in our worksheet computation. However this now makes our mutation rates very similar to actual laboratory meiosis rates, see Table 8. Conversely then, if actual laboratory meiosis rates are indeed the correct long term phylogenetic rates have we have shown it follows *a priori* that the long term generation time is 30 years, i.e giving theoretical proof of Fenner

**Table 8:**
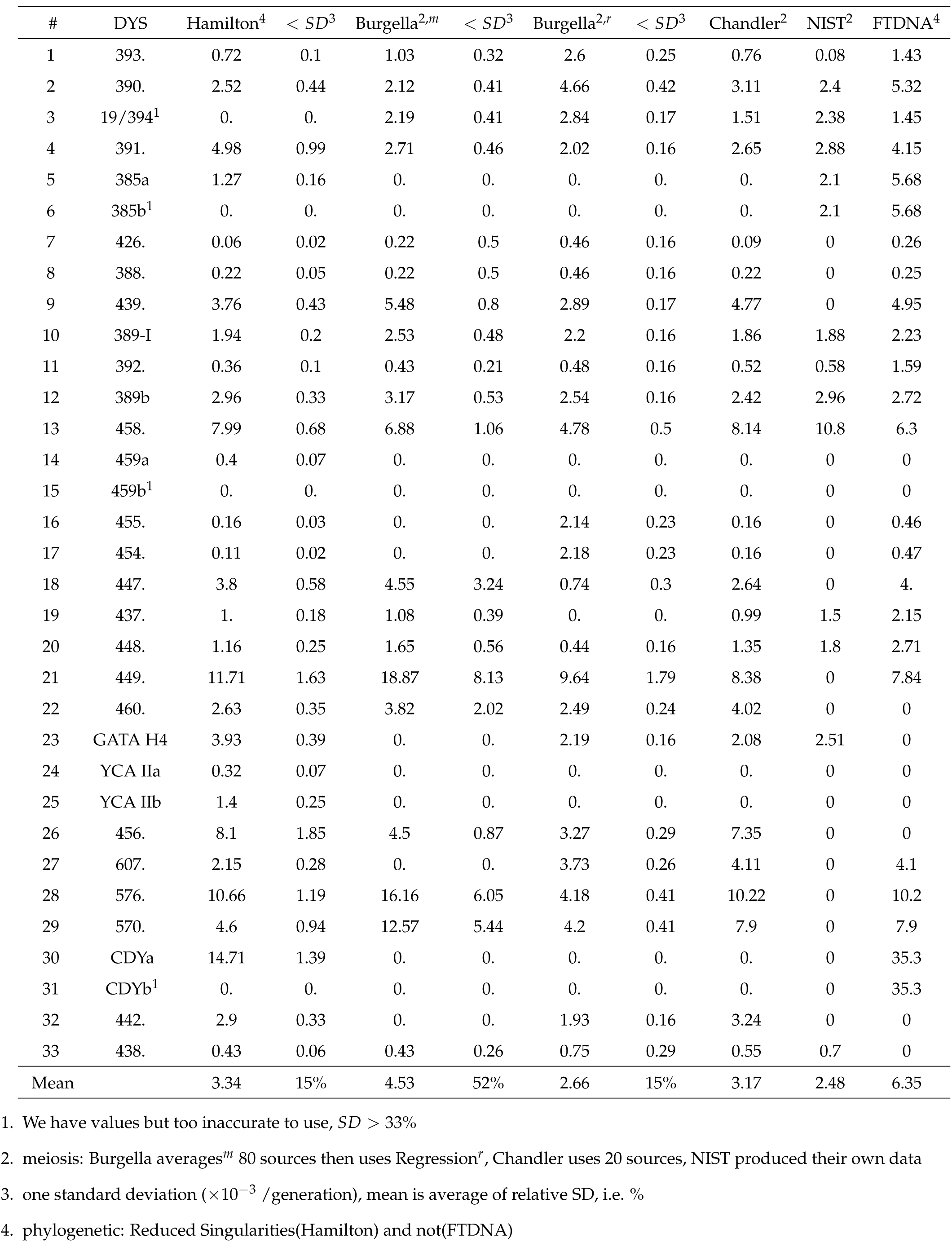
Rate of Mutation ×10^*−* 3^ /generation, at DYS: Meiosis vs Phylogenetic

**Table 9:** Asymmetric Constants

| # | DYS | A<sub>j</sub> | SD | SD to 0.5 |
| --- | --- | --- | --- | --- |
| 1. | 393 | 0.675 | 0.087 | 2. |
| 2. | 390 | 0.463 | 0.093 | 0.4 |
| 3. | 19/394 | 0.973 | 0.032 | 14.7 |
| 4. | 391 | 0.029 | 0.008 | 62.8 |
| 5. | 385a | 0.699 | 0.096 | 2.1 |
| 6. | 385b | 0.82 | 0.085 | 3.8 |
| 7. | 426 | 0.37 | 0.232 | 0.6 |
| 8. | 388 | 0.91 | 0.072 | 5.7 |
| 9. | 439 | 0.734 | 0.359 | 0.7 |
| 10. | 389-I | 0.779 | 0.105 | 2.7 |
| 11. | 392 | 0.954 | 0.04 | 11.2 |
| 12. | 389b | 0.703 | 0.325 | 0.6 |
| 13. | 458 | 0.512 | 0.137 | 0.1 |
| 14. | 459a | 0.139 | 0.125 | 2.9 |
| 15. | 459b | 0.003 | 0.001 | 353. |
| 16. | 455 | 0.277 | 0.168 | 1.3 |
| 17. | 454 | 0.962 | 0.03 | 15.4 |
| 18. | 447 | 0.154 | 0.025 | 13.6 |
| 19. | 437 | 0.09 | 0.09 | 4.6 |
| 20. | 448 | 0.216 | 0.172 | 1.6 |
| 21. | 449 | 0.518 | 0.15 | 0.1 |
| 22. | 460 | 0.107 | 0.05 | 7.9 |
| 23. | GATAH4 | 0.17 | 0.198 | 1.7 |
| 24. | YCAIIa | 0.195 | 0.163 | 1.9 |
| 25. | YCAIIb | 0.19 | 0.175 | 1.8 |
| 26. | 456 | 0.671 | 0.416 | 0.4 |
| 27. | 607 | 0.243 | 0.103 | 2.5 |
| 28. | 576 | 0.387 | 0.157 | 0.7 |
| 29. | 570 | 0.448 | 0.077 | 0.7 |
| 30. | CDYa | 0.37 | 0.181 | 0.7 |
| 31. | CDYb | 0.258 | 0.082 | 2.9 |
| 32. | 442 | 0.603 | 0.17 | 0.6 |
| 33. | 438 | 0.715 | 0.215 | 1. |
| Mean |  | 0.26<sup>1</sup> | 0.134<sup>2</sup> |  |

1. Mean difference from 0.5.
2. Showing average is 2 SD from 0.5

**Table 10:**
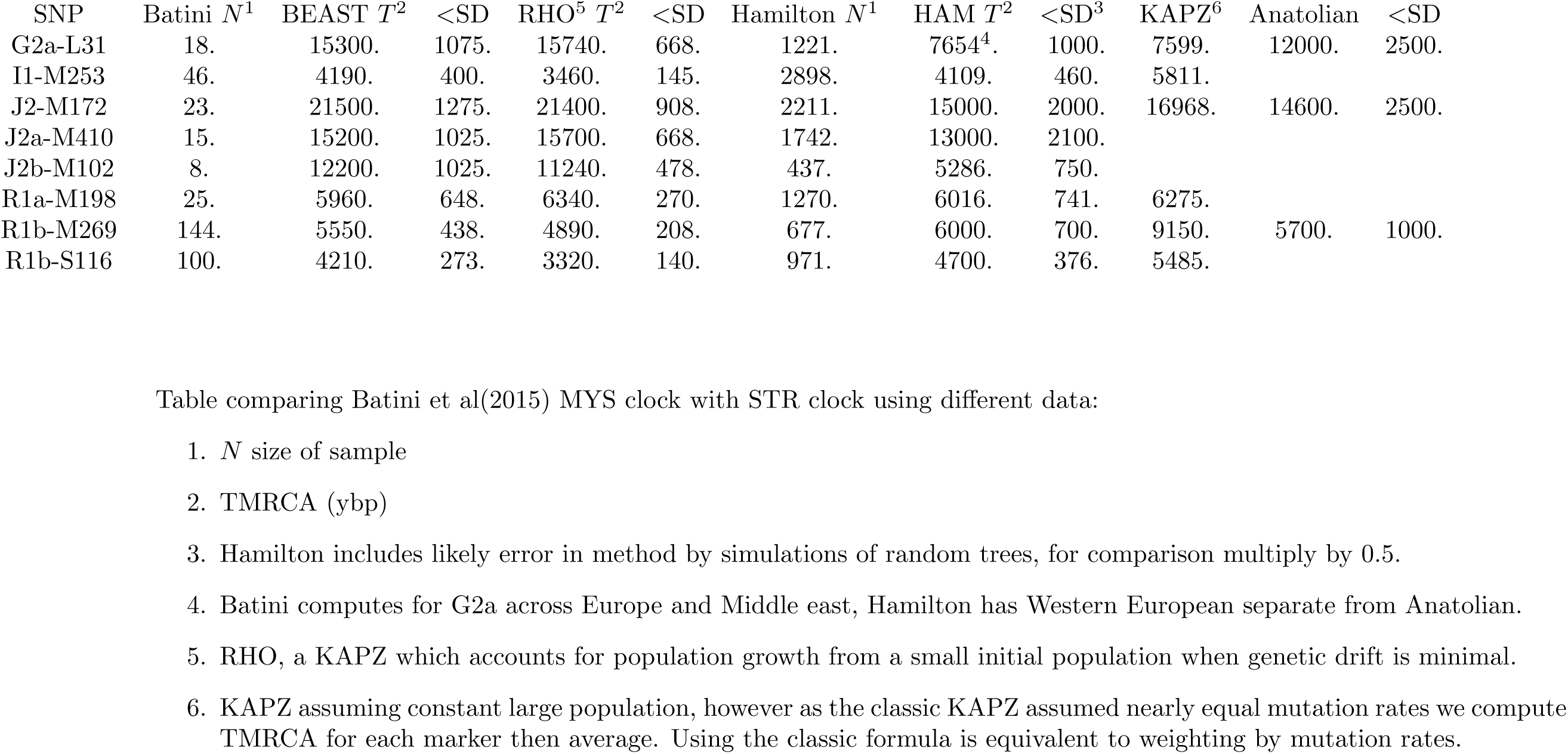
Comparison of BEAST, RHO, KAPZ with HAM, different data

| SNP | Batini $N^1$ | BEAST $T^2$ | <SD | RHO<sup>5</sup> $T^2$ | <SD | Hamilton $N^1$ | HAM $T^2$ | <SD<sup>3</sup> | KAPZ<sup>6</sup> | Anatolian | <SD |
| --- | --- | --- | --- | --- | --- | --- | --- | --- | --- | --- | --- |
| G2a-L31 | 18. | 15300. | 1075. | 15740. | 668. | 1221. | 7654<sup>4</sup>. | 1000. | 7599. | 12000. | 2500. |
| I1-M253 | 46. | 4190. | 400. | 3460. | 145. | 2898. | 4109. | 460. | 5811. |  |  |
| J2-M172 | 23. | 21500. | 1275. | 21400. | 908. | 2211. | 15000. | 2000. | 16968. | 14600. | 2500. |
| J2a-M410 | 15. | 15200. | 1025. | 15700. | 668. | 1742. | 13000. | 2100. |  |  |  |
| J2b-M102 | 8. | 12200. | 1025. | 11240. | 478. | 437. | 5286. | 750. |  |  |  |
| R1a-M198 | 25. | 5960. | 648. | 6340. | 270. | 1270. | 6016. | 741. | 6275. |  |  |
| R1b-M269 | 144. | 5550. | 438. | 4890. | 208. | 677. | 6000. | 700. | 9150. | 5700. | 1000. |
| R1b-S116 | 100. | 4210. | 273. | 3320. | 140. | 971. | 4700. | 376. | 5485. |  |  |

Table comparing Batini et al(2015) MYS clock with STR clock using different data:
1. $N$ size of sample 2. TMRCA (ybp) 3. Hamilton includes likely error in method by simulations of random trees, for comparison multiply by 0.5. 4. Batini computes for G2a across Europe and Middle east, Hamilton has Western European separate from Anatolian. 5. RHO, a KAPZ which accounts for population growth from a small initial population when genetic drift is minimal. 6. KAPZ assuming constant large population, however as the classic KAPZ assumed nearly equal mutation rates we compute TMRCA for each marker then average. Using the classic formula is equivalent to weighting by mutation rates.

## Supplementary Information (online)

1. Distribution about mode for each DYS of R1b1a2
2. Randomly generated Branching Spectrum
3. Complete Mathematica worksheet for R1b1a2
4. Complete Mathematica worksheet for computing Asymmetric constants
5. Complete Mathematica worksheet for computing Mutation rates
6. Summary of Mathematica worksheets for computing TMRCA of the SNP: G2a2 (Western Europe), R1b1a2, R1a1a, I1, L21, U106, J2, P312
7. DNA data for R1b1a2

## SI 1: distribution about mode for DYS markers j=1, ¨33 for R1b1a2

Real data (blue) shows deviation from ideal distribution(red), especially for markers

#: 1, 9, 13, 18, 21, 23, 26, 27, 28, 29, 30, 31

i.e. 33% have significant deviation. (Ideal distribution, red, showing natural asymmetry).

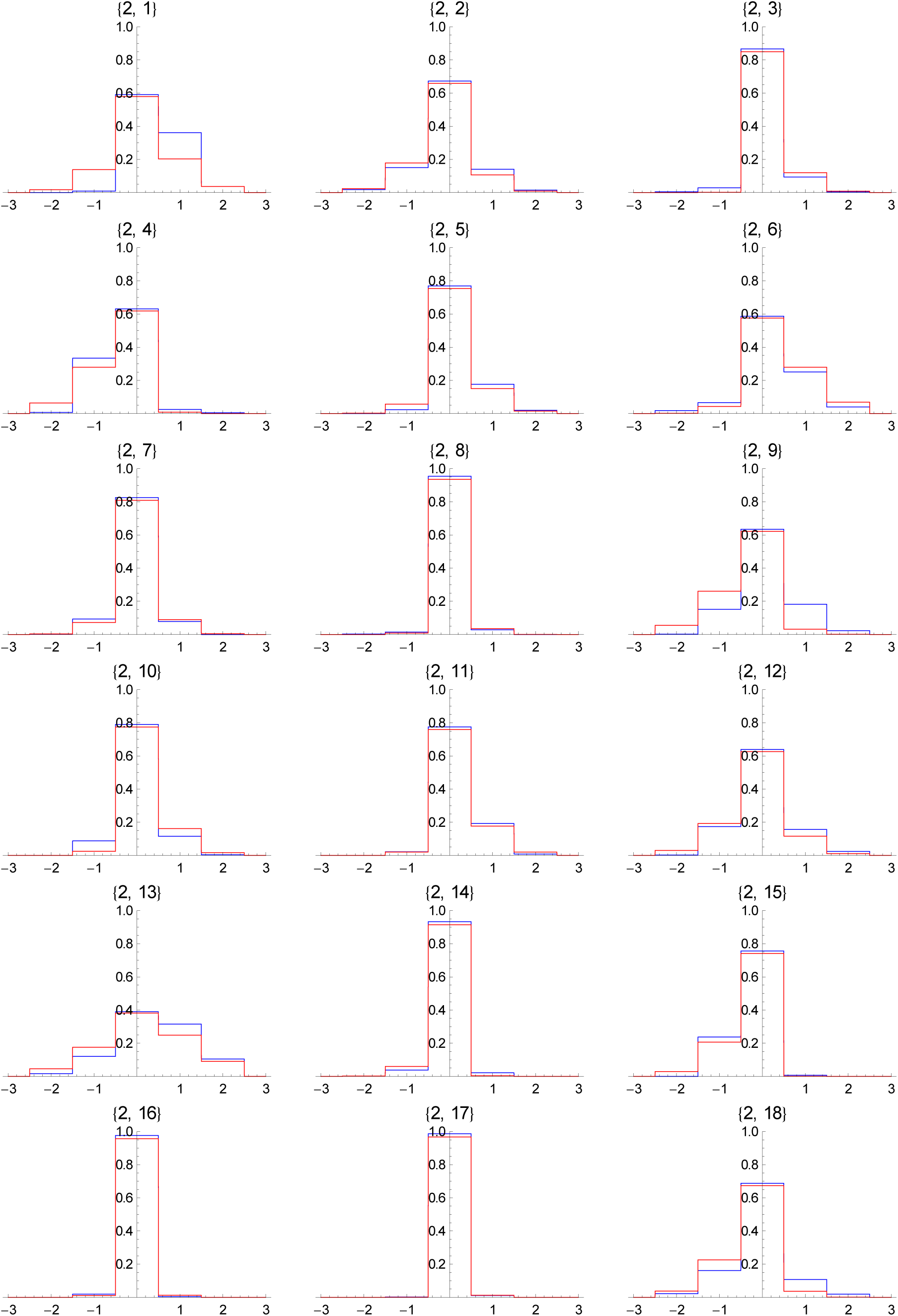

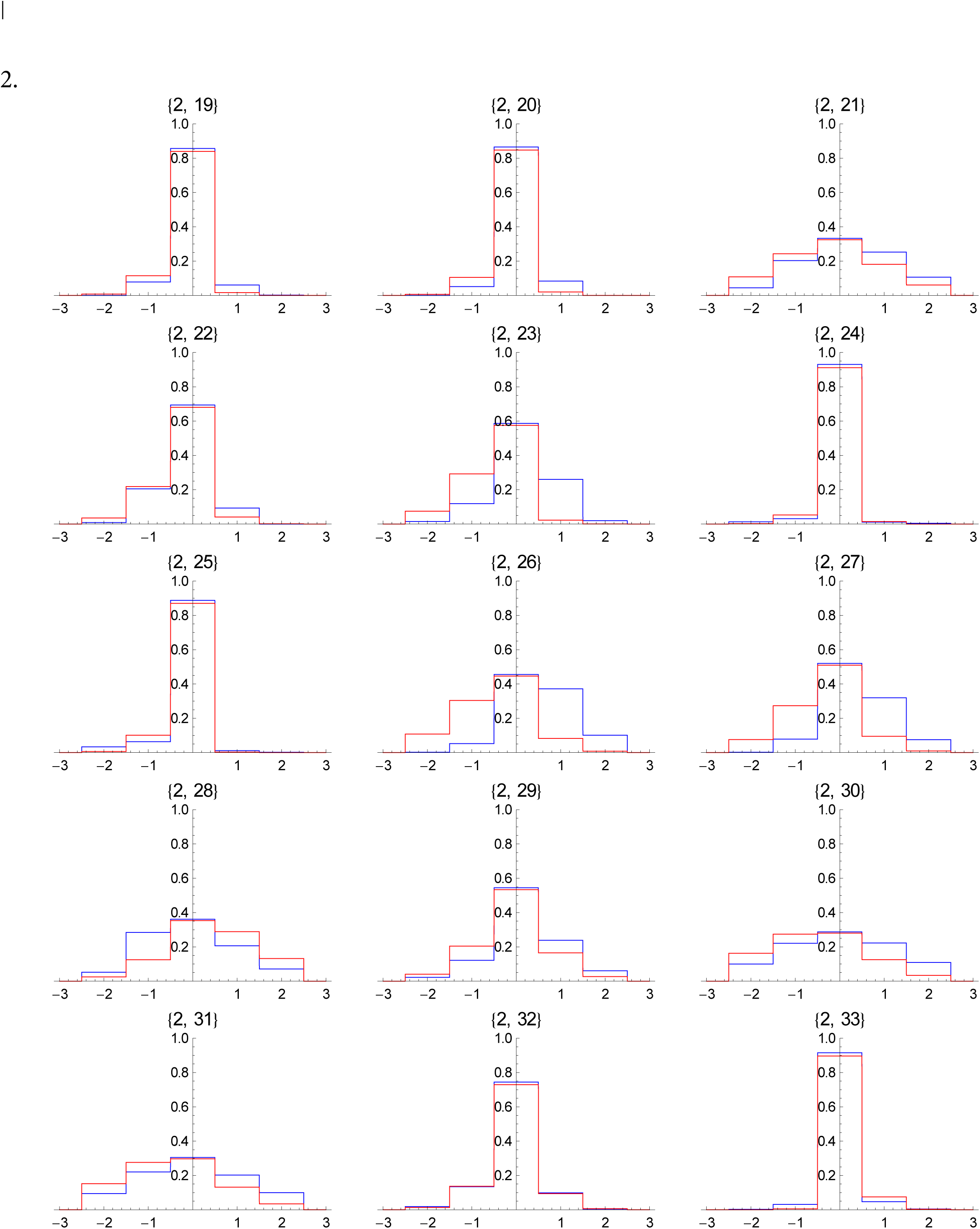

## SI 2 : Random generation of branching spectrum

The purpose of this is to show how branching produces distribution of times (spectrum) which are similar to actual experimental data. In each example the TMRCA is 200 generations but whith different values of the branching constant a.

Regarding this as pseudo R1b1a2 distributions we assume same size sample

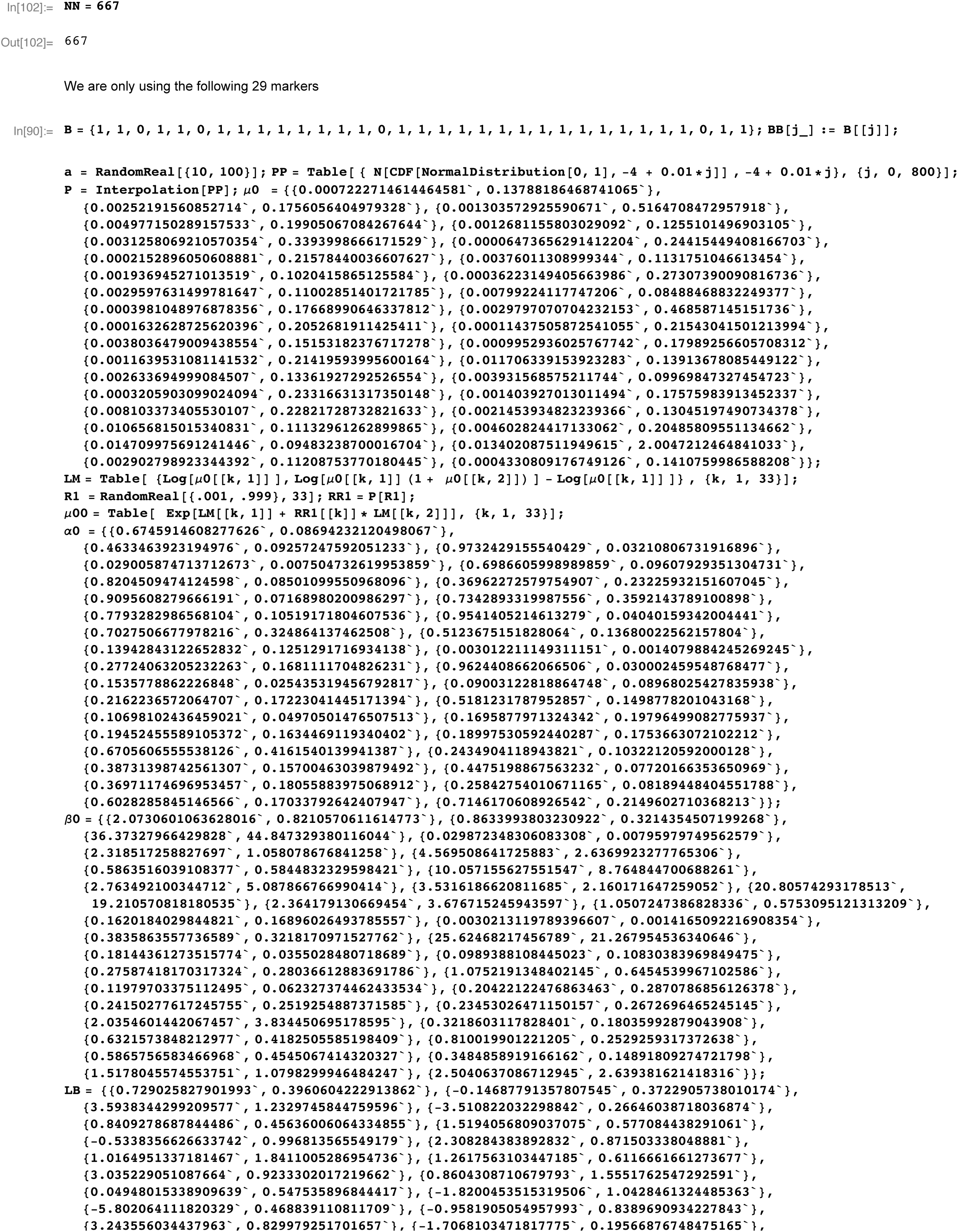

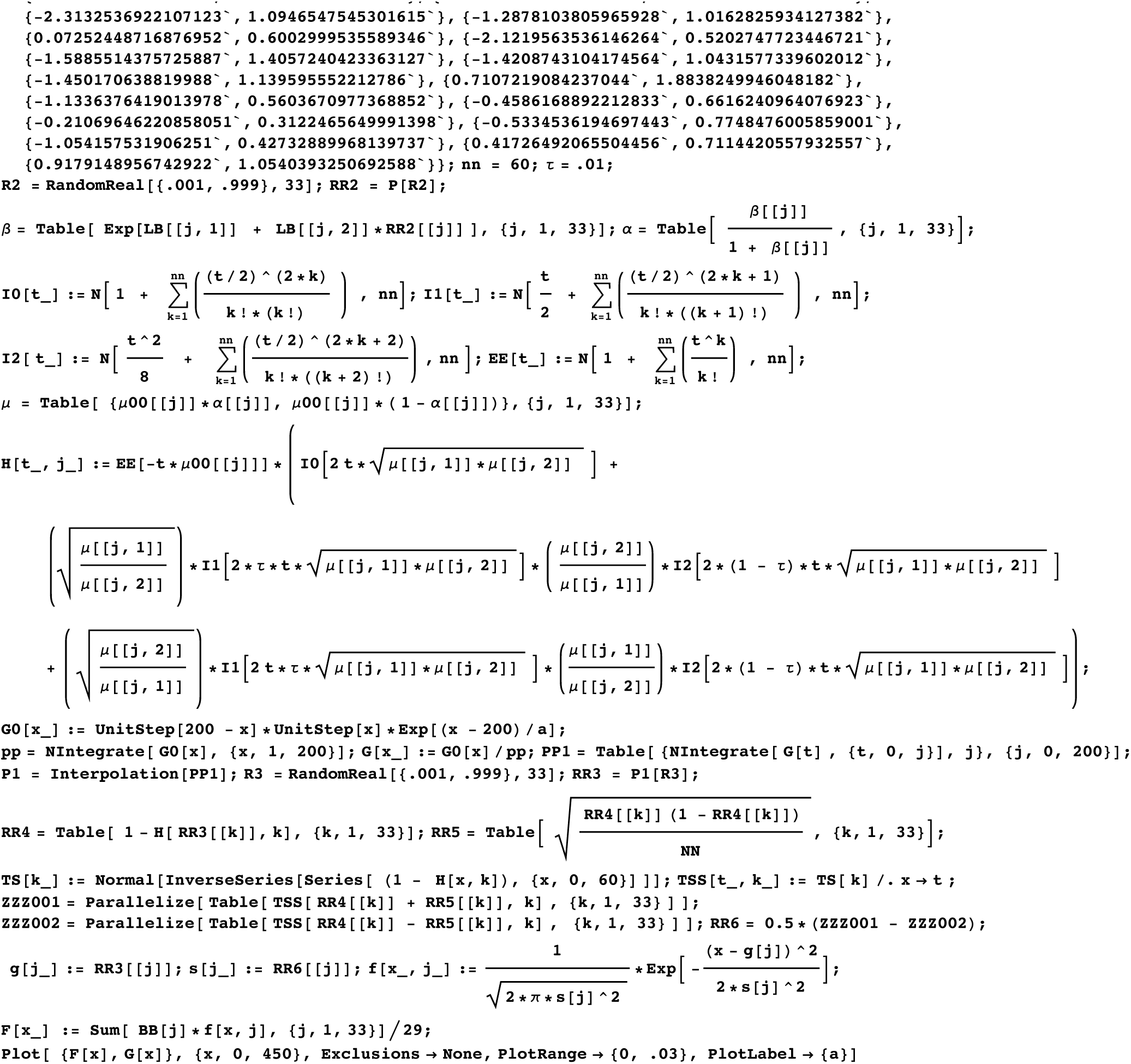

Each of the following shows the Branching Spectrum (Blue) resulting a spreadout generation of branches (density function in Red)

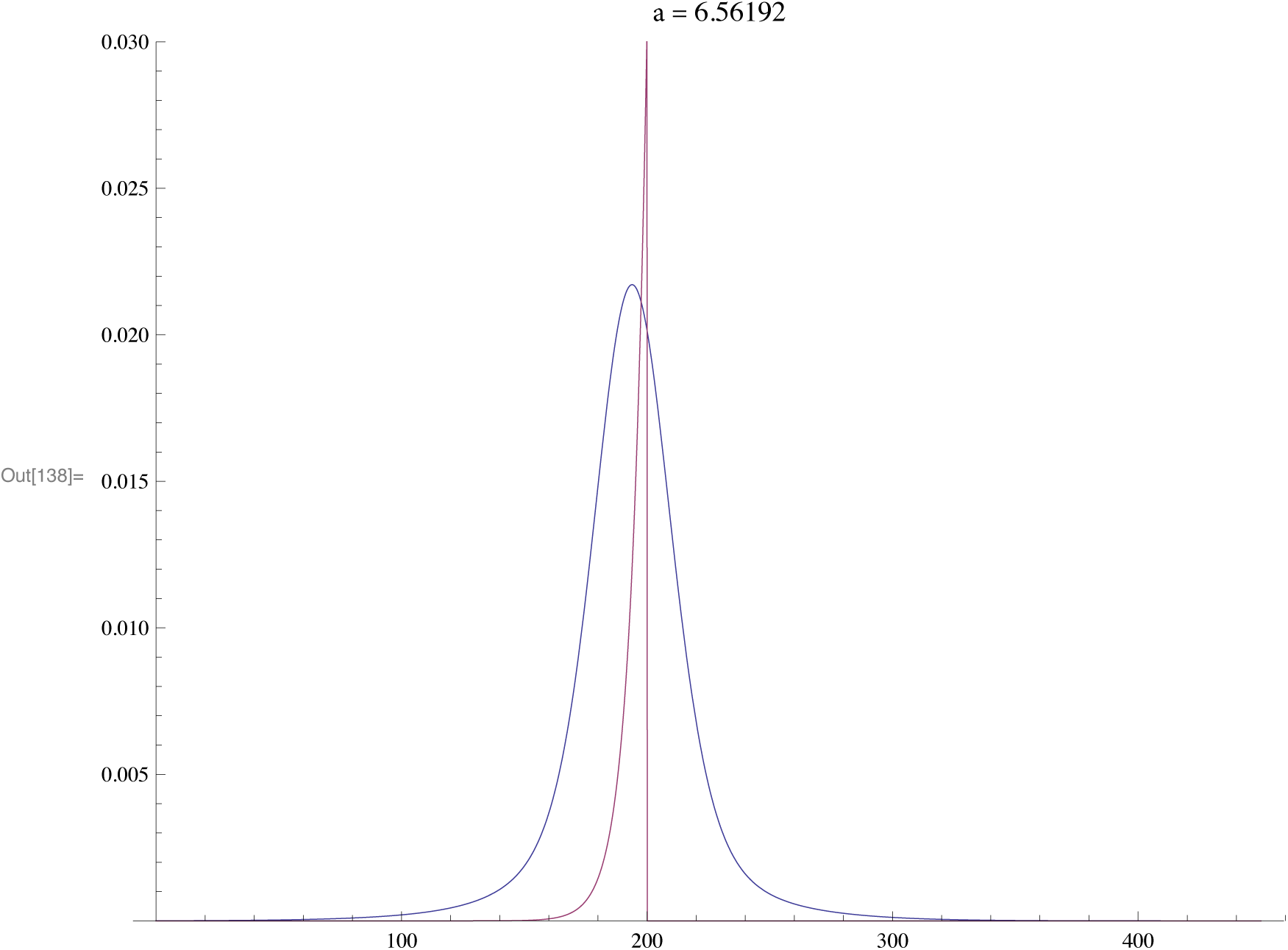

For a fast expansion the mean is almost exactly the TMRCA

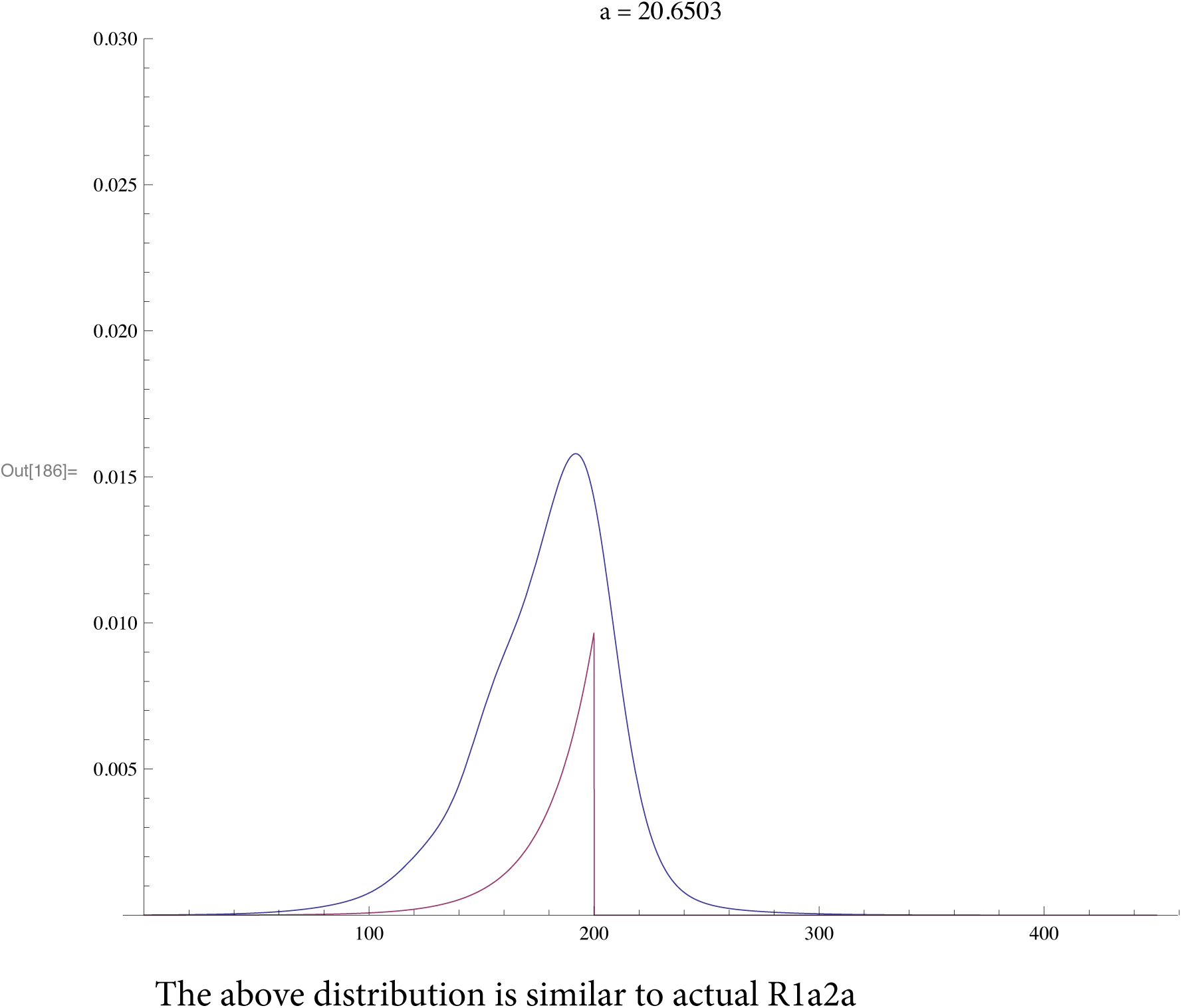

The above distribution is similar to actual R1a2a

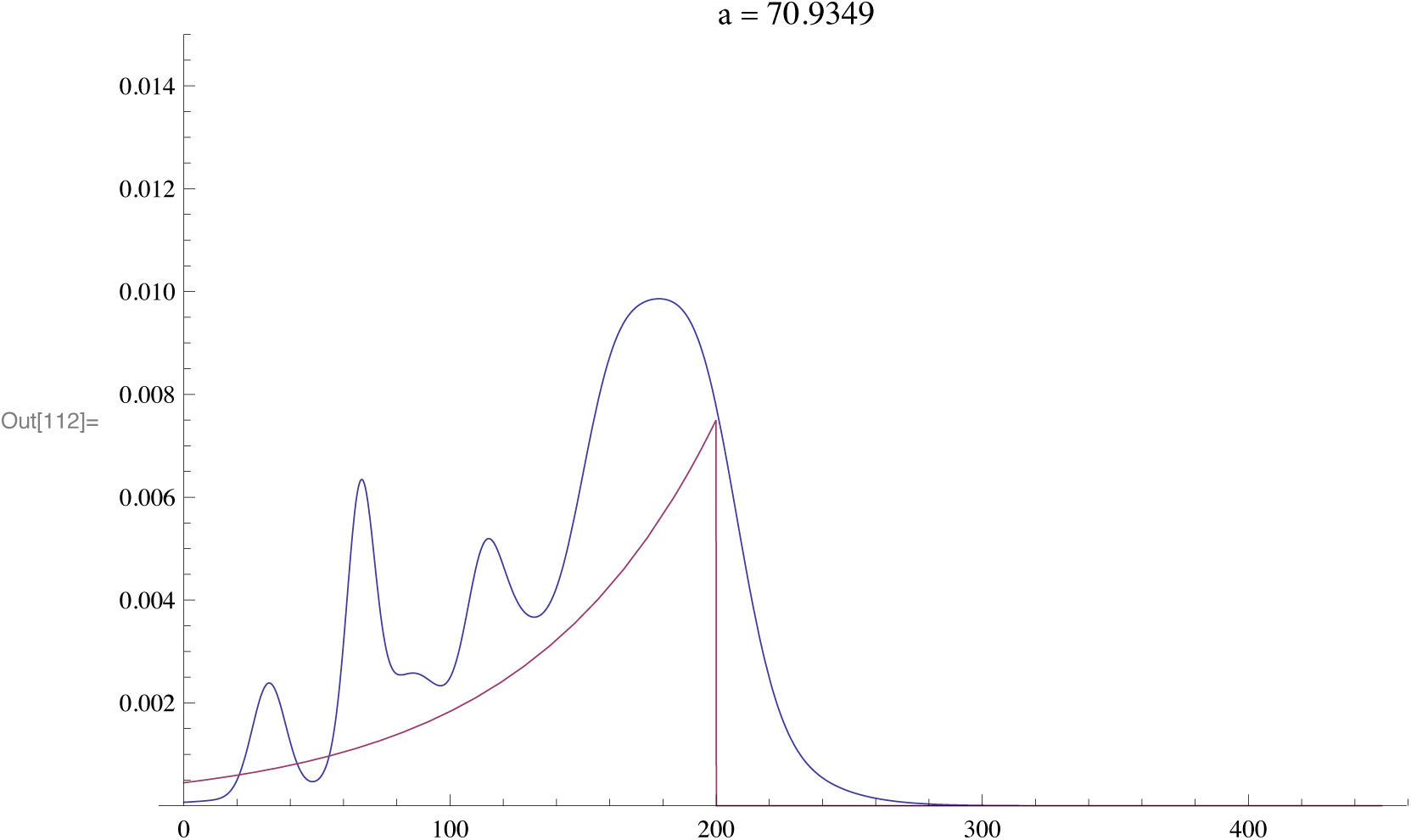

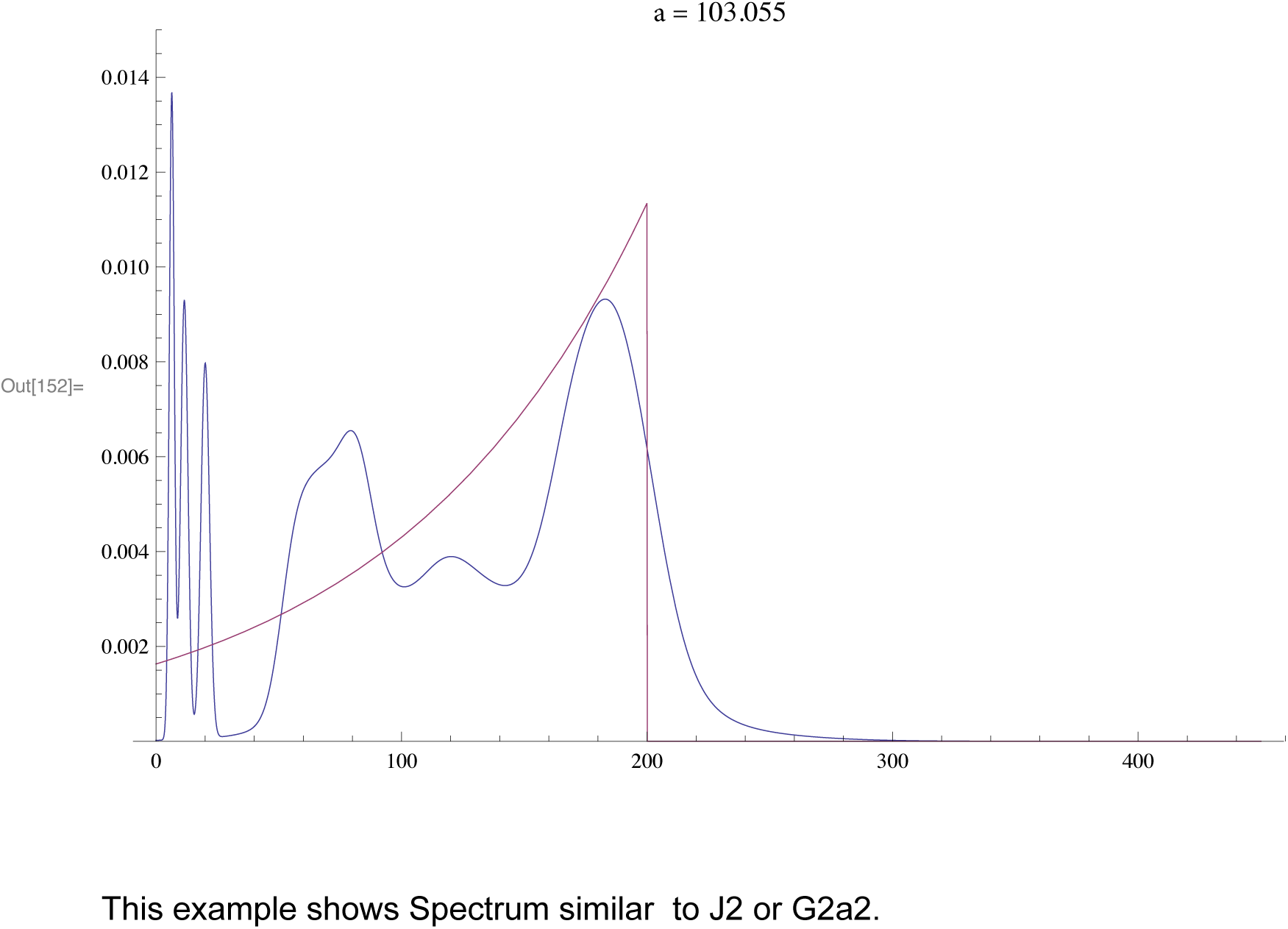

This example shows Spectrum similar to J2 or G2a2.

## SI 3 : Mathematica worksheet for R1b1a2

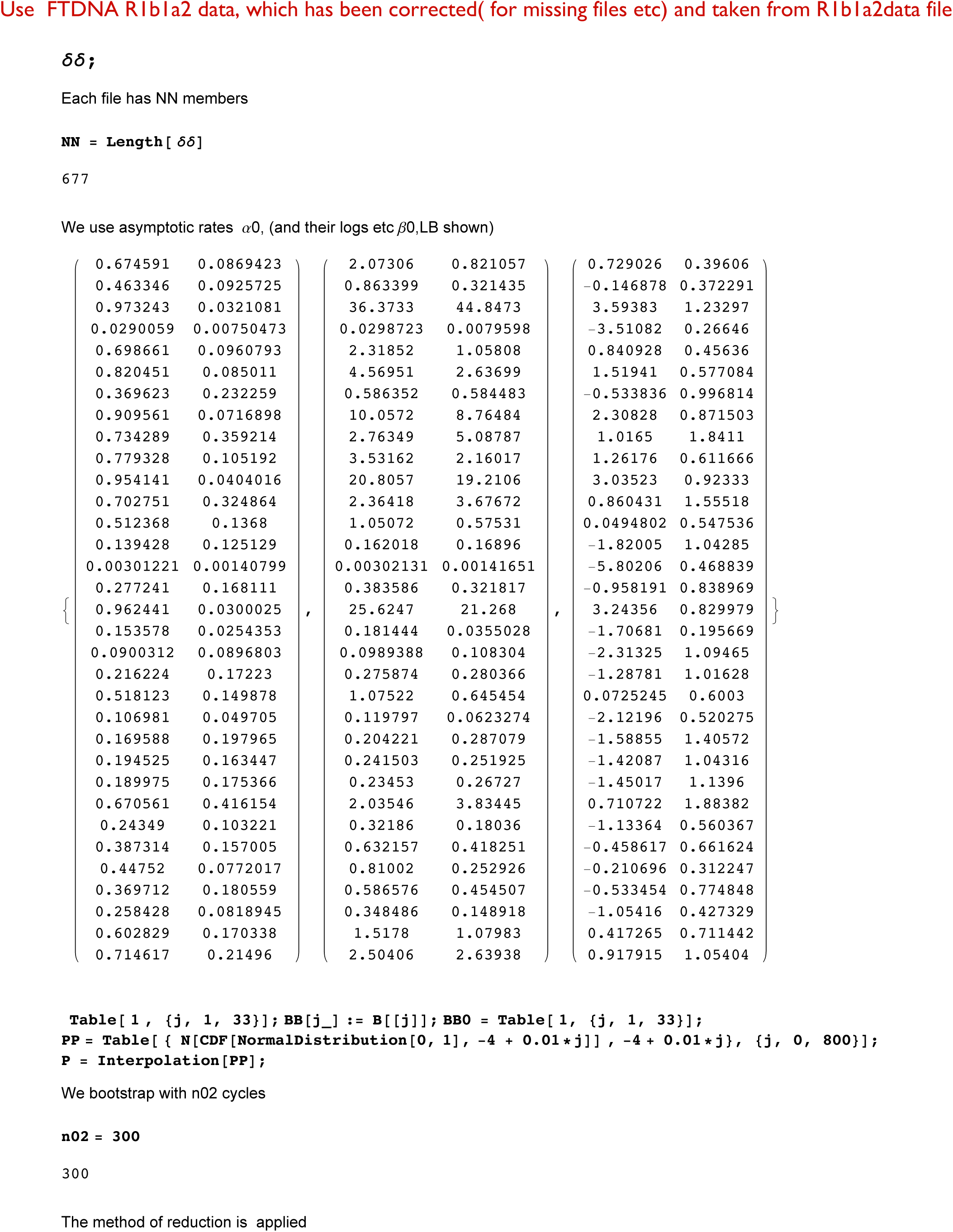

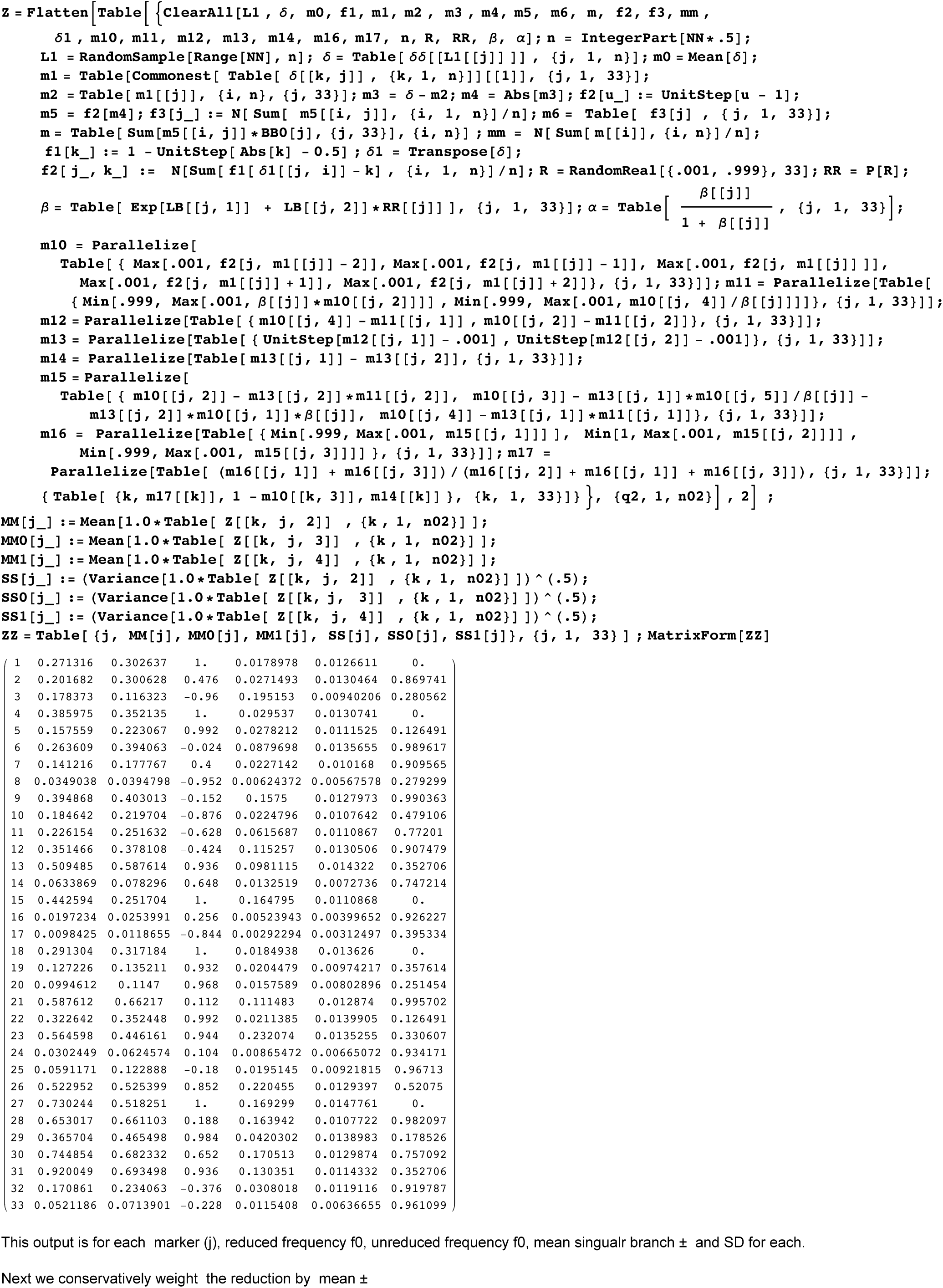

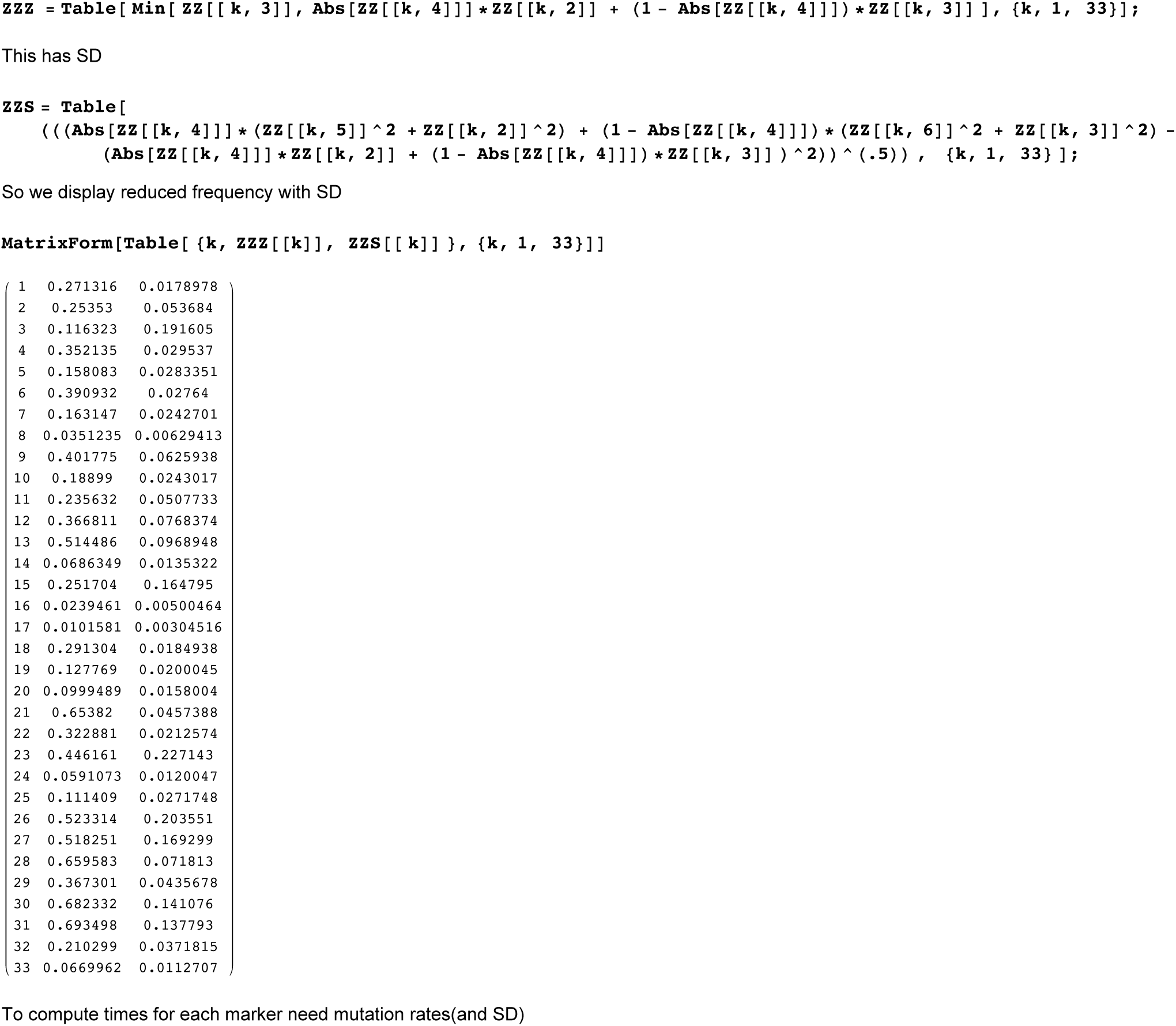

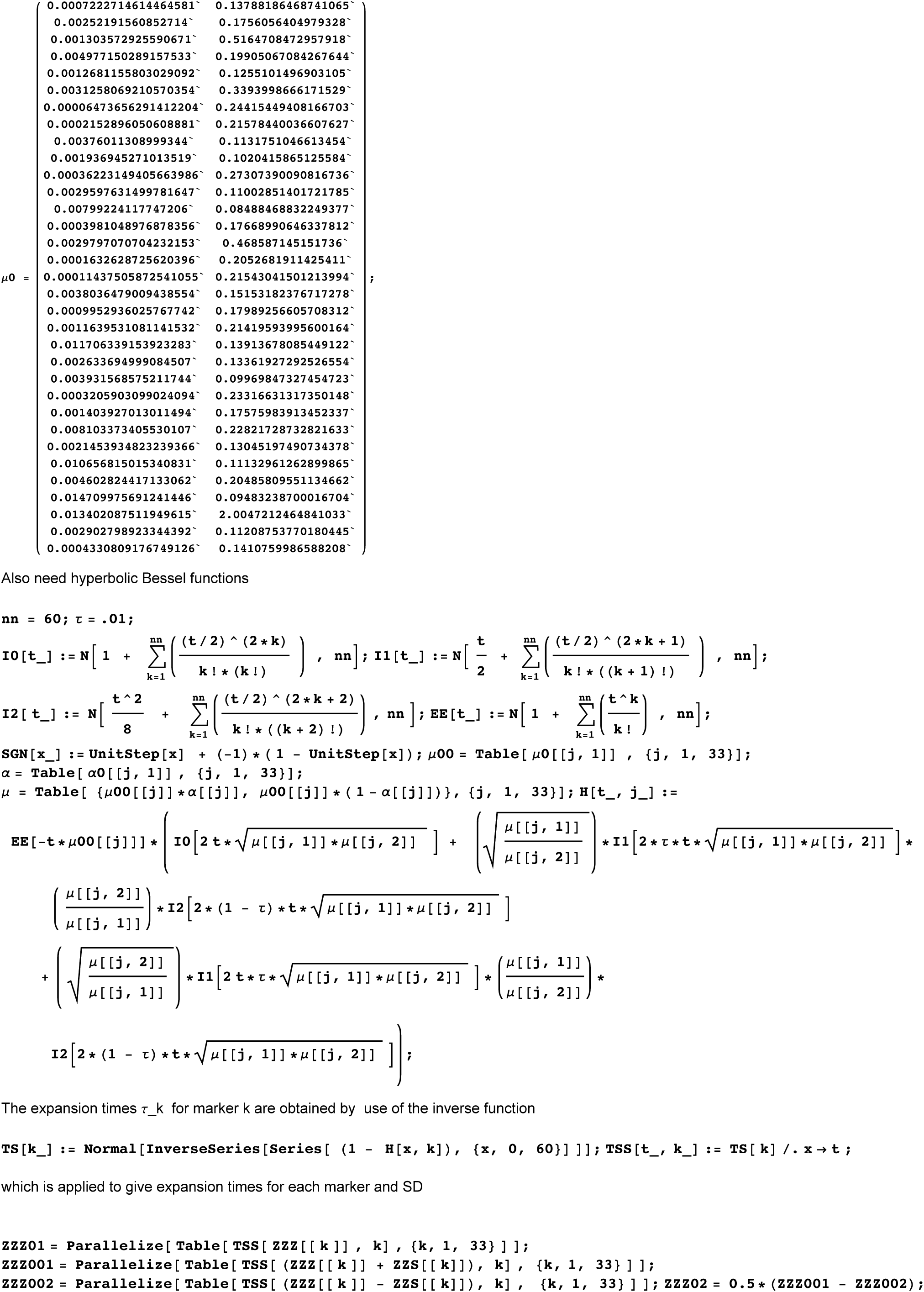

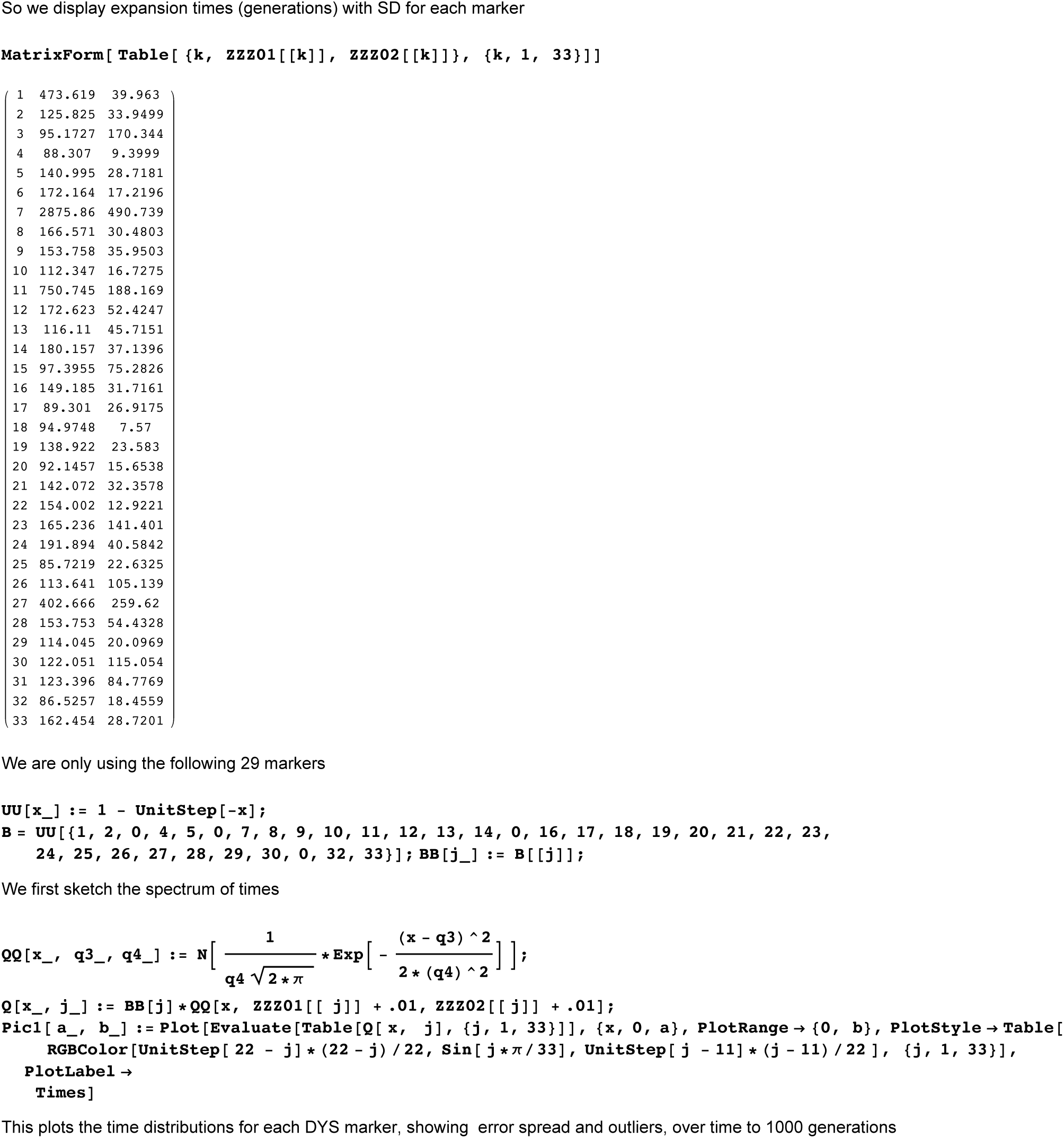

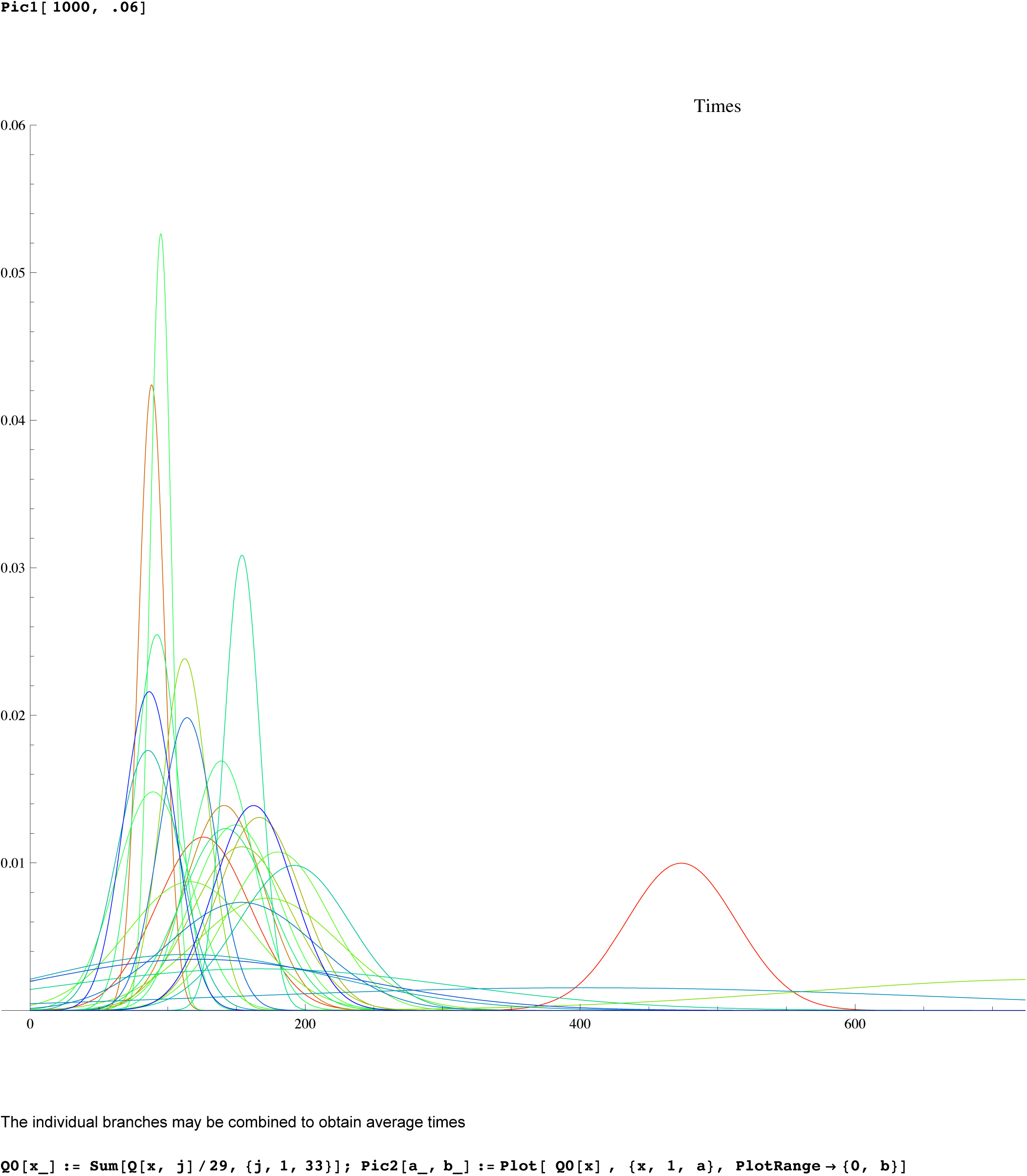

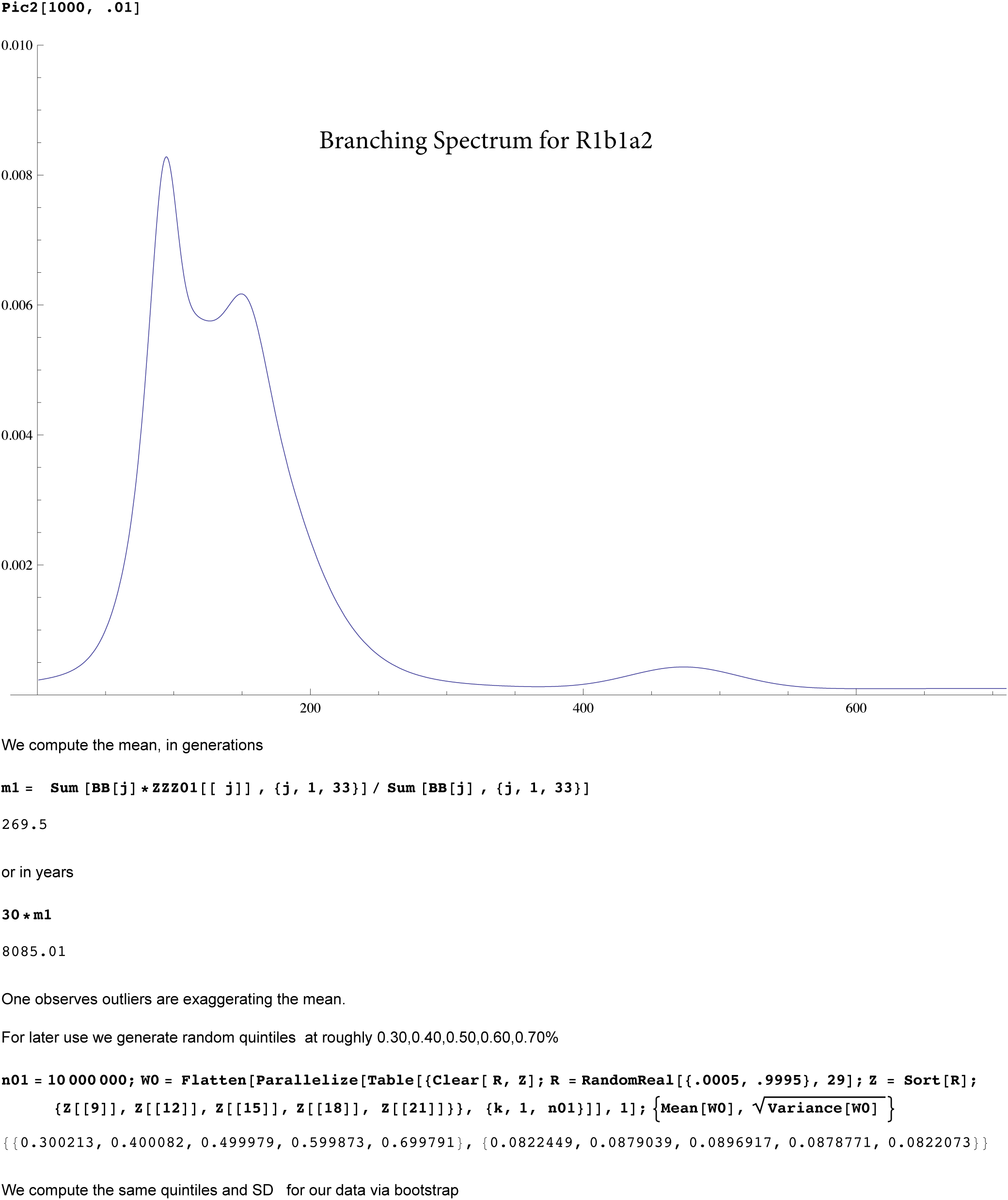

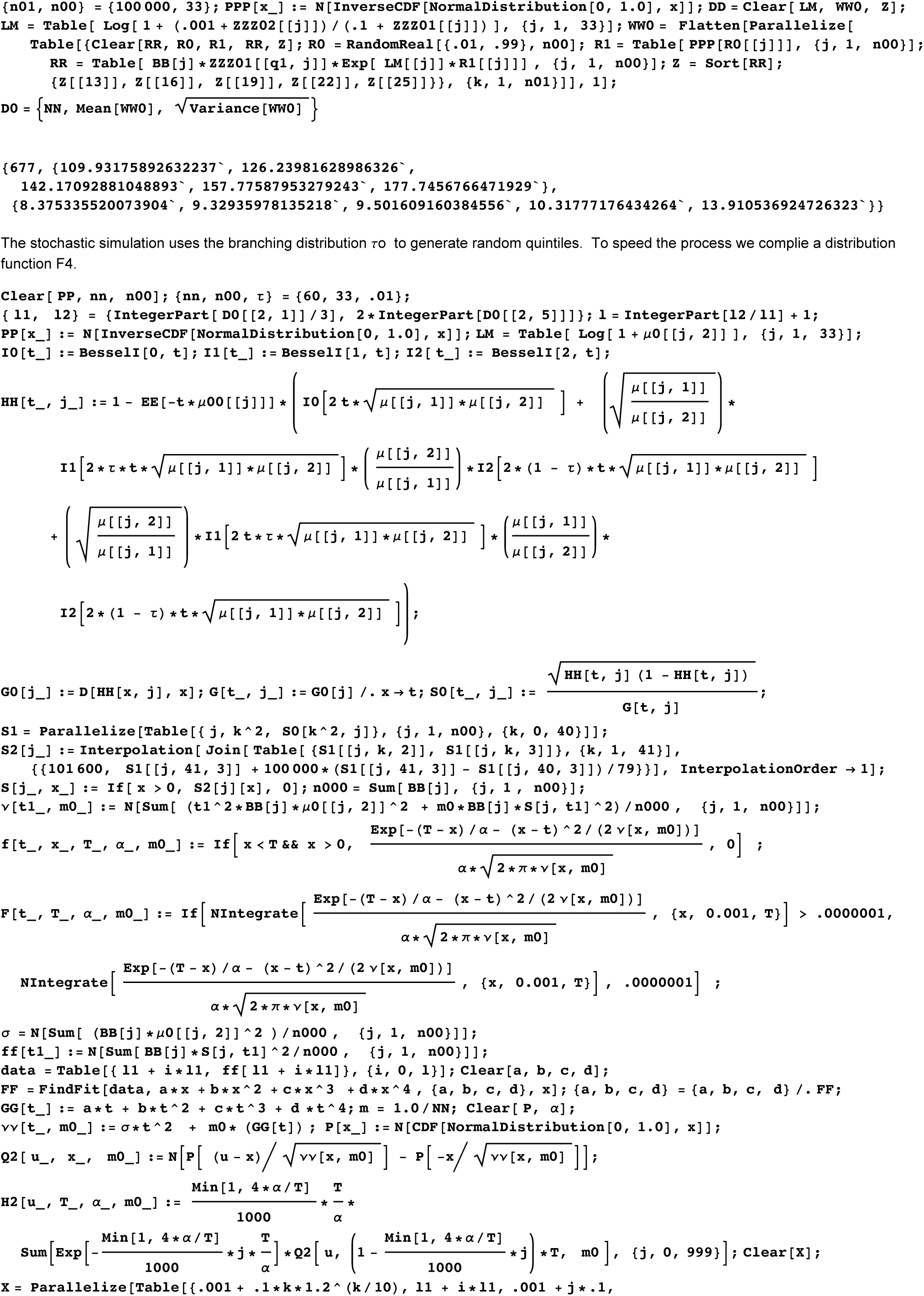

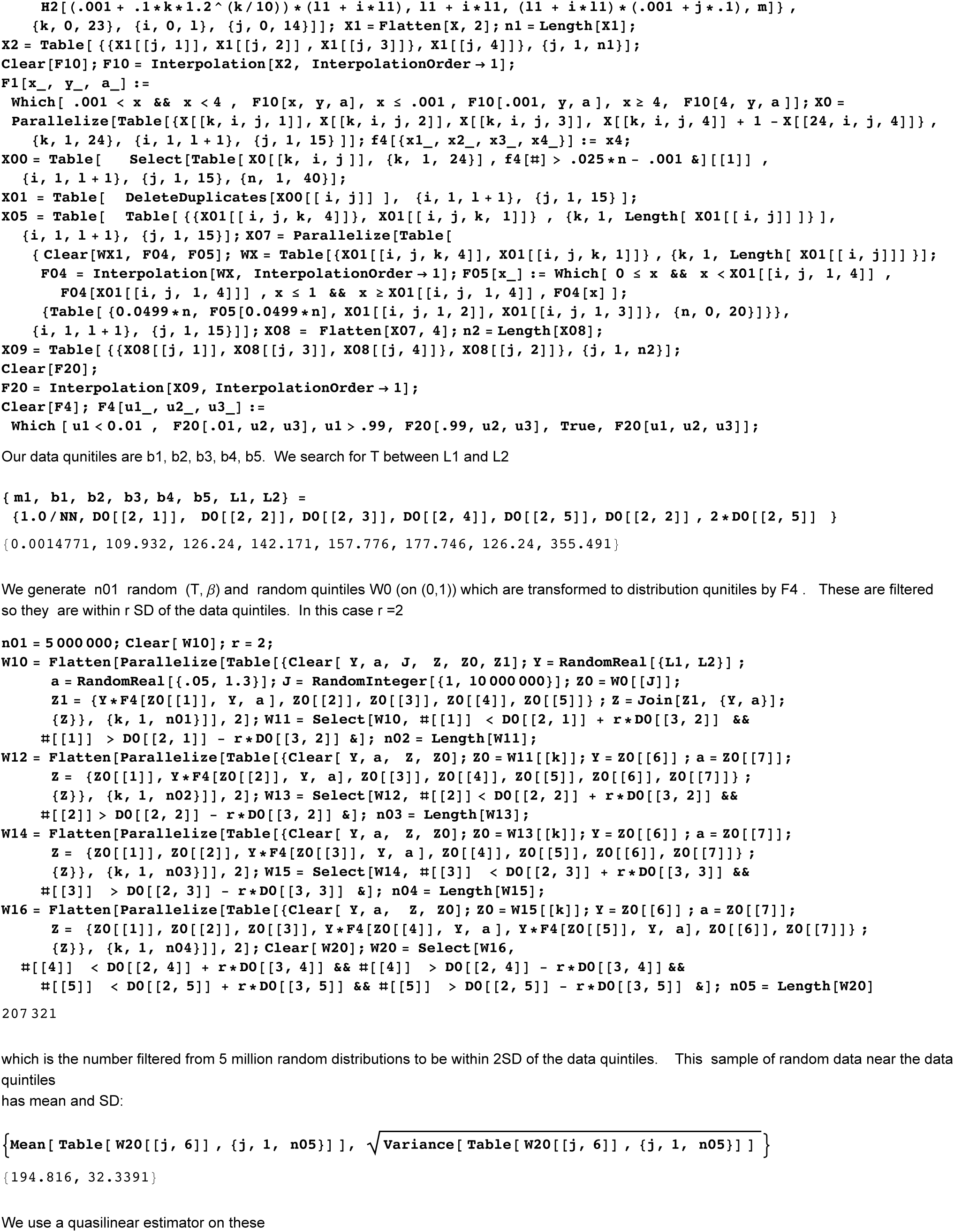

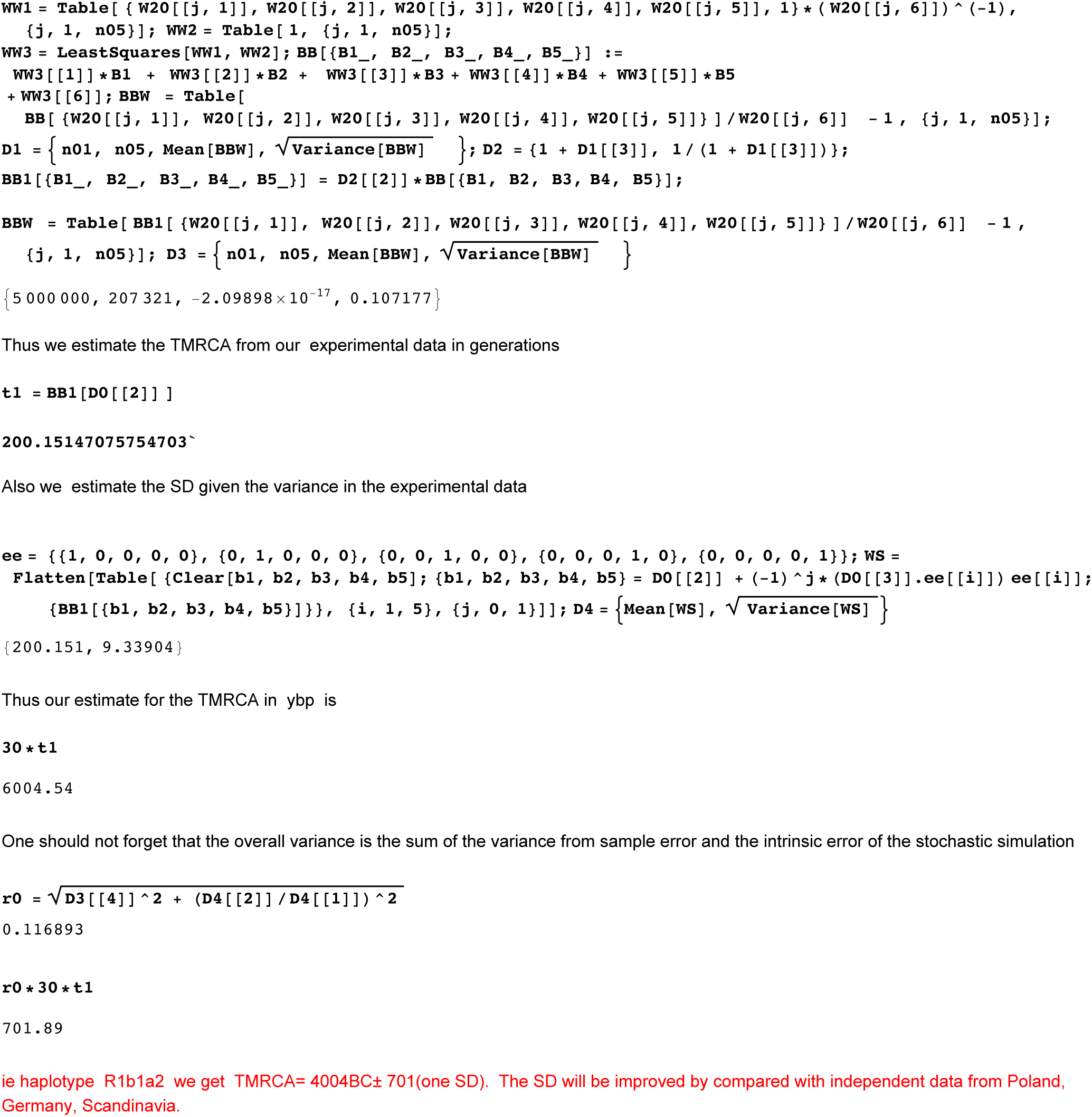

## SI 4 : Mathematica worksheet for computing asymmetric constants

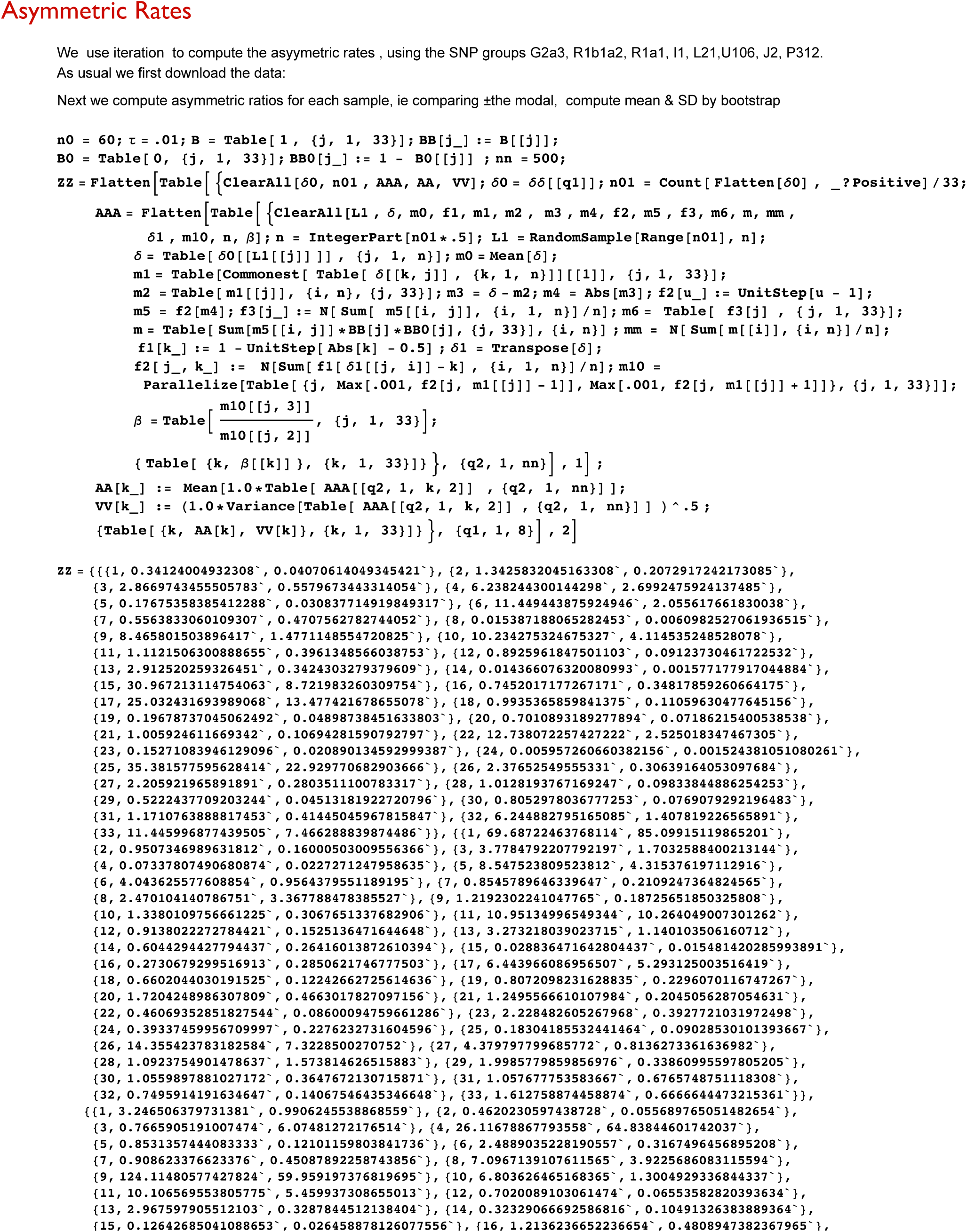

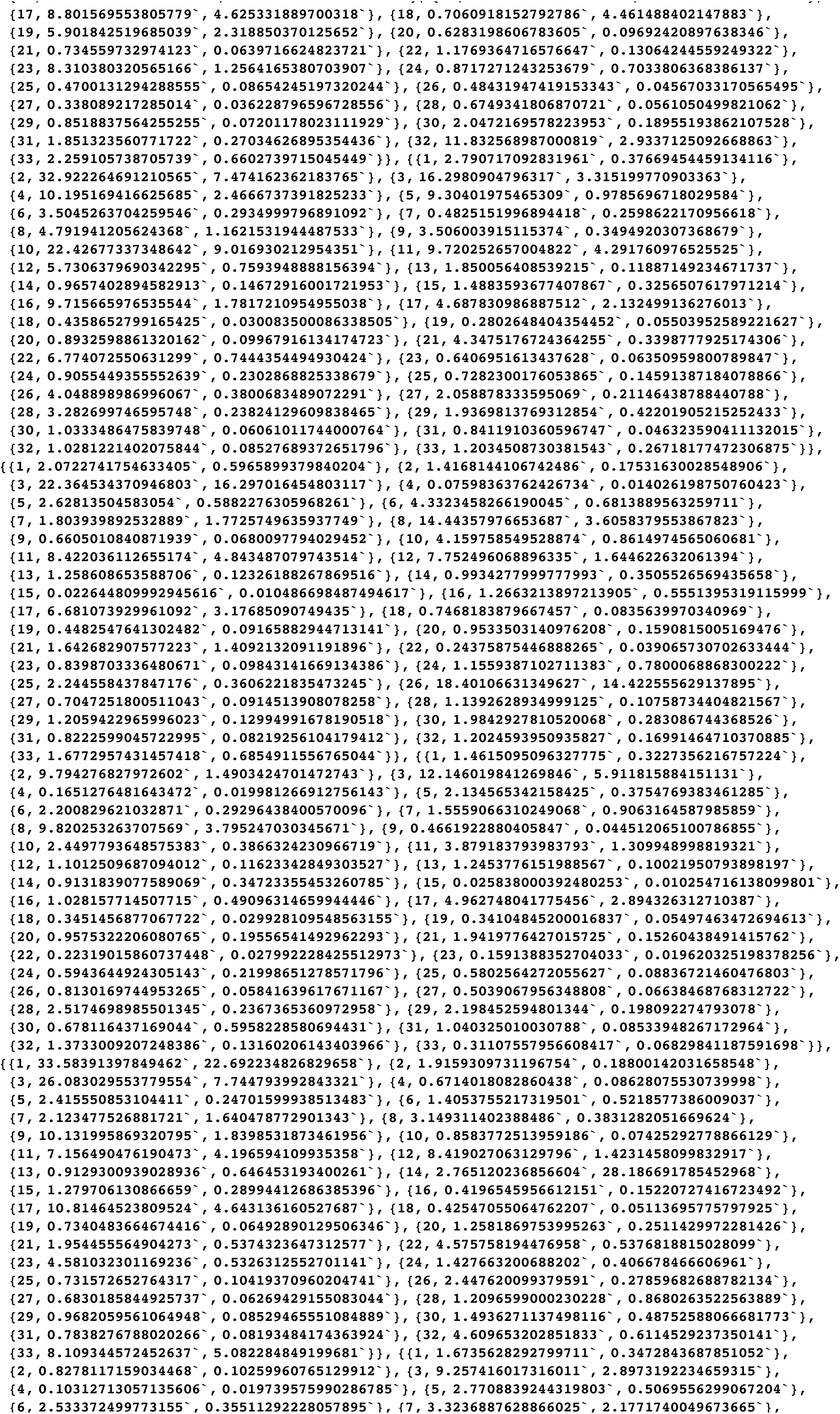

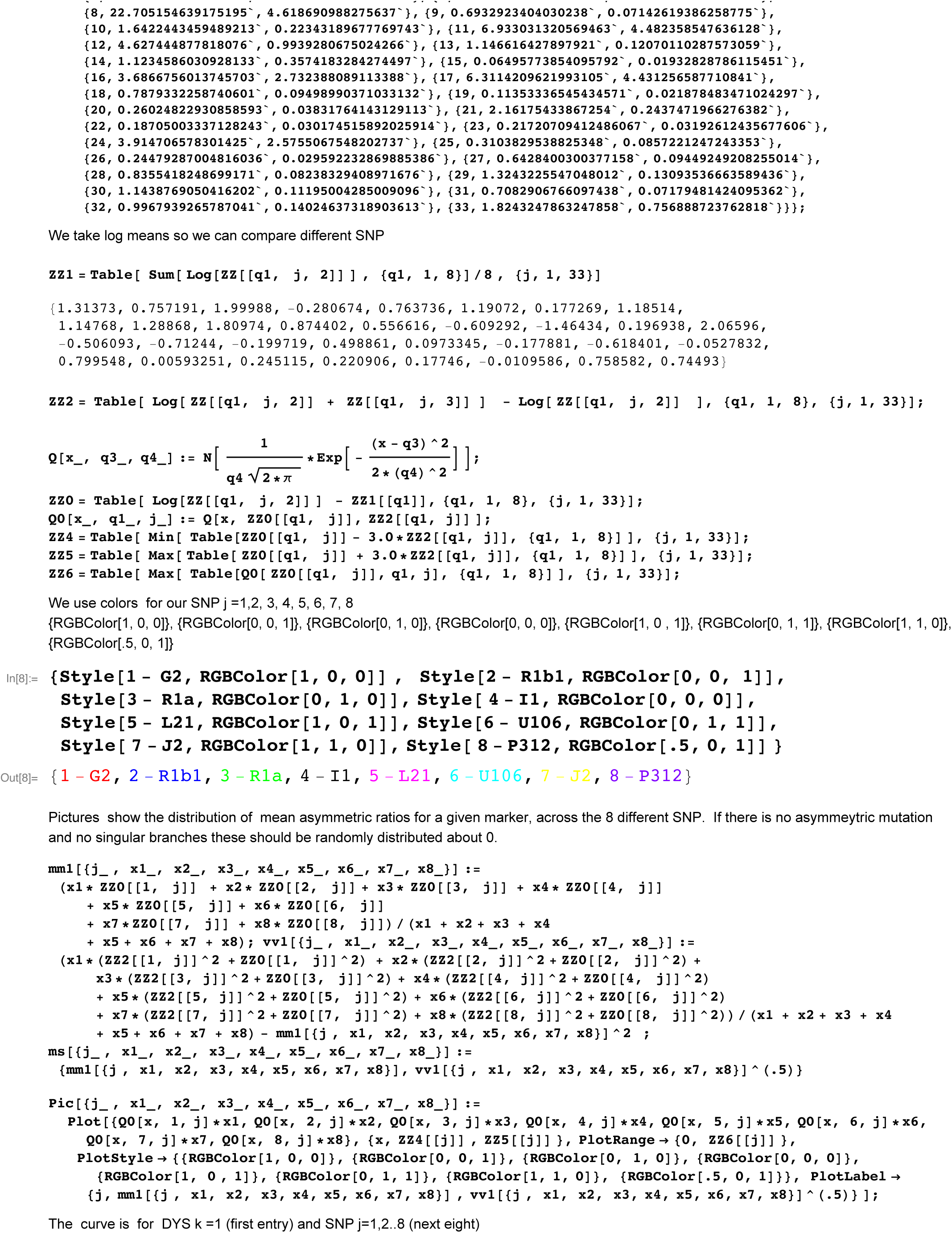

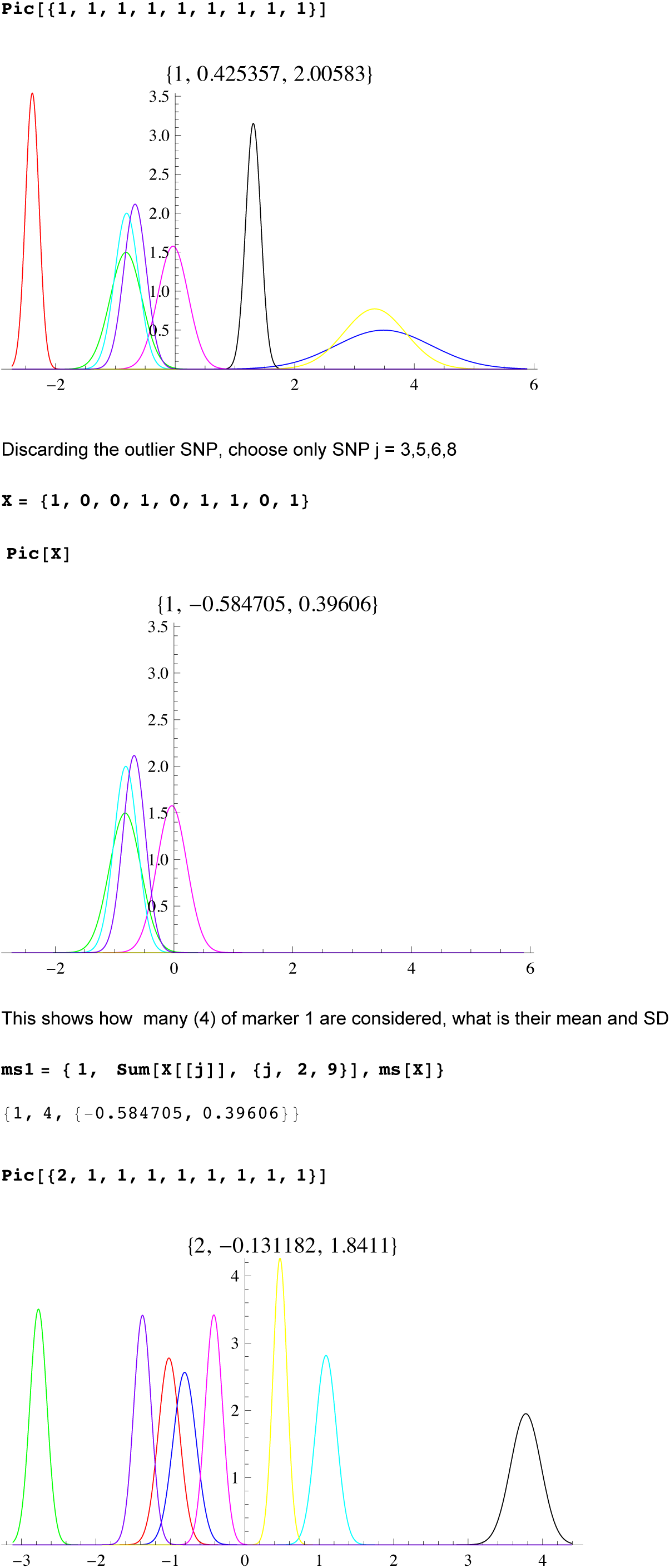

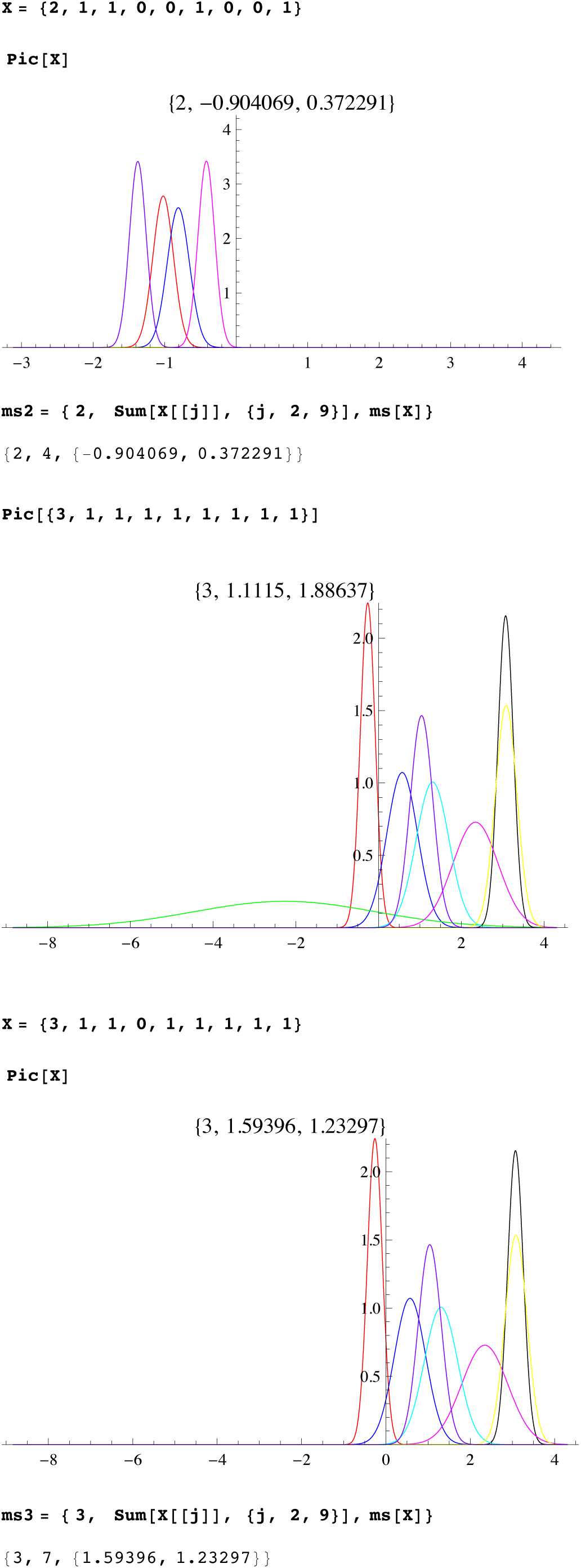

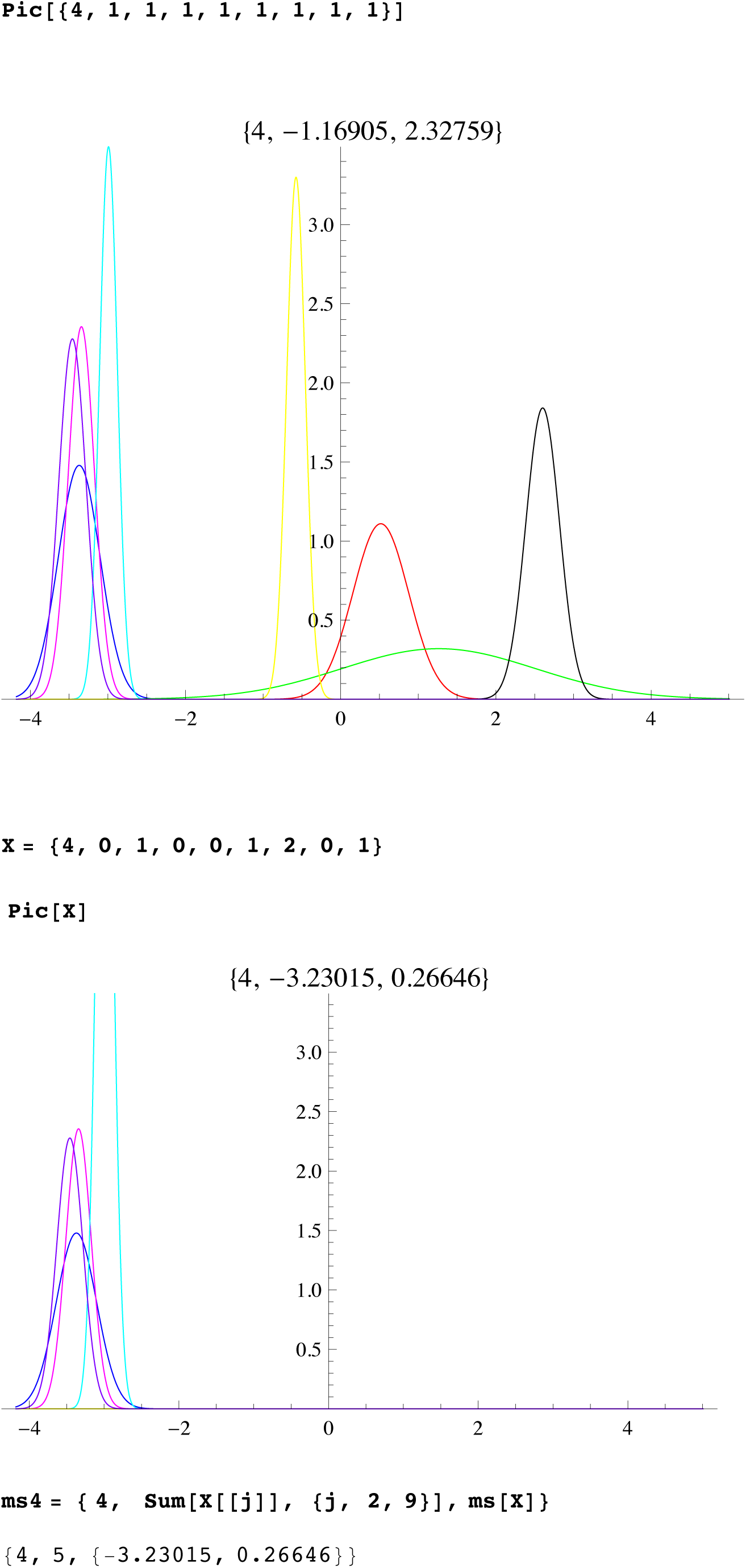

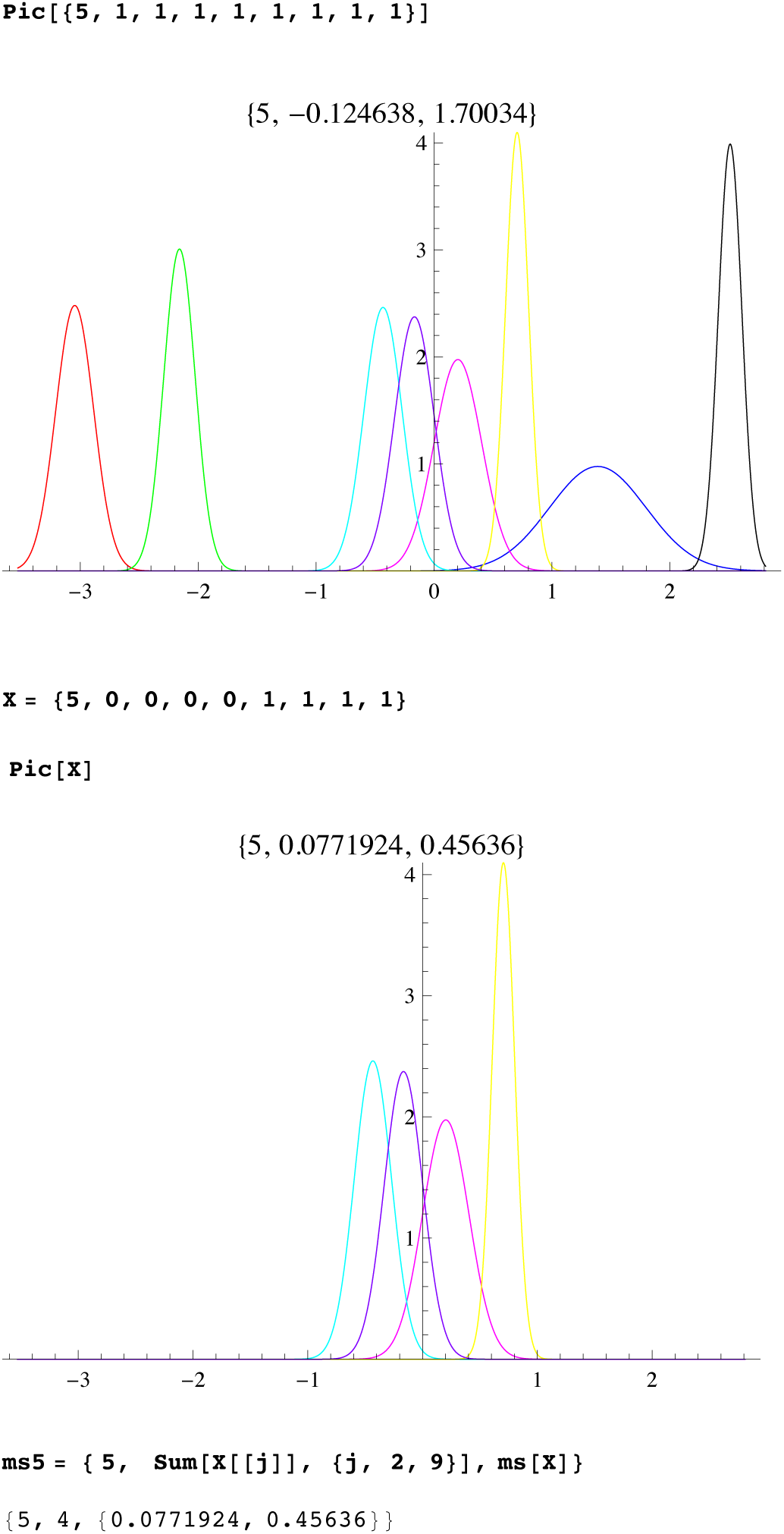

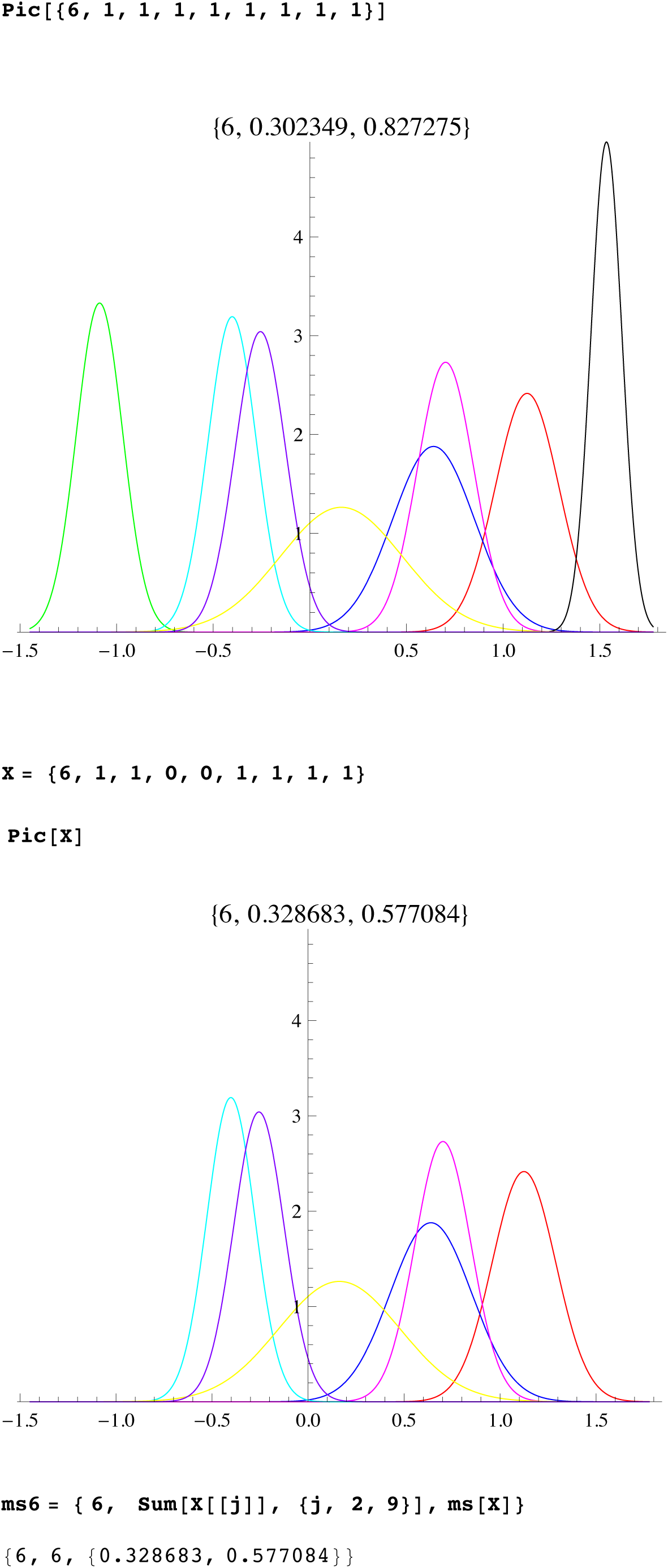

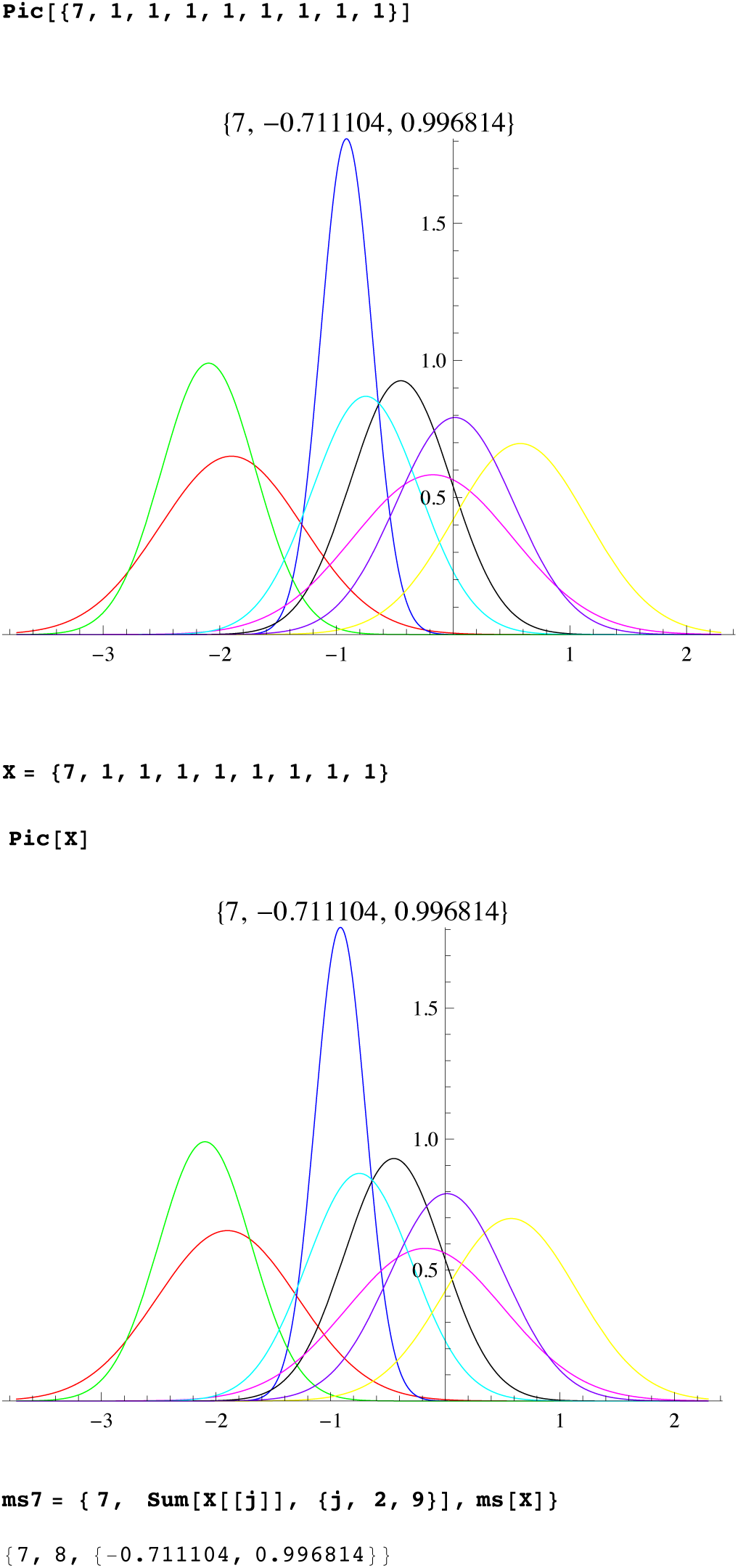

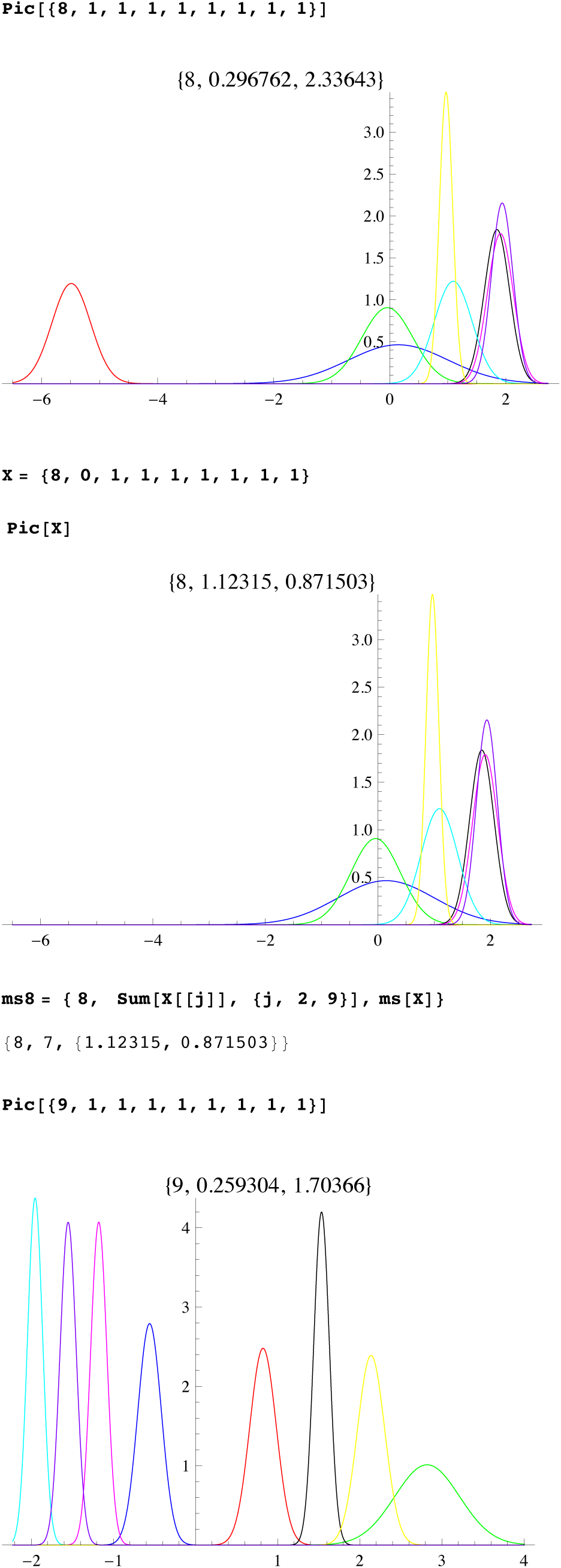

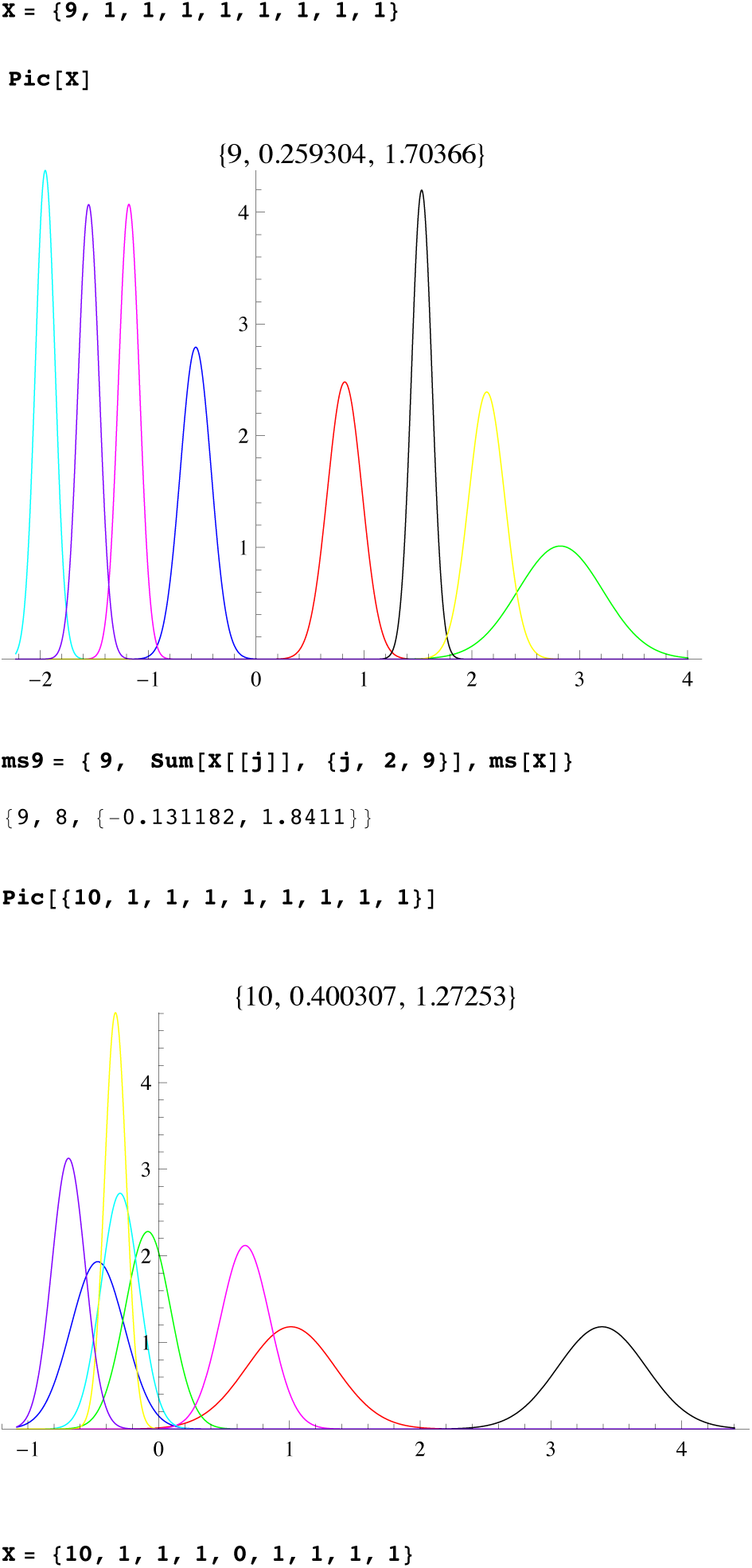

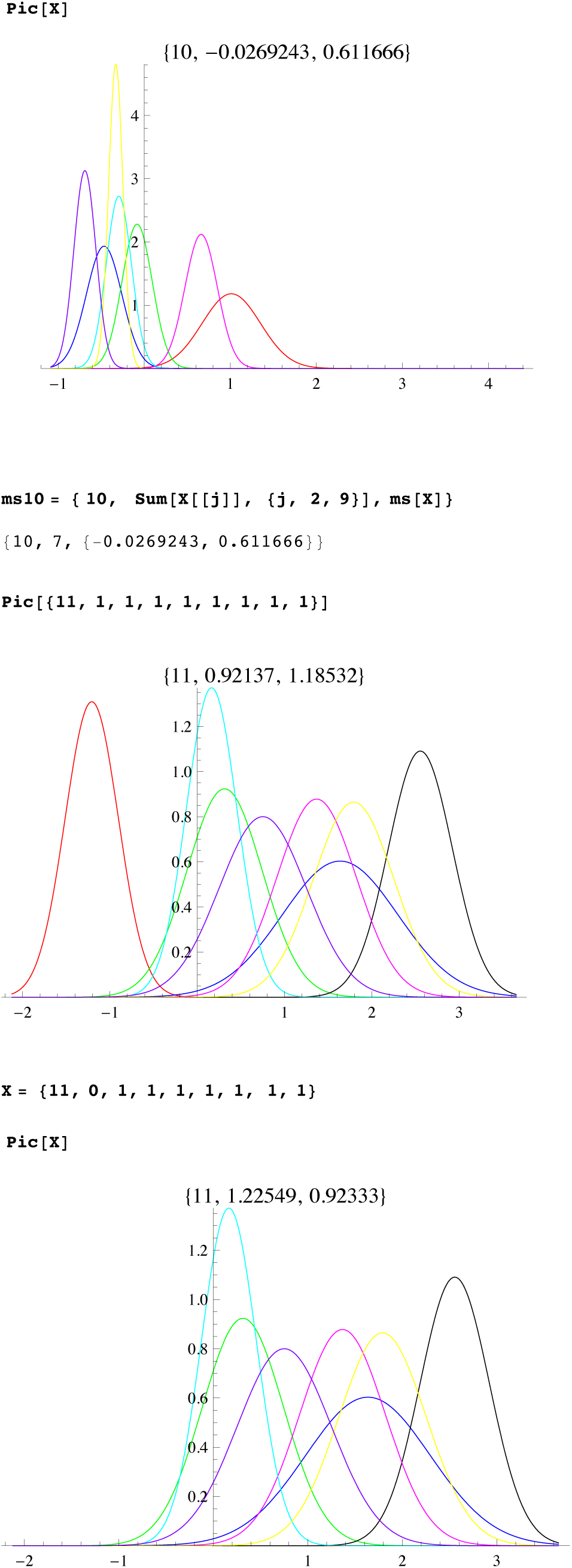

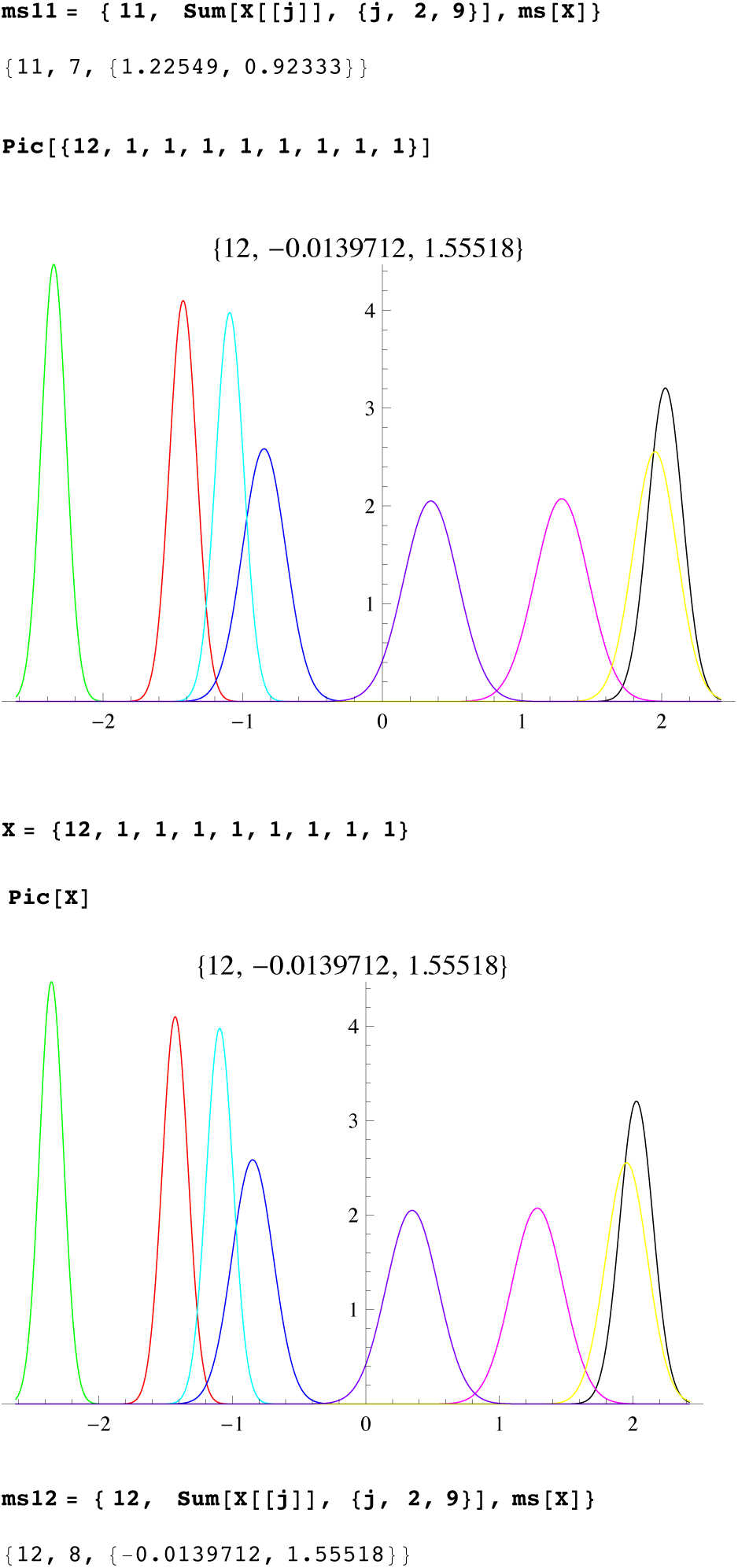

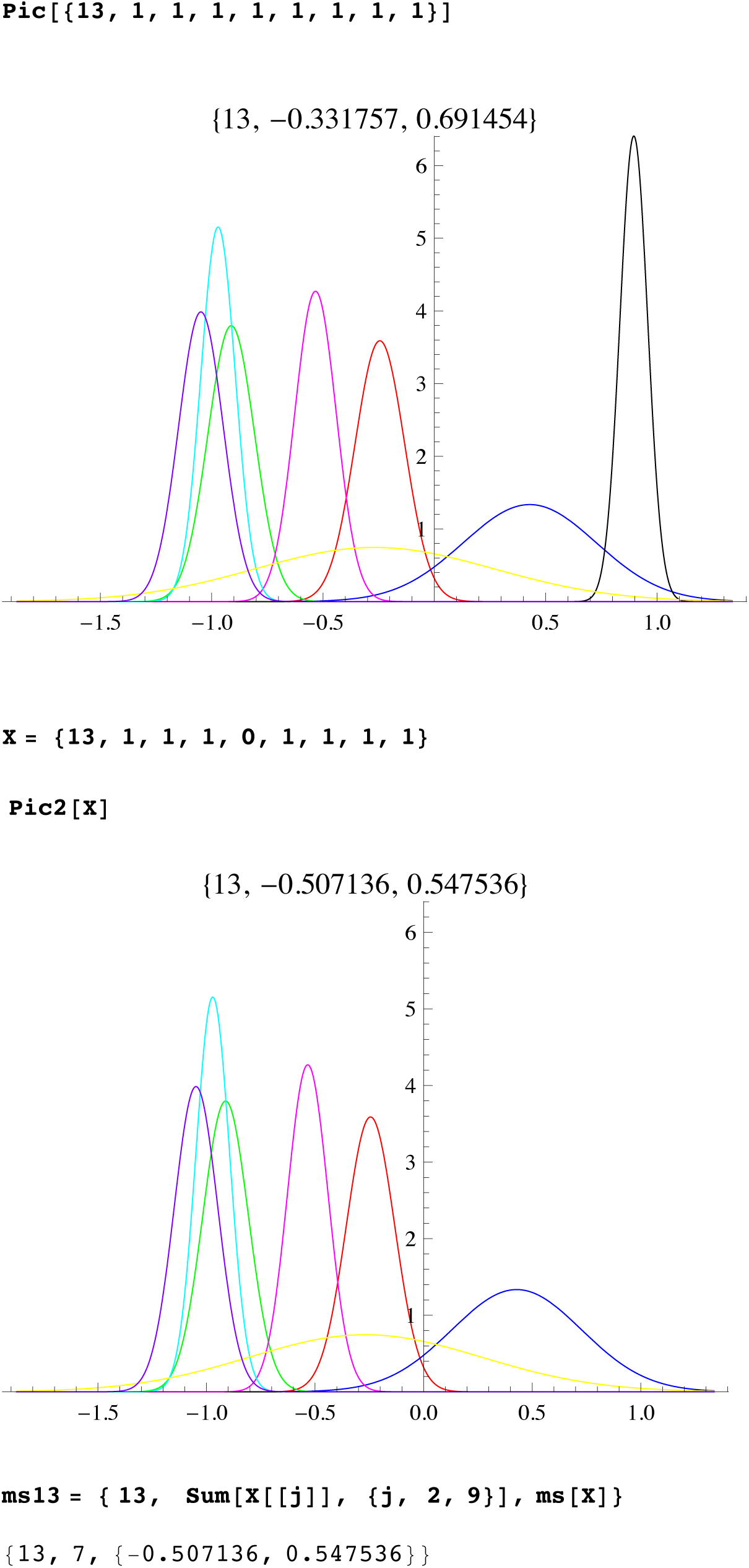

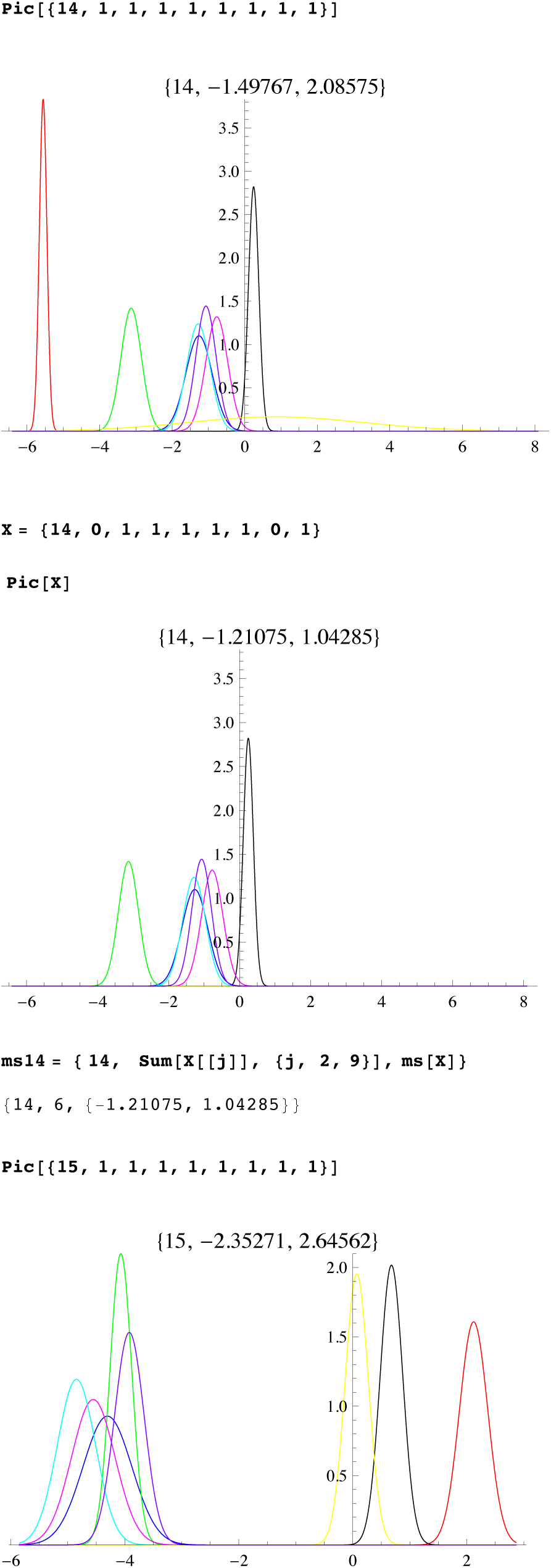

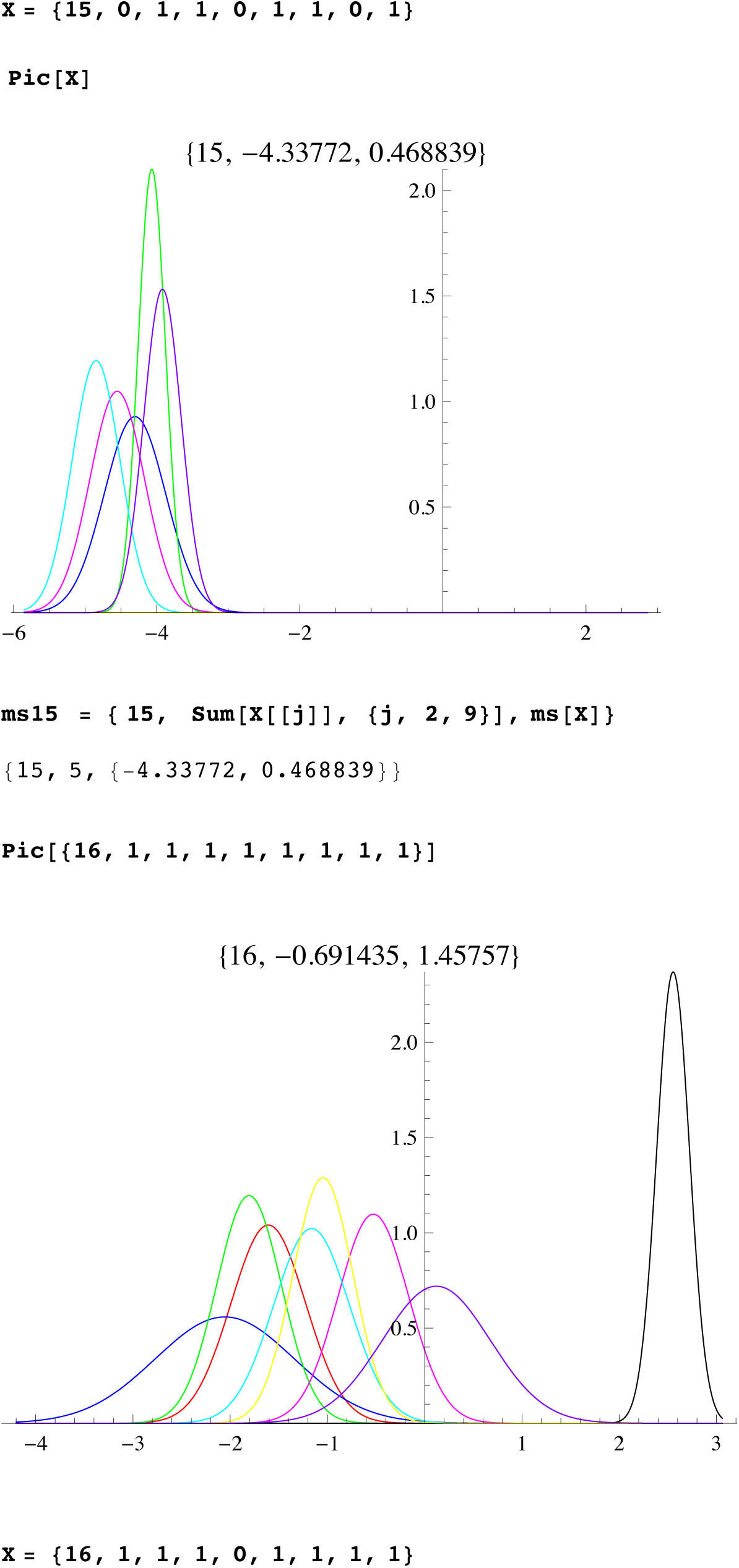

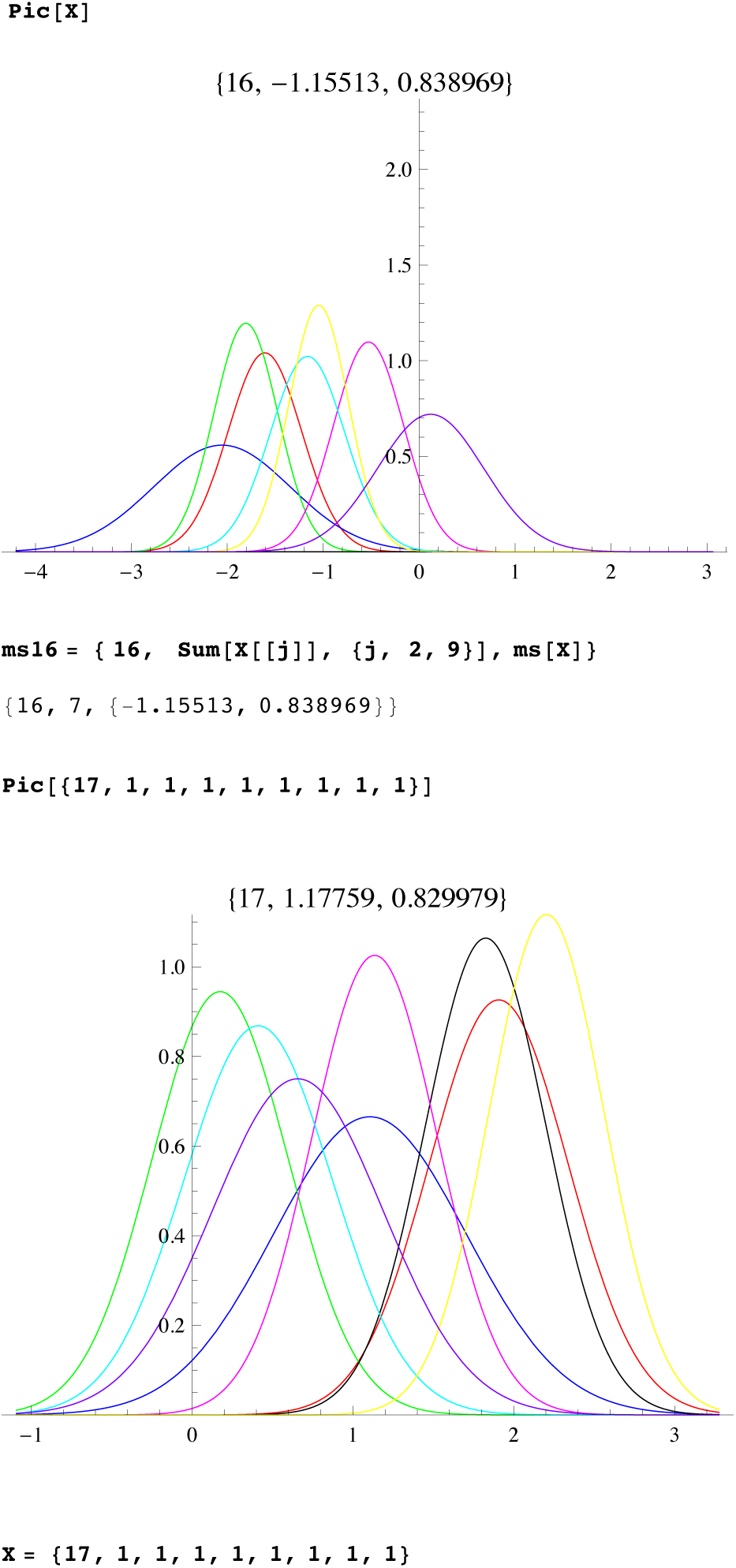

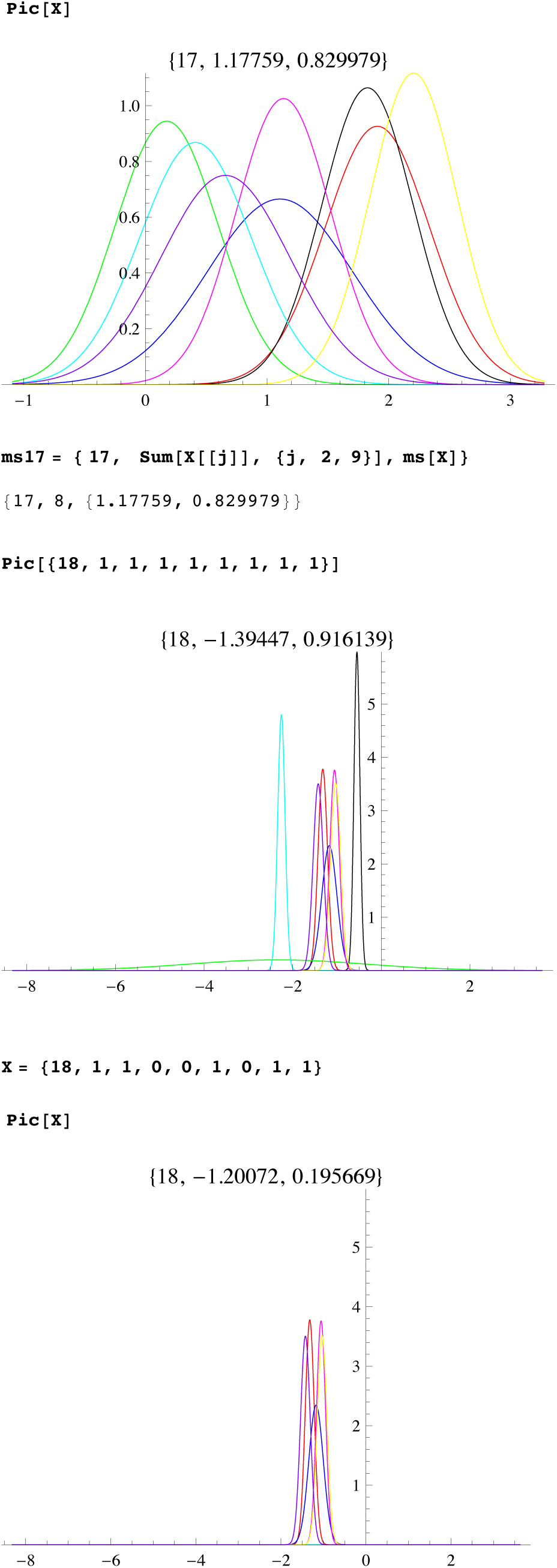

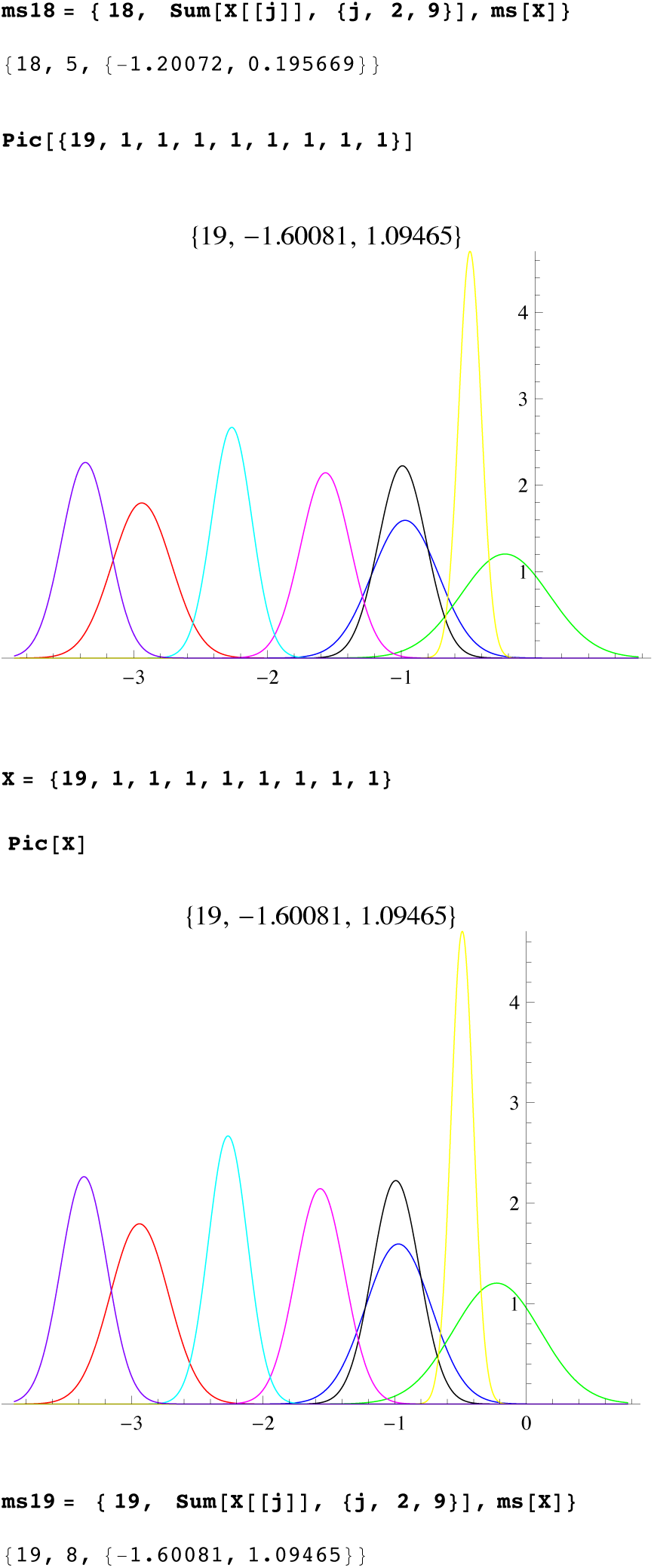

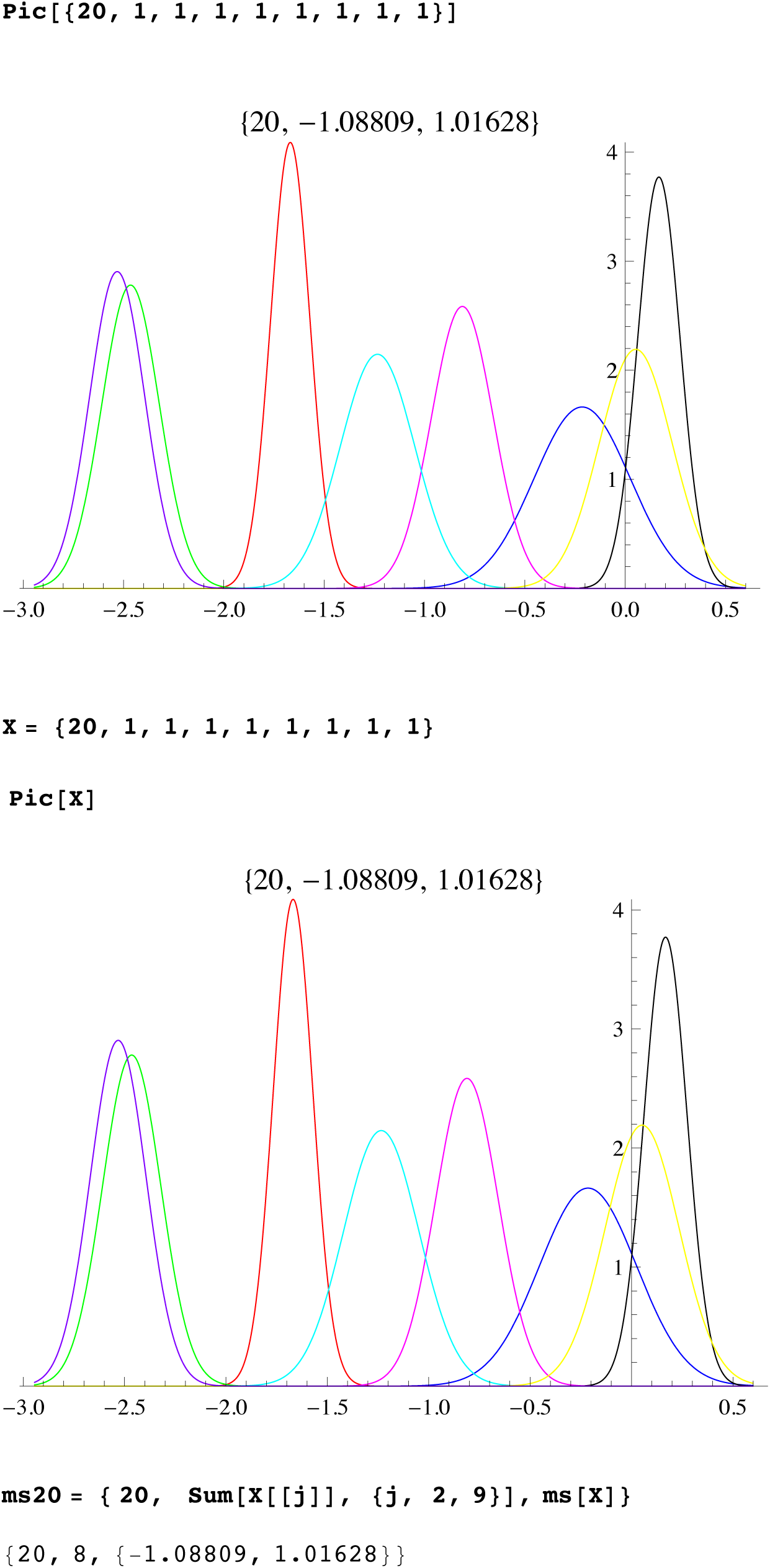

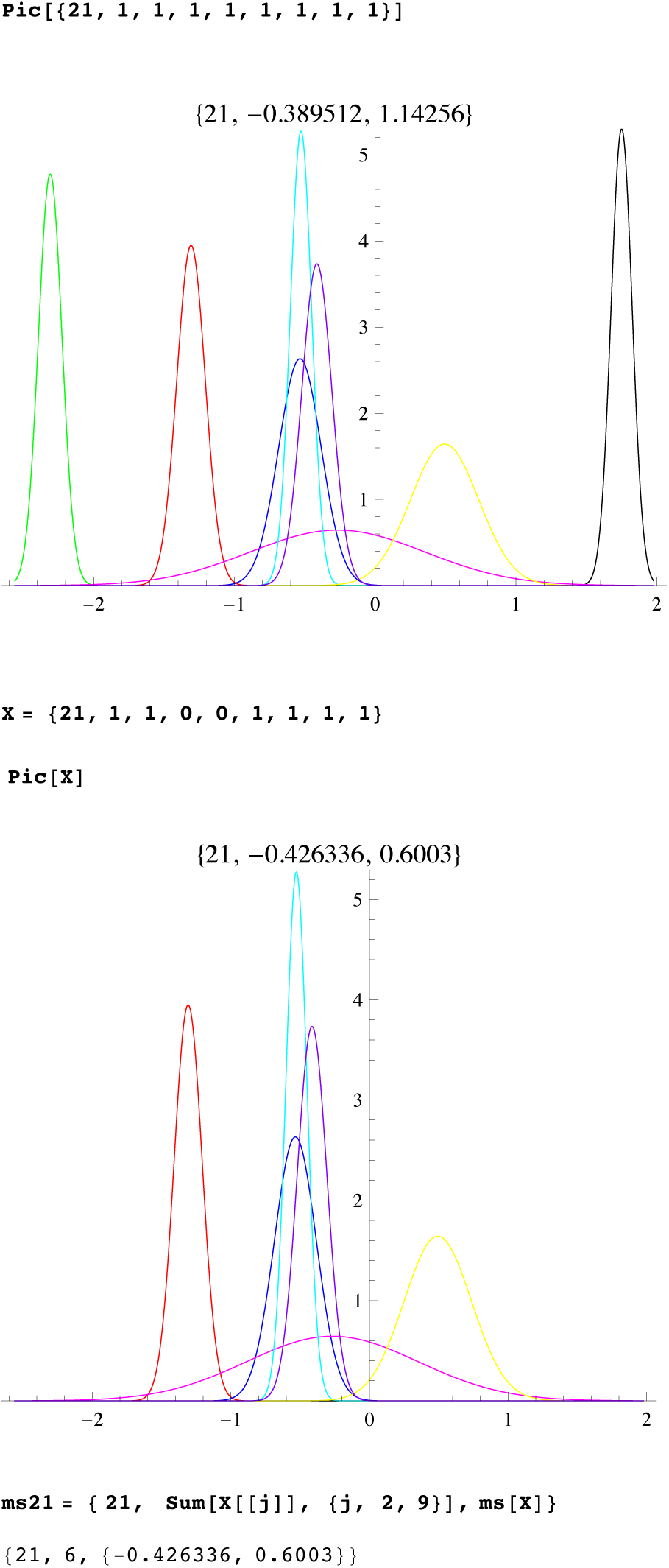

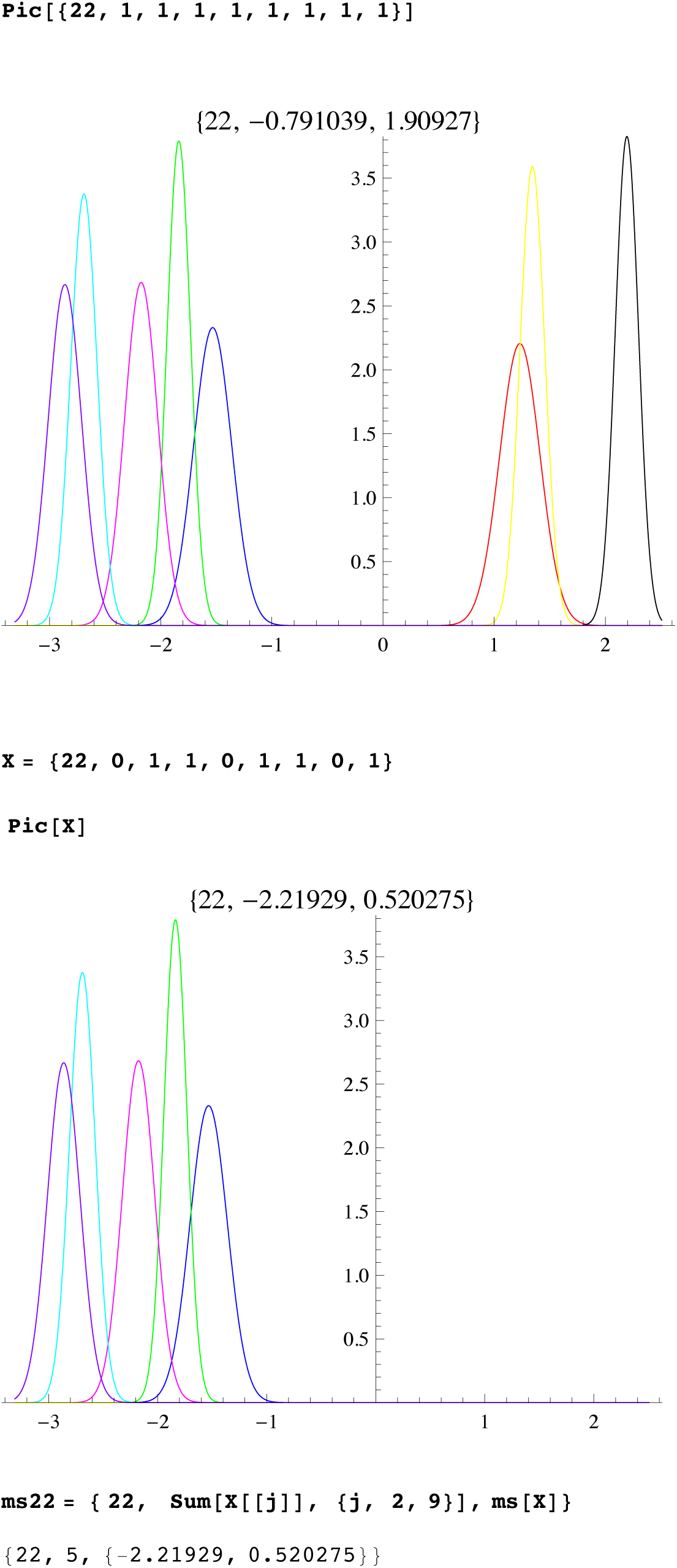

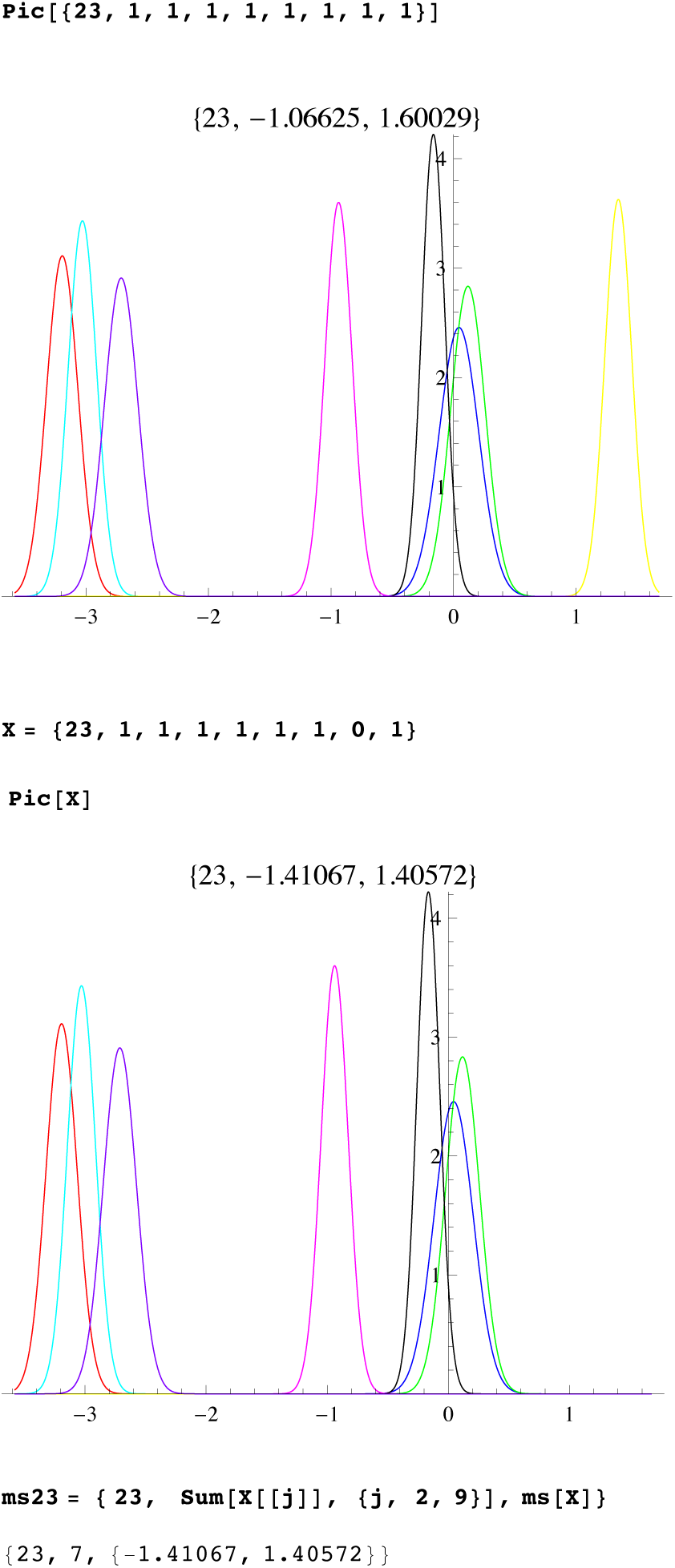

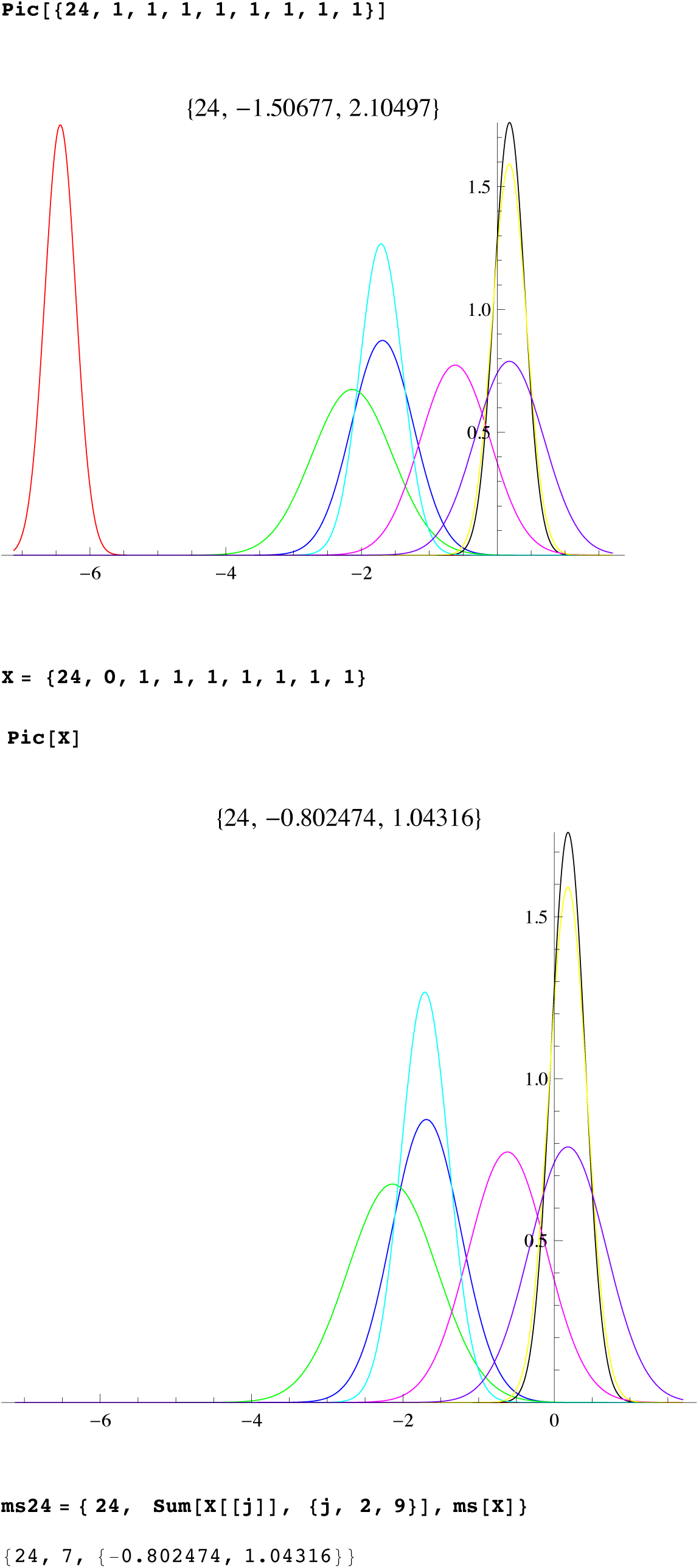

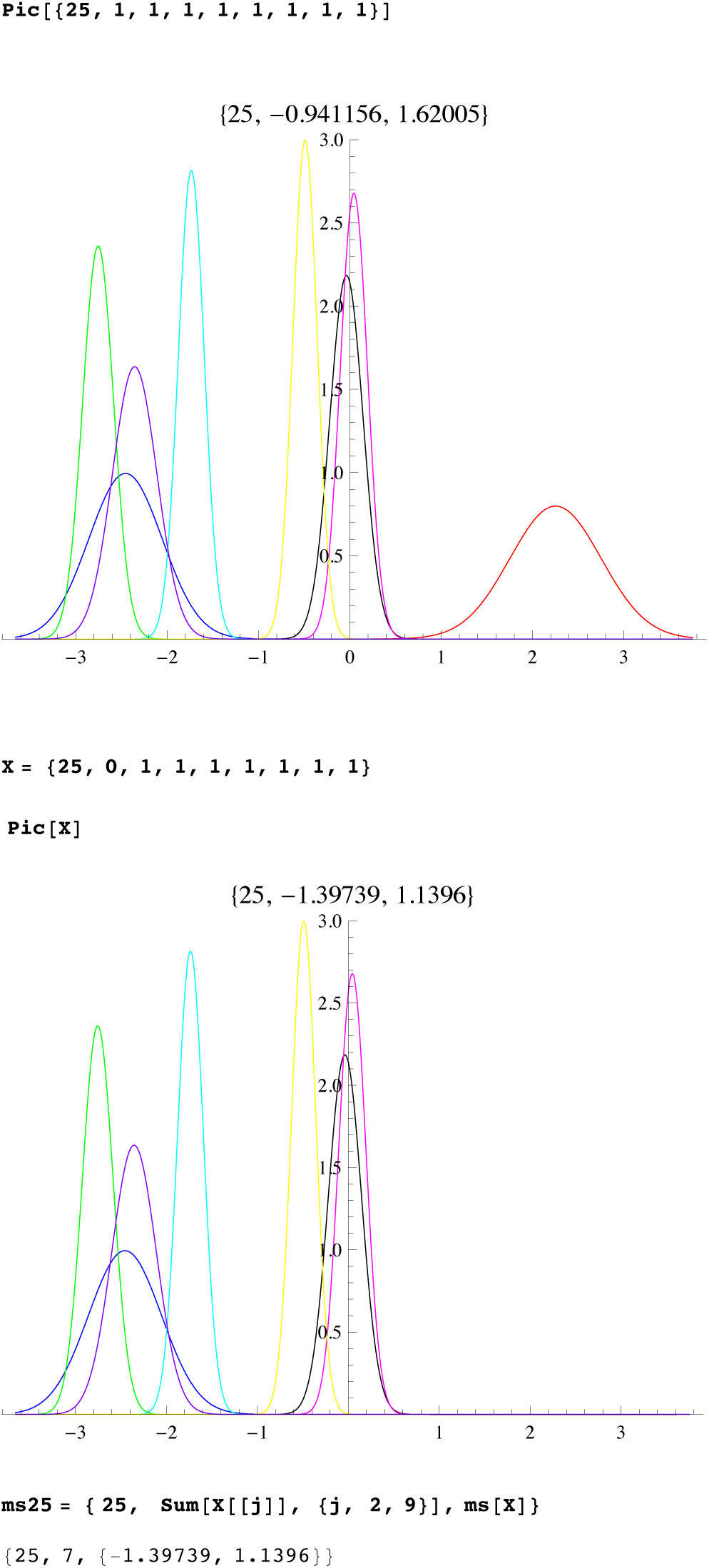

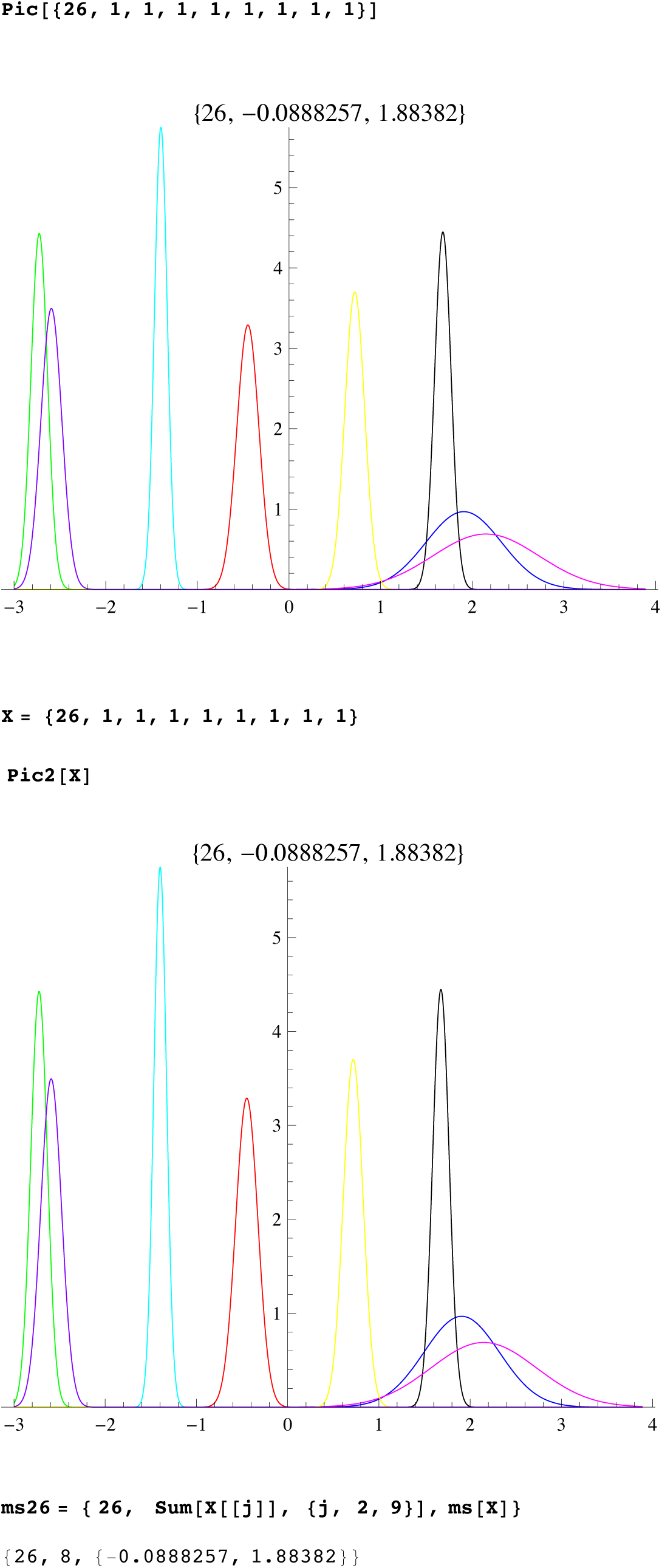

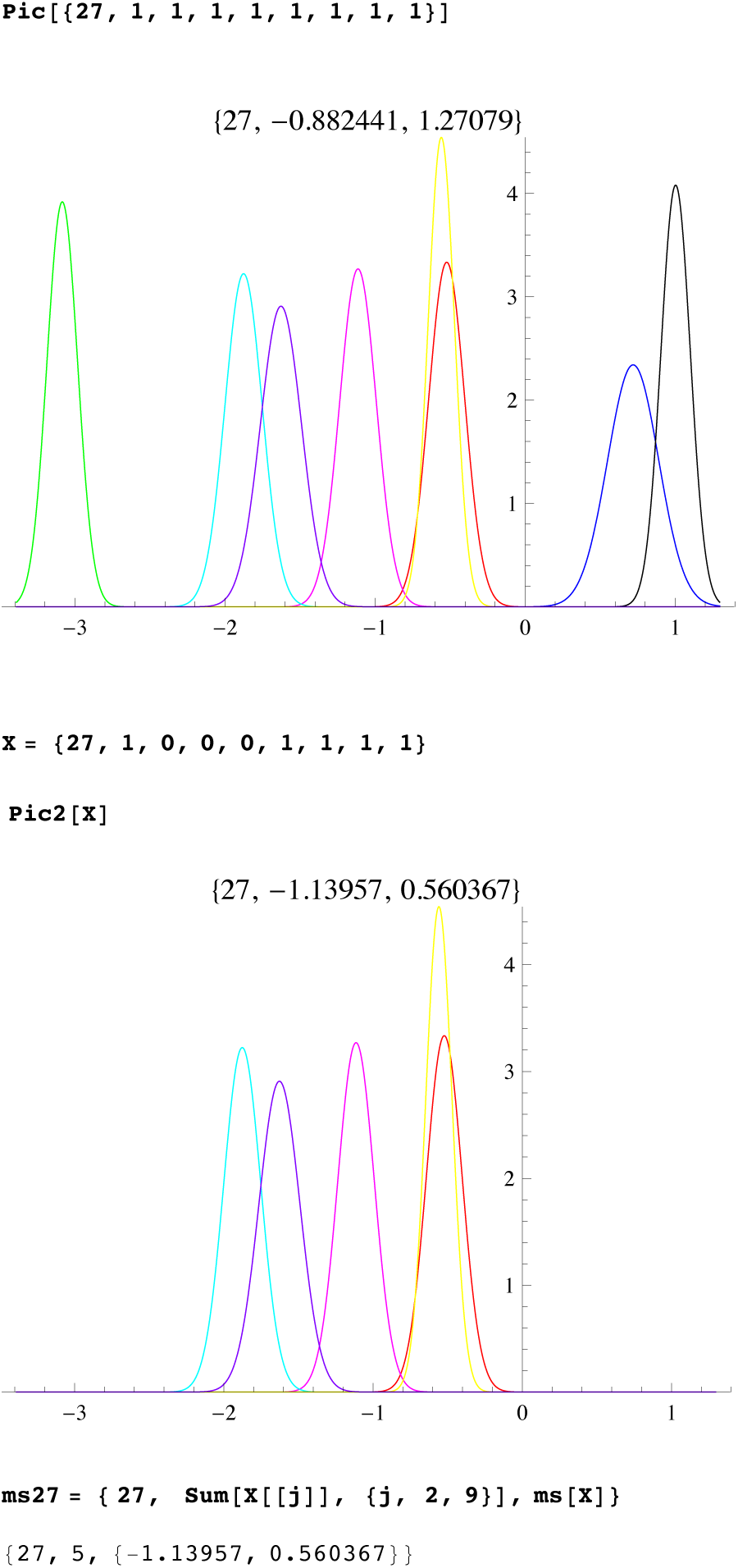

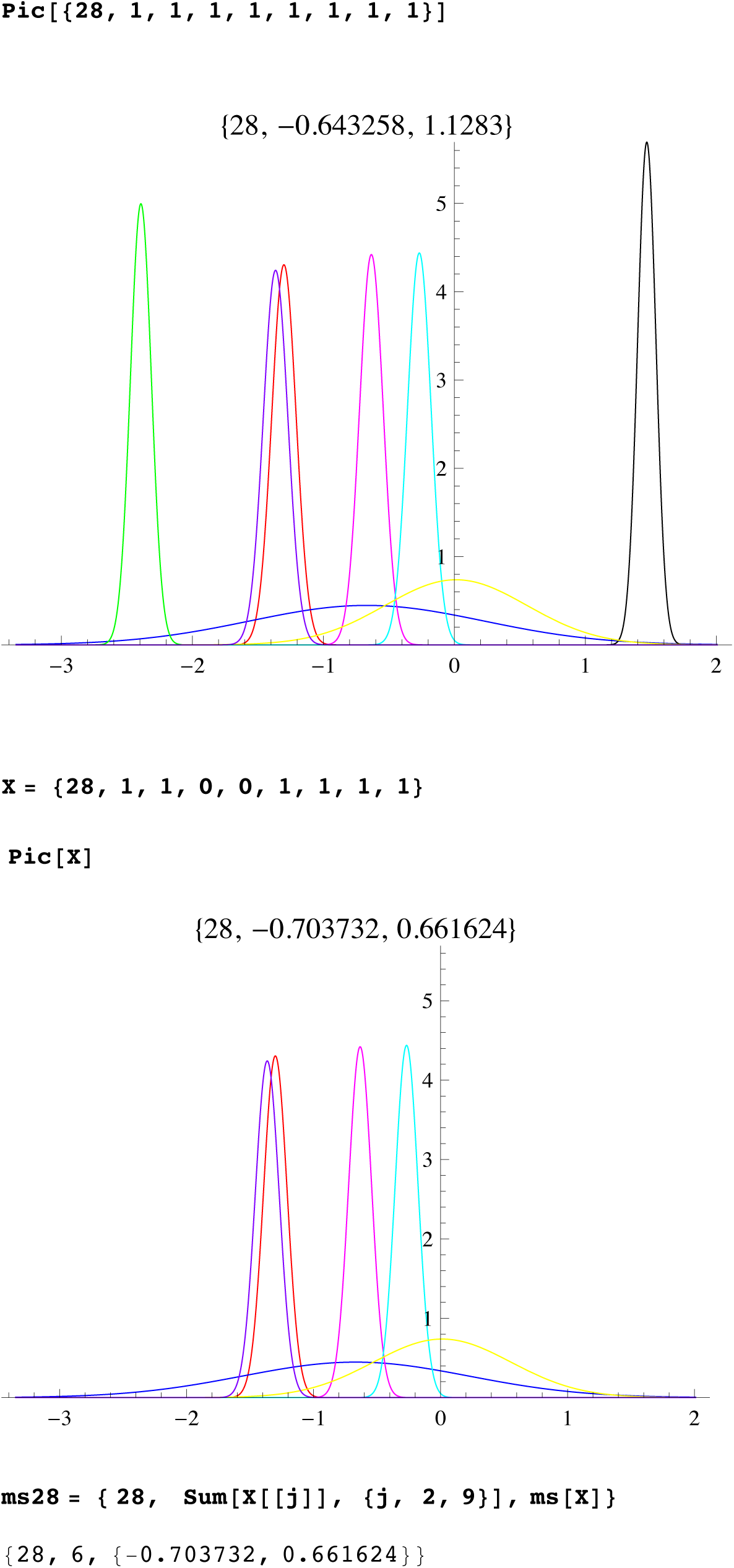

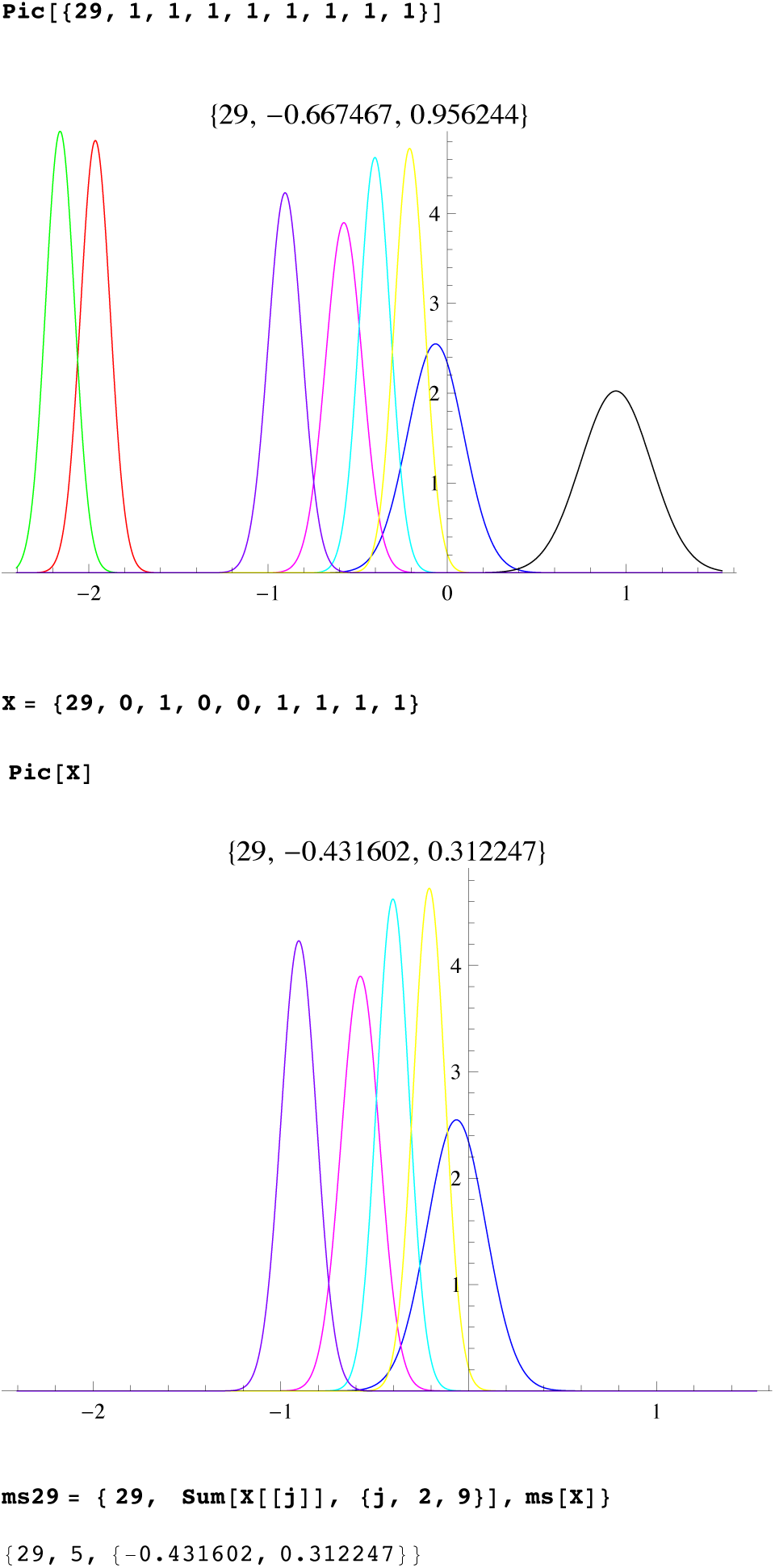

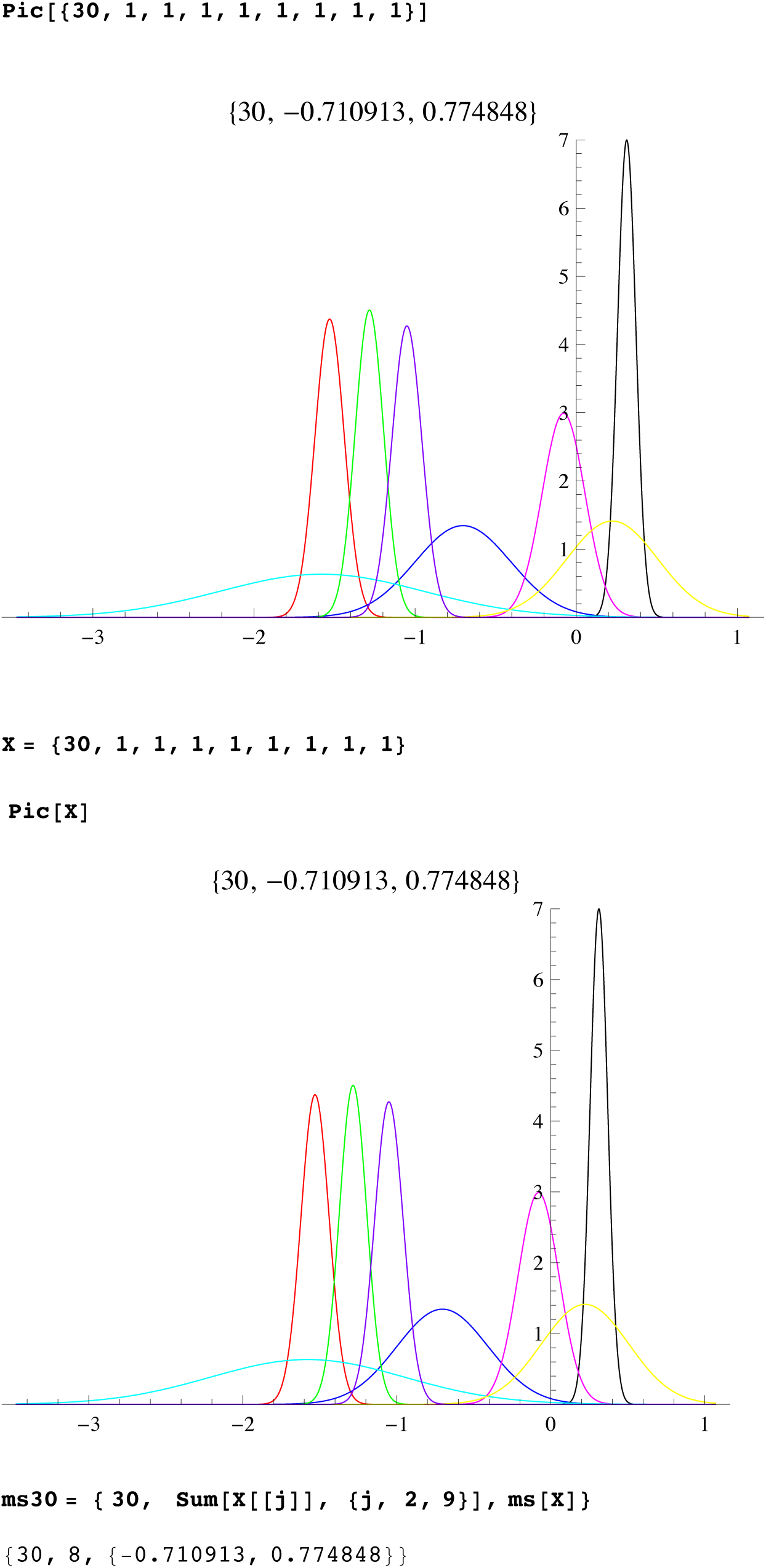

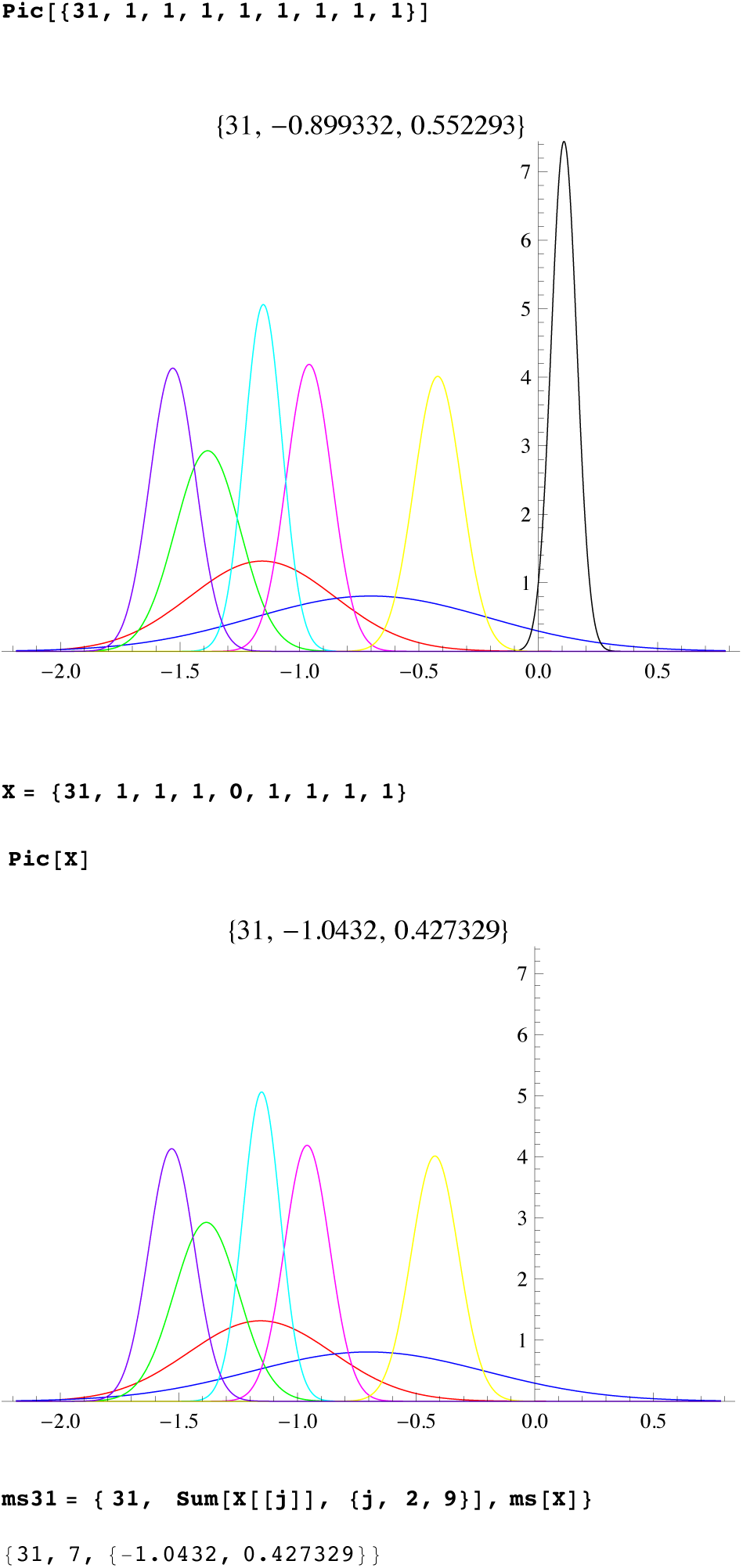

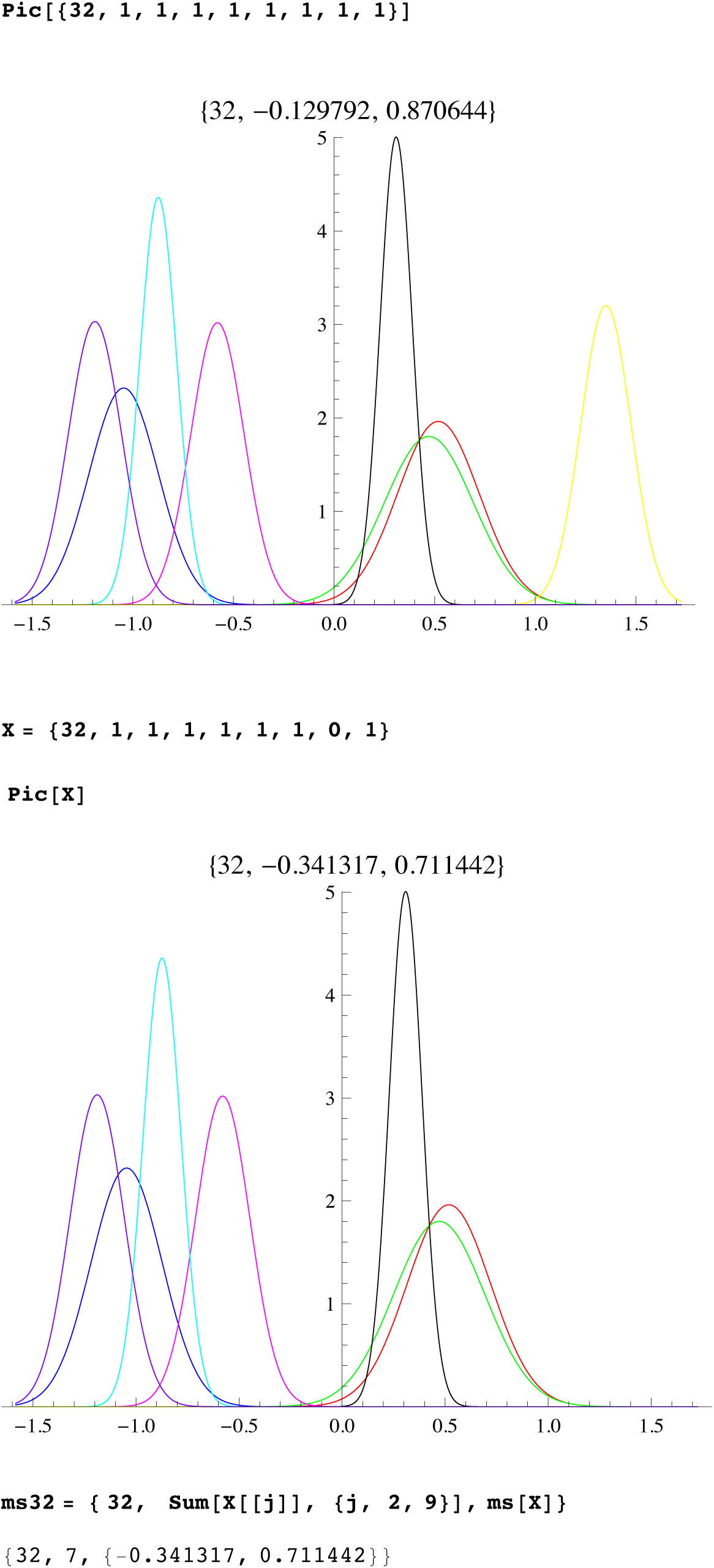

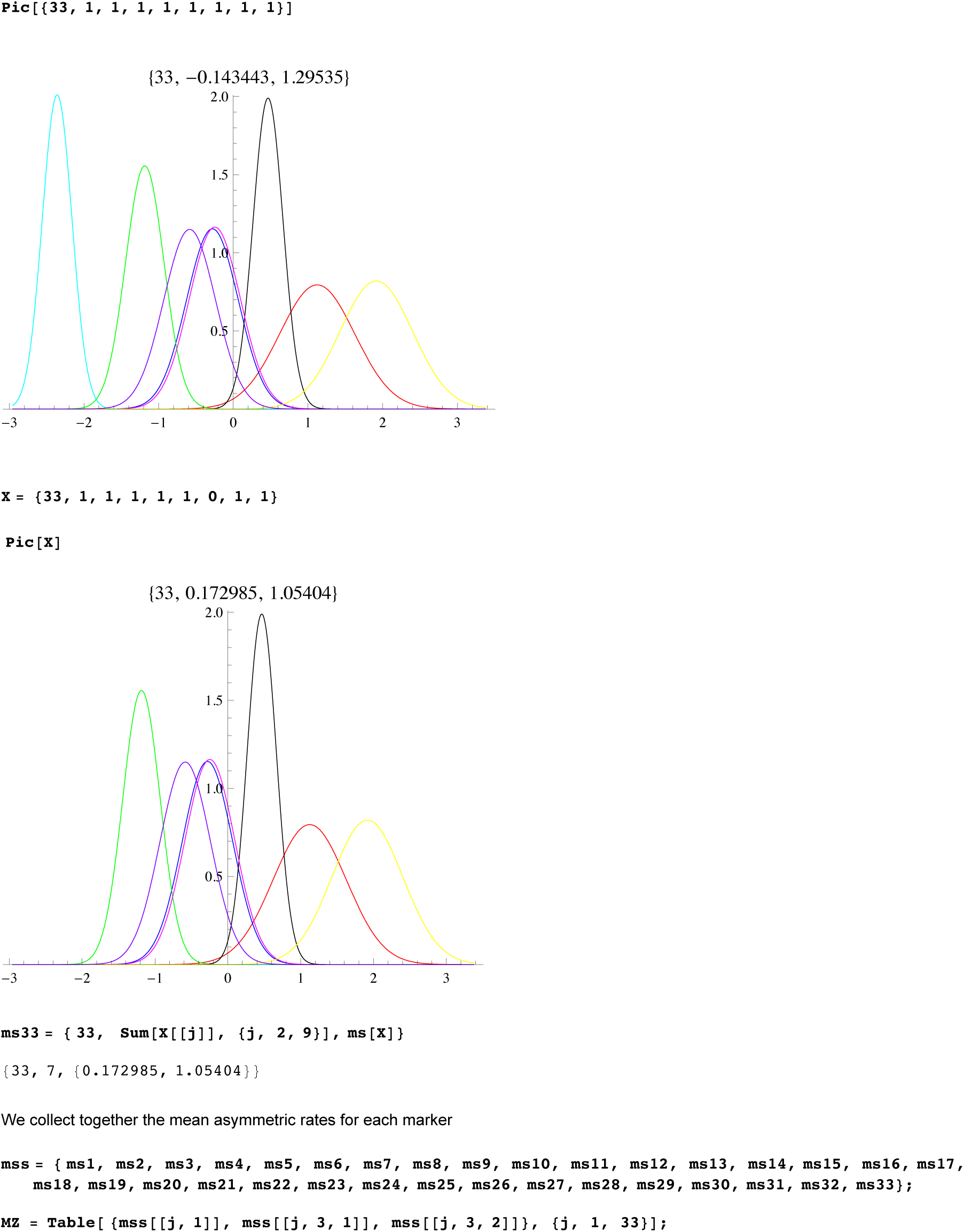

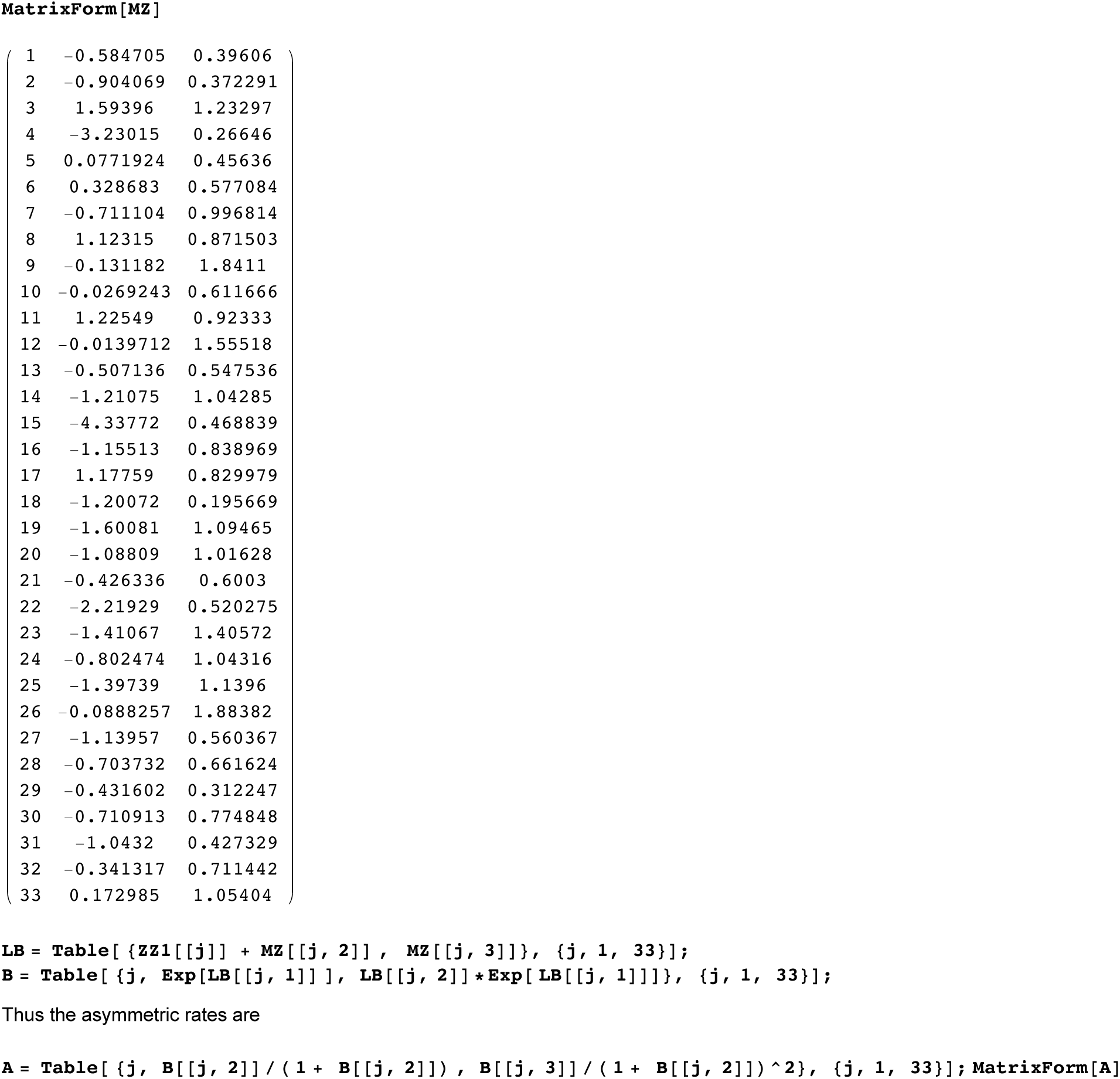

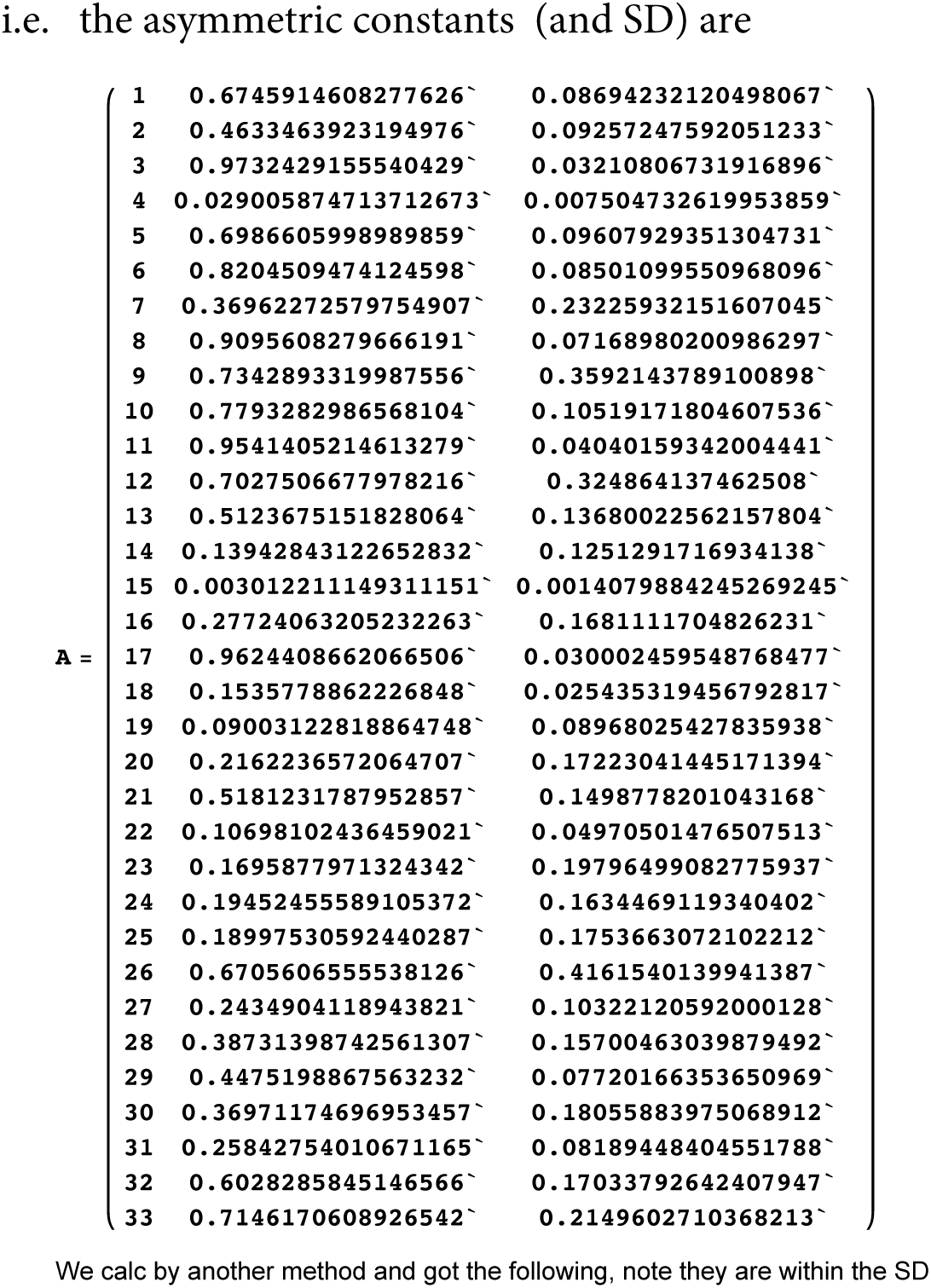

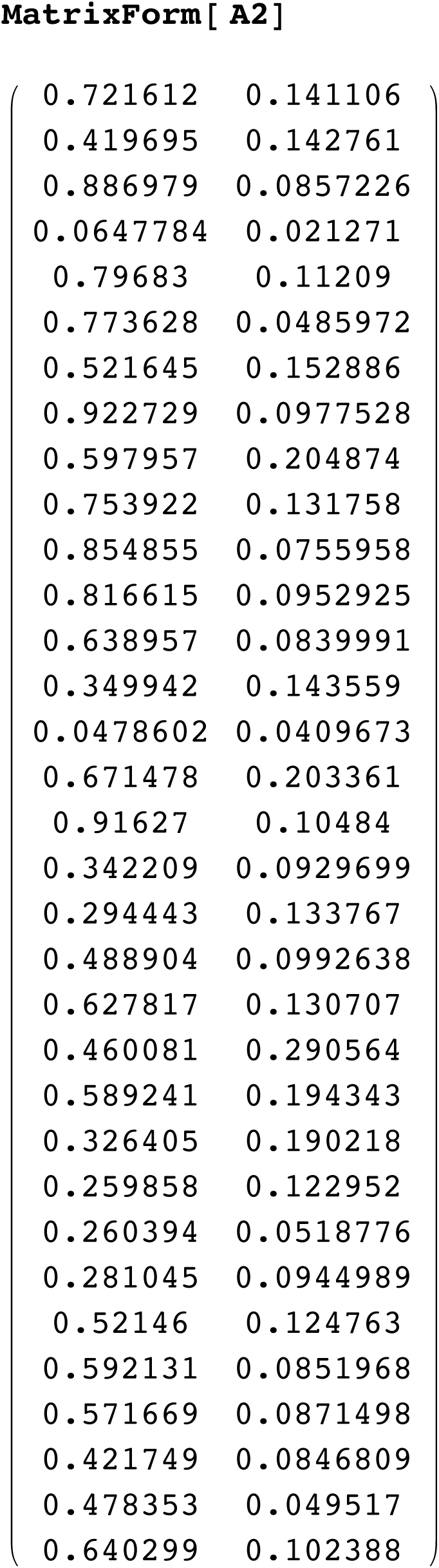

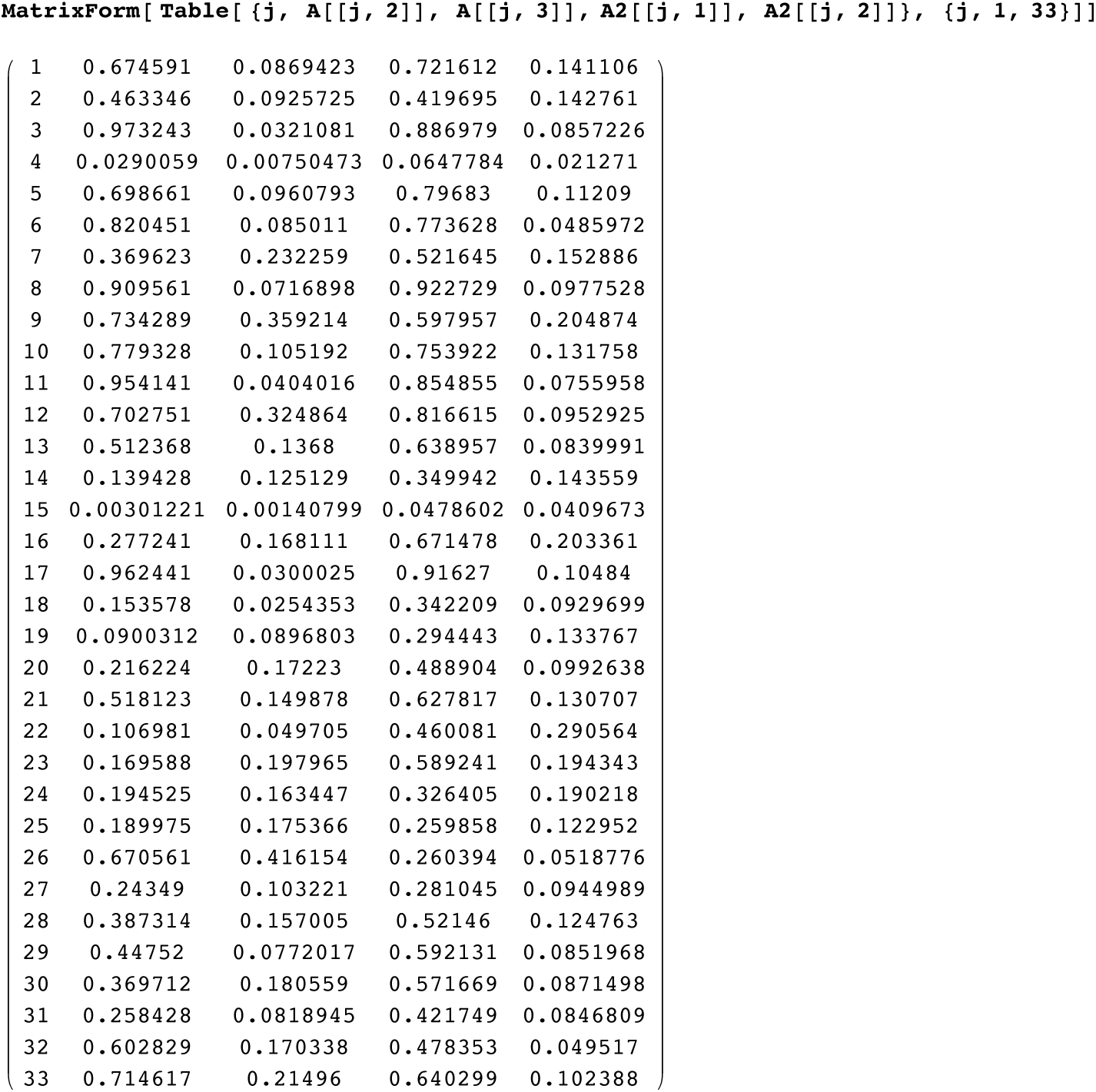

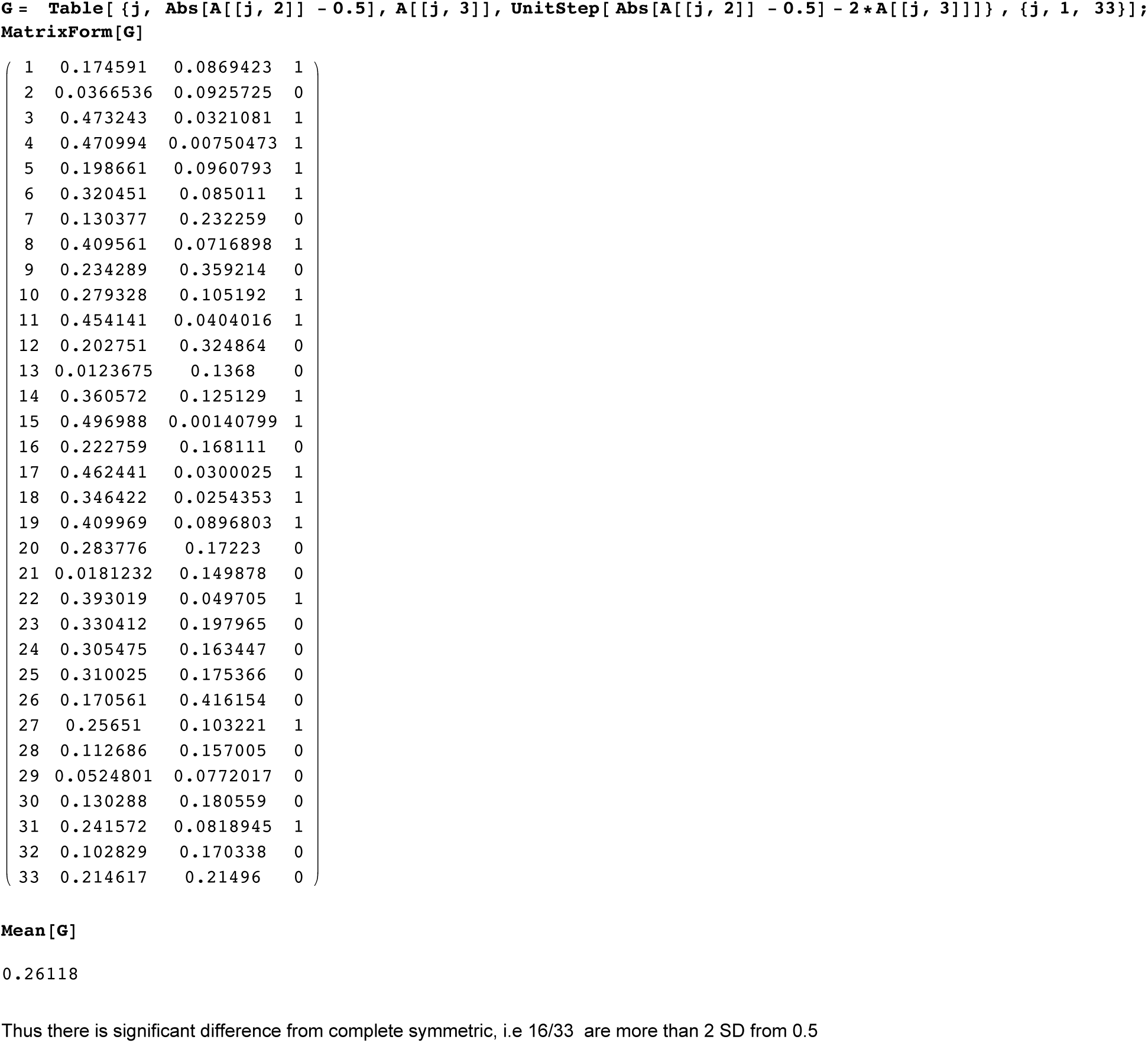

## SI 5 : Mathematica worksheet for computing mutation rates

We compute the mutation rates using SNP G2a3, R1b1a2, R1a1, I1, L21,U106, J2, P312. First we download this data for these SNP

**δδ;**

The asymmetric rates are entered. We only use the asymmetric rates and its log ratio LB

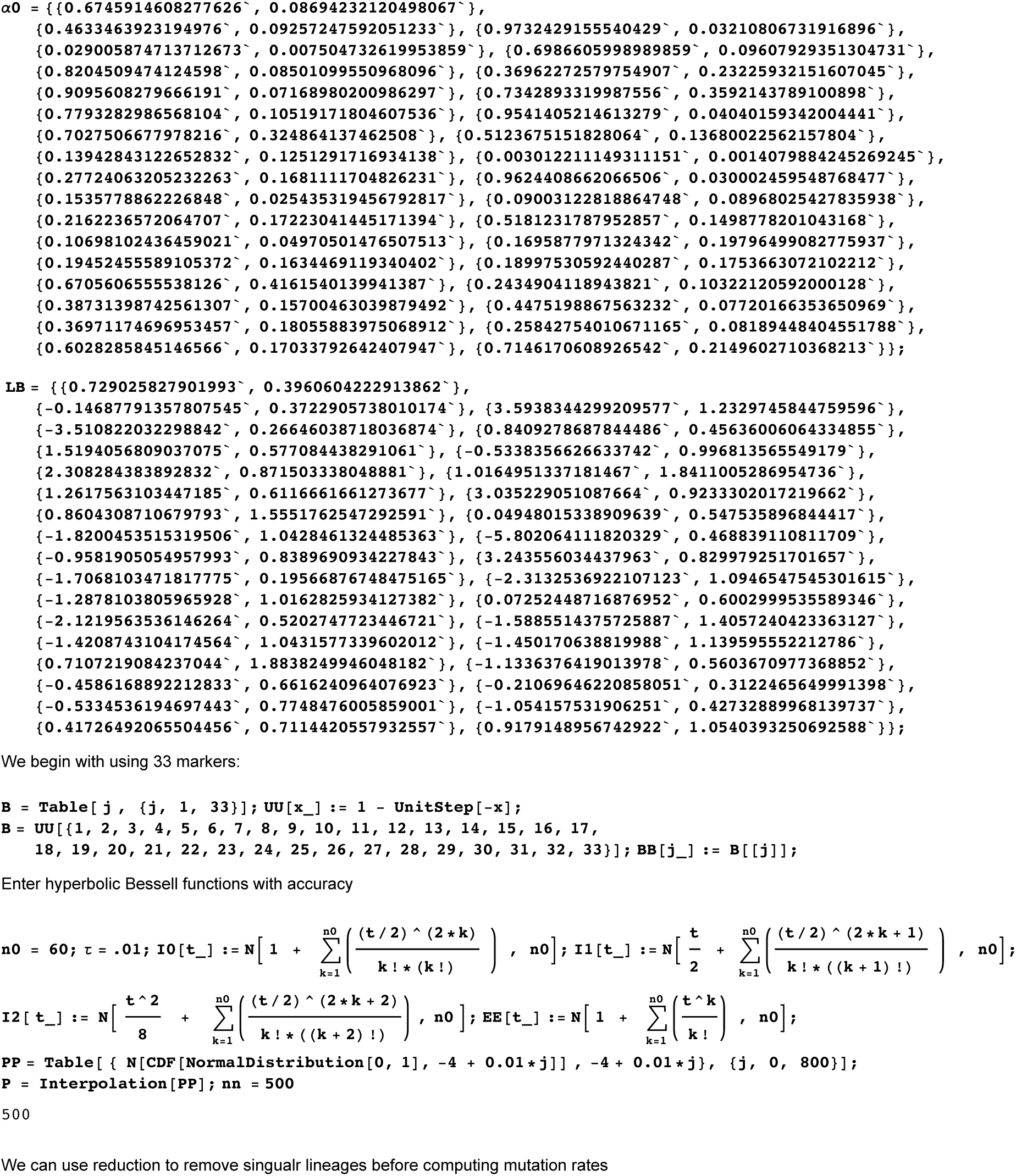

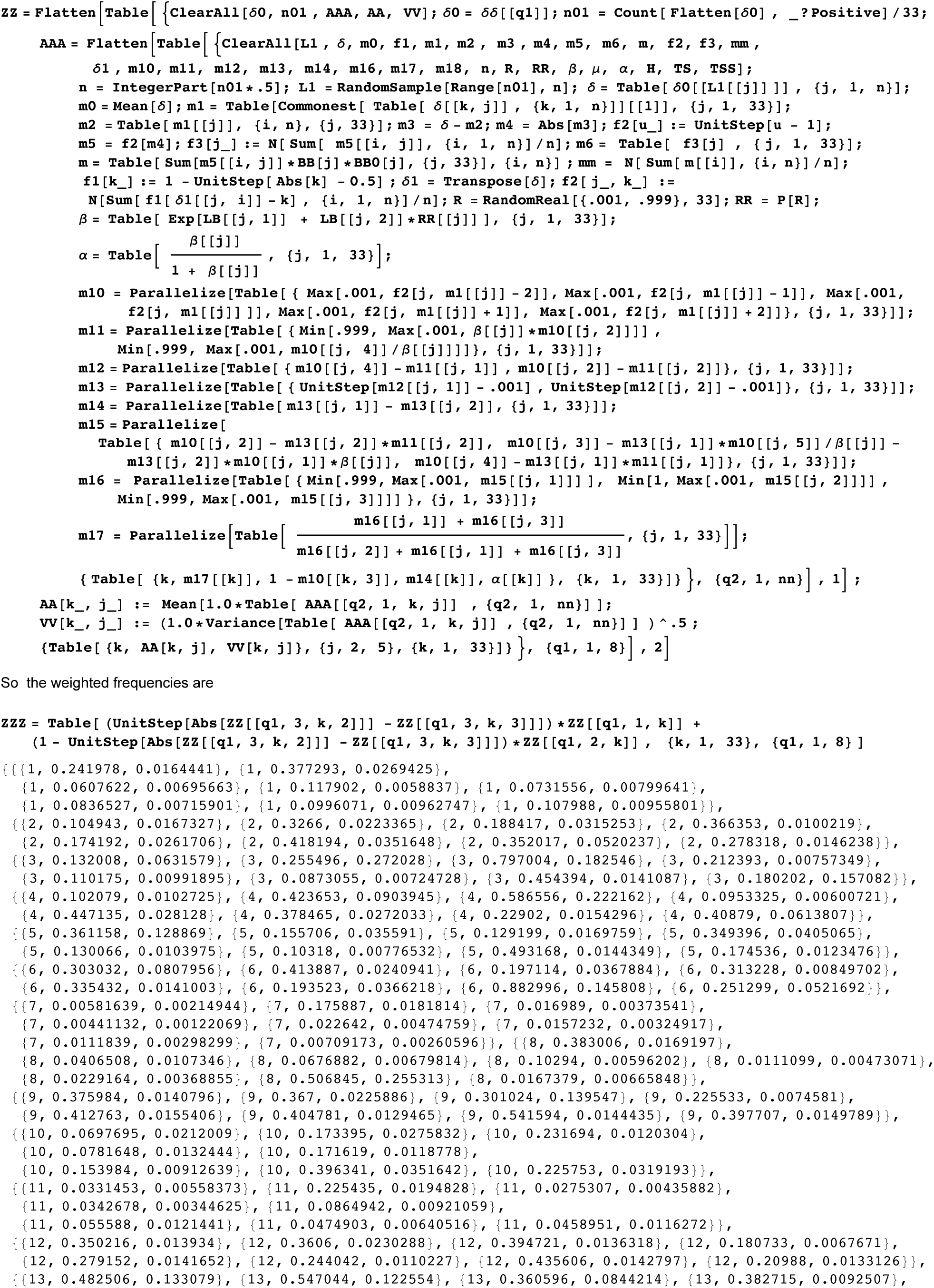

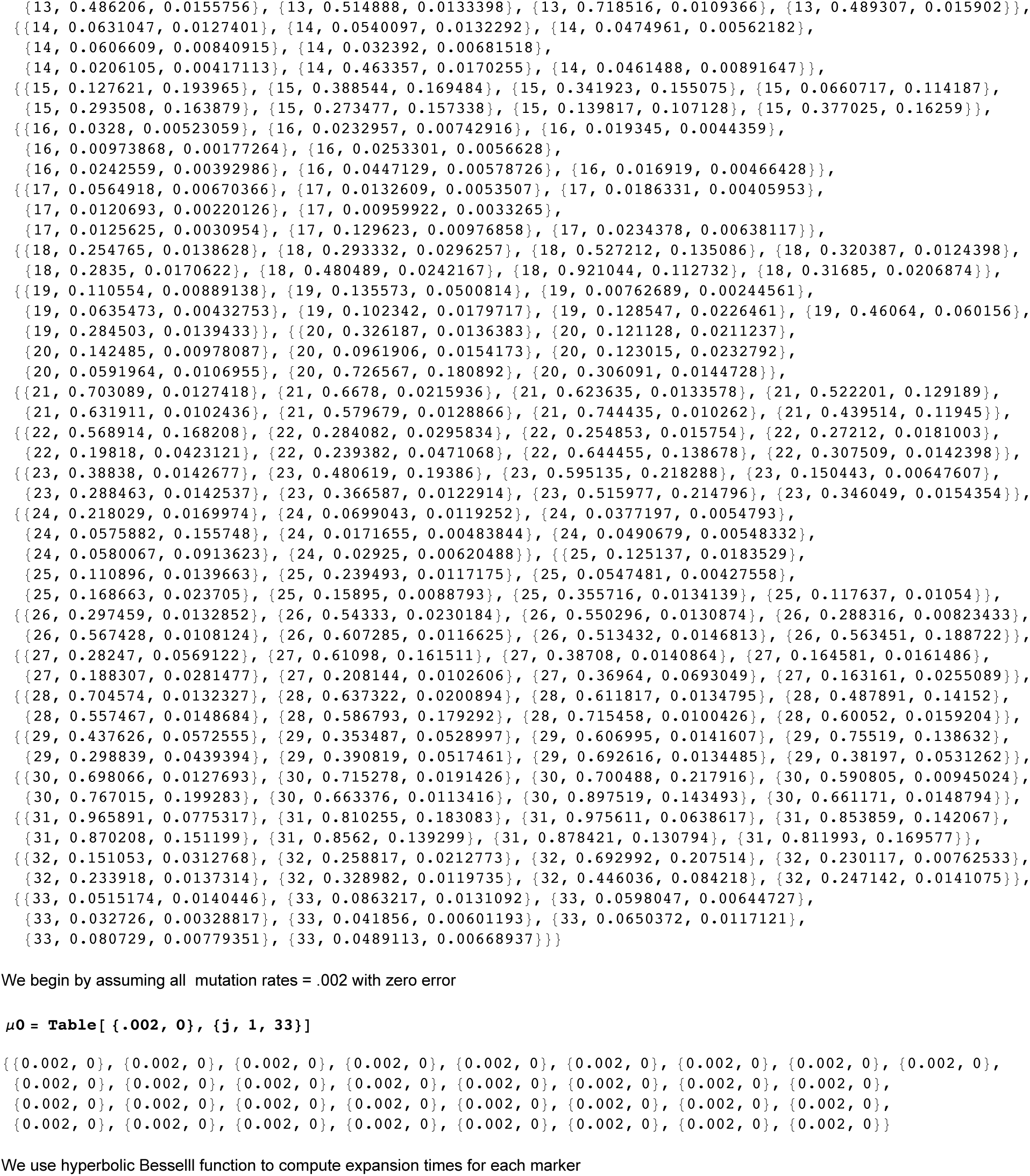

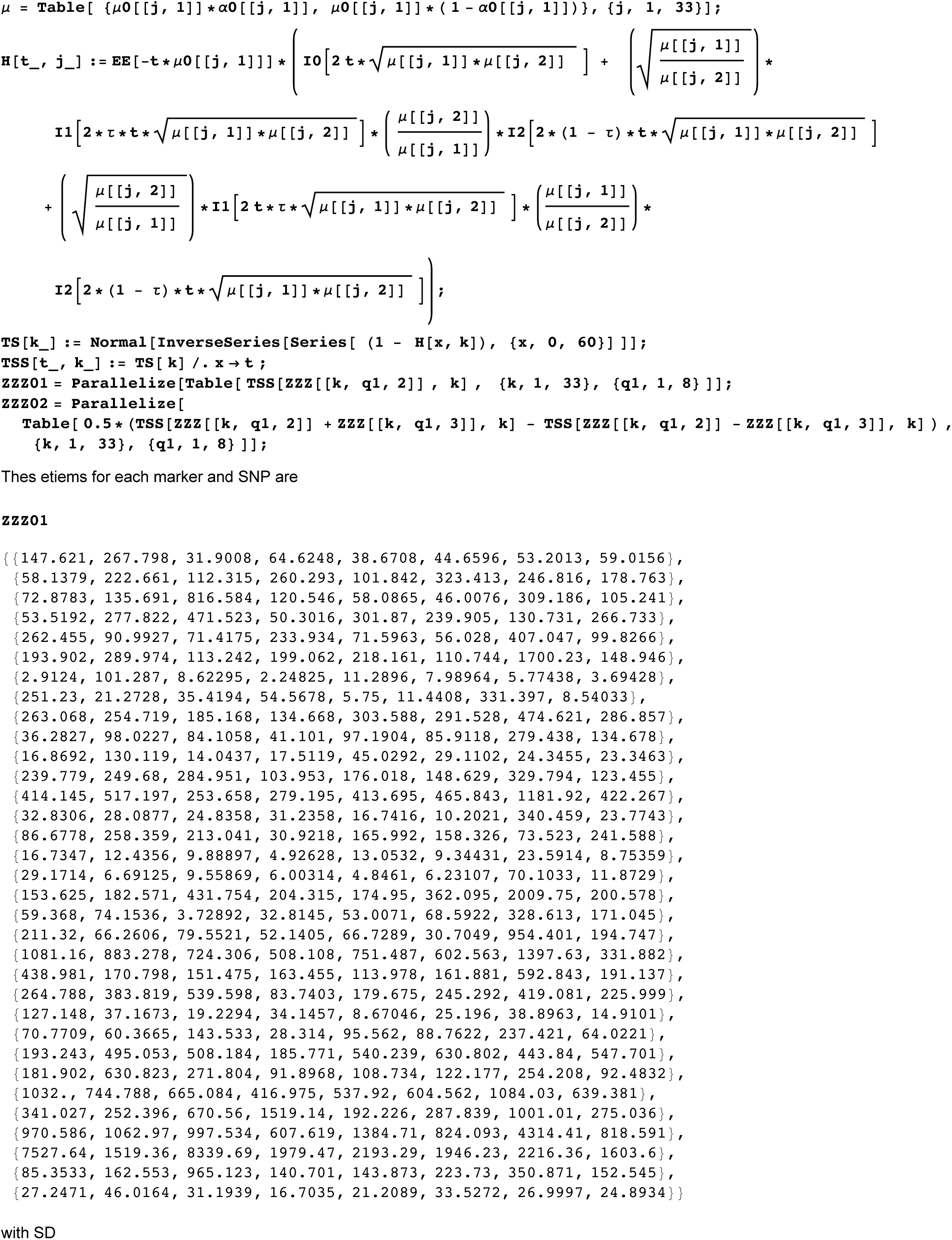

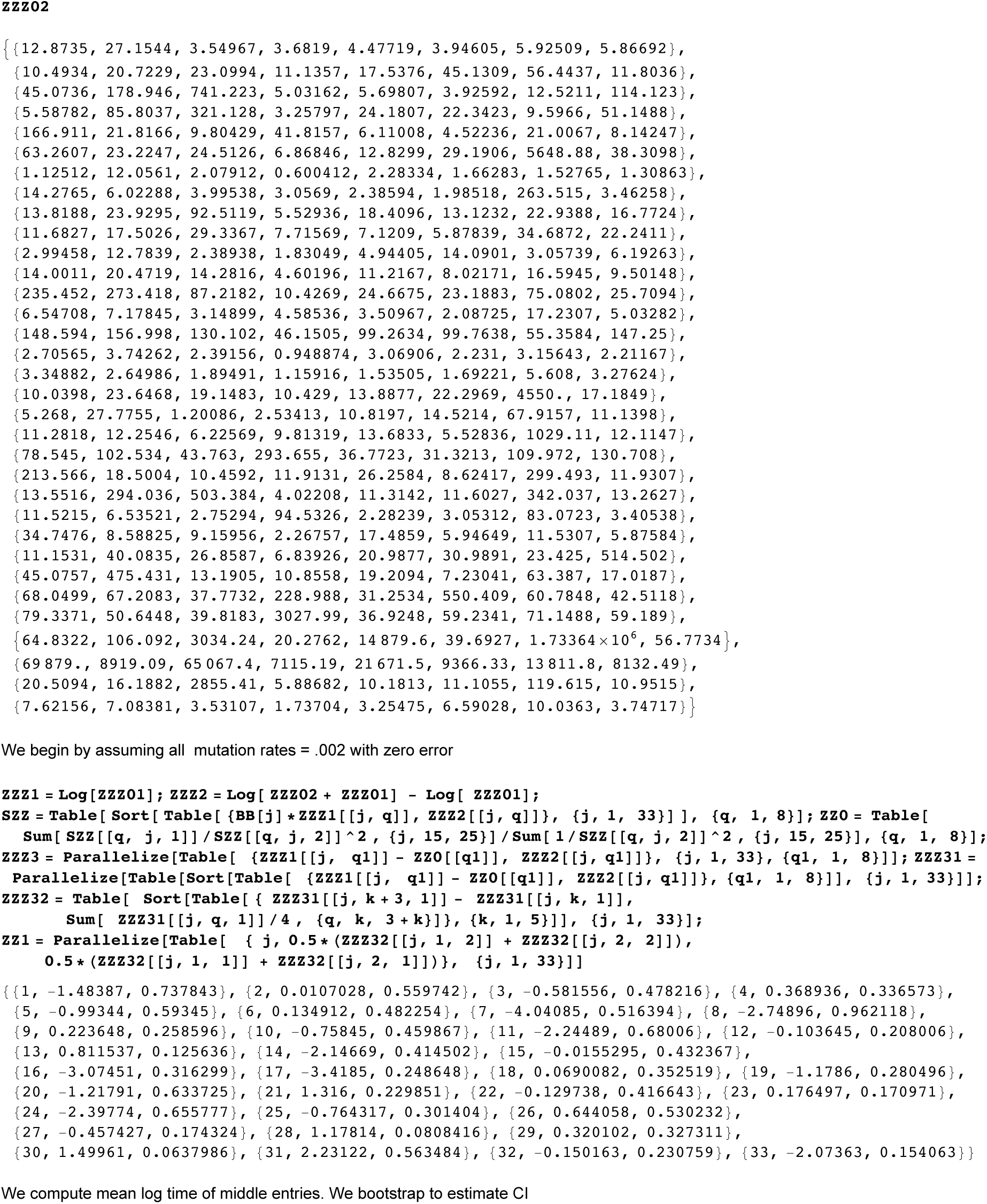

## SI 6: TMRCA for G2a2, R1b1a2, R1a1a, I1, L21, U106, J2, P312

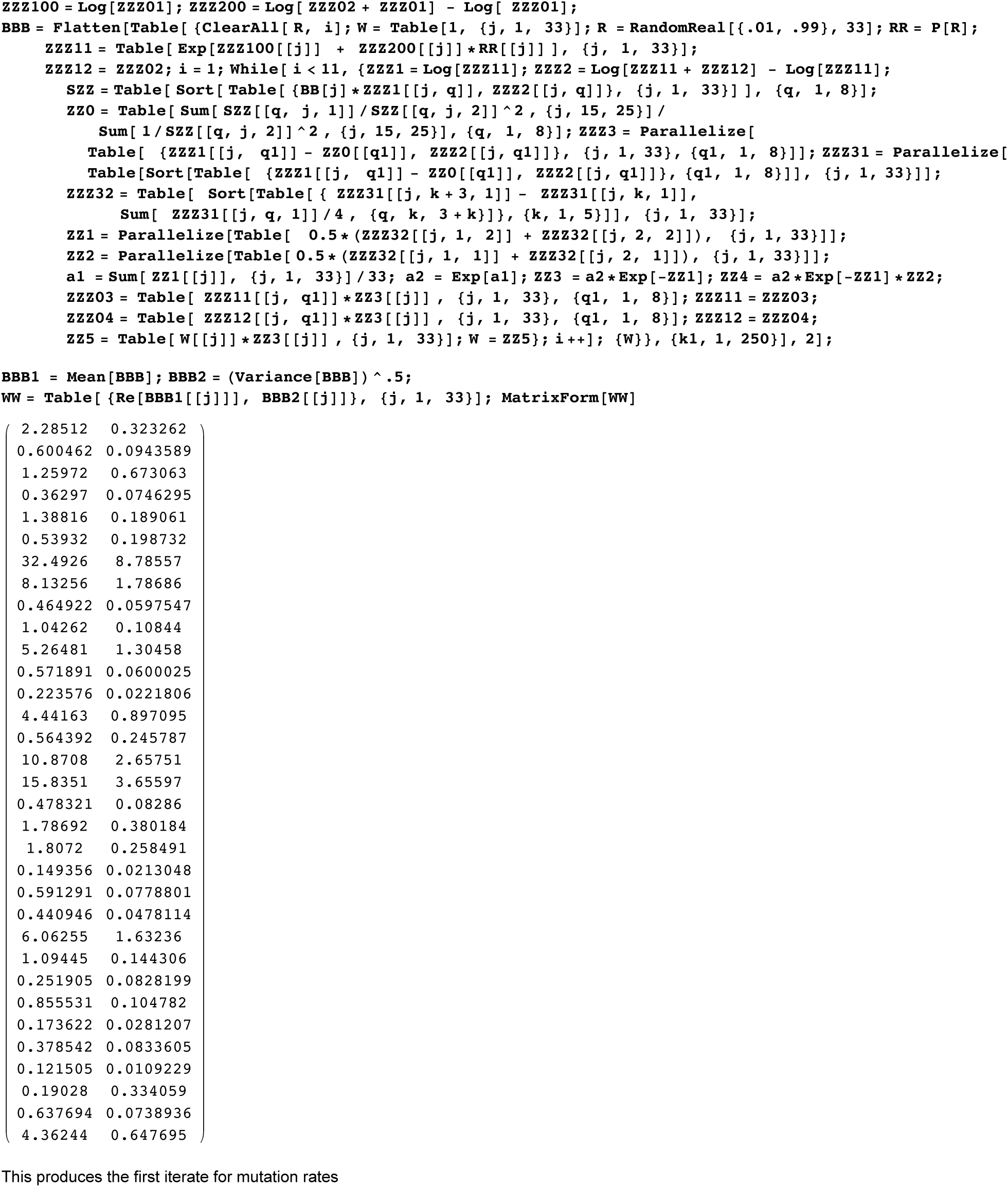

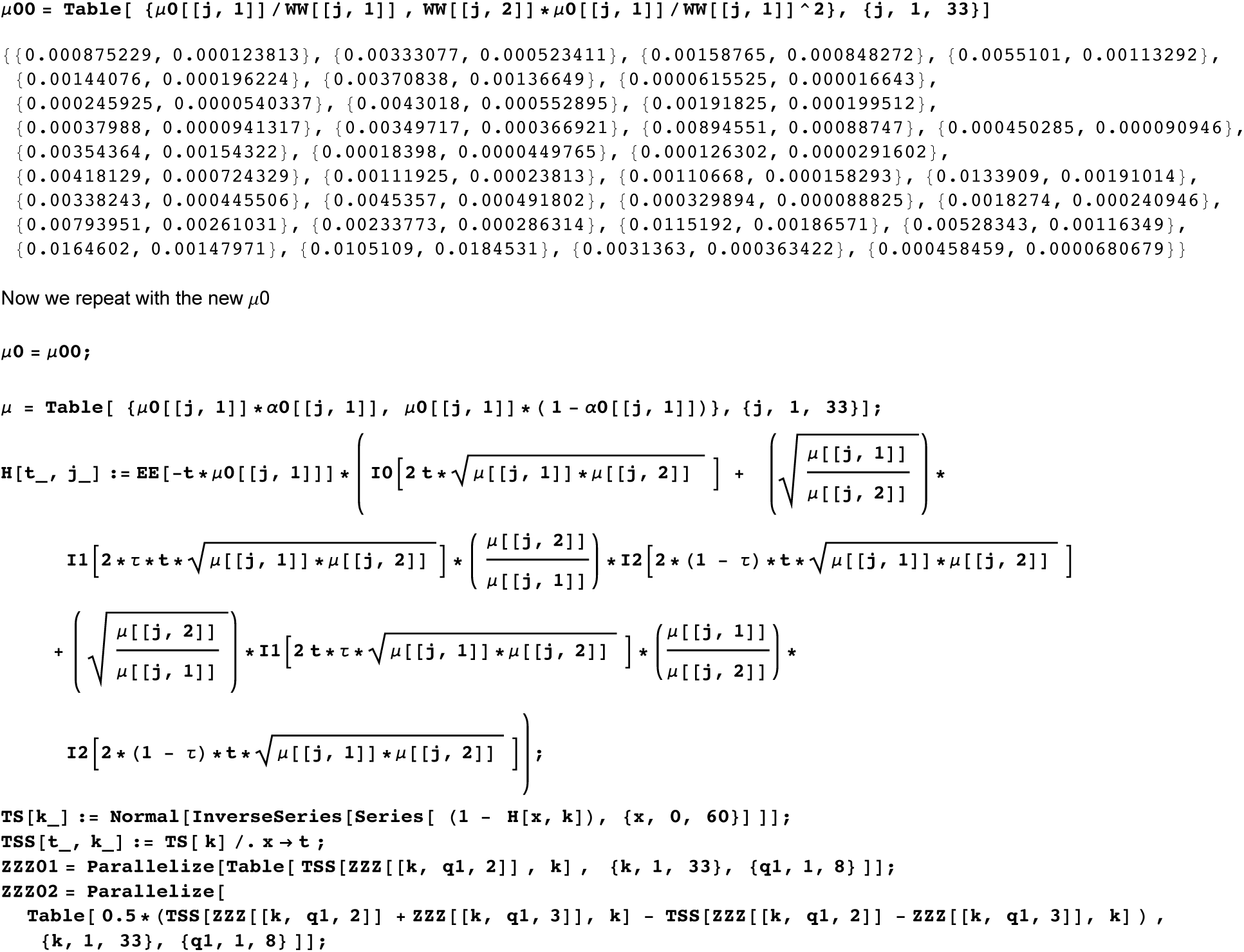

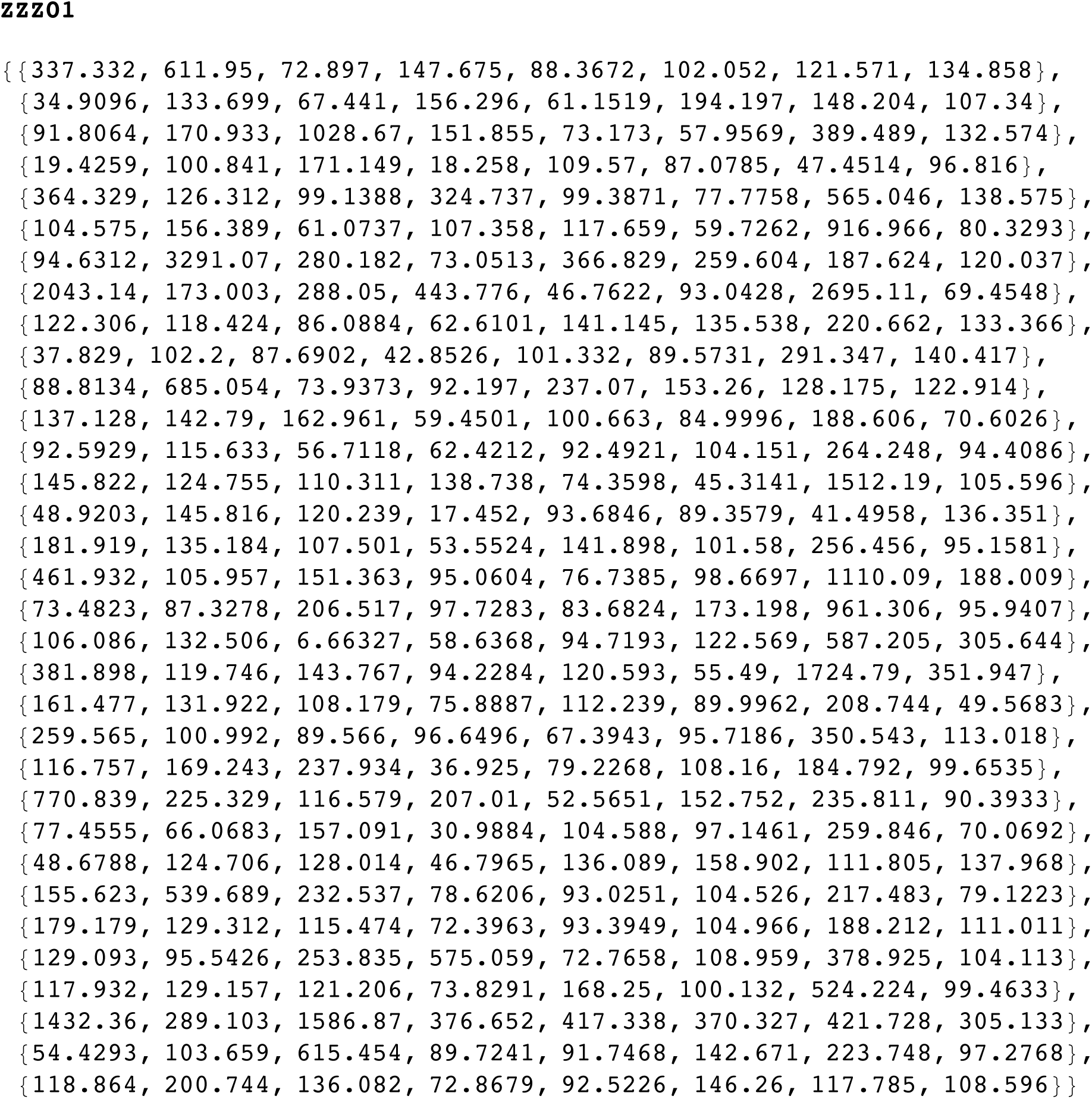

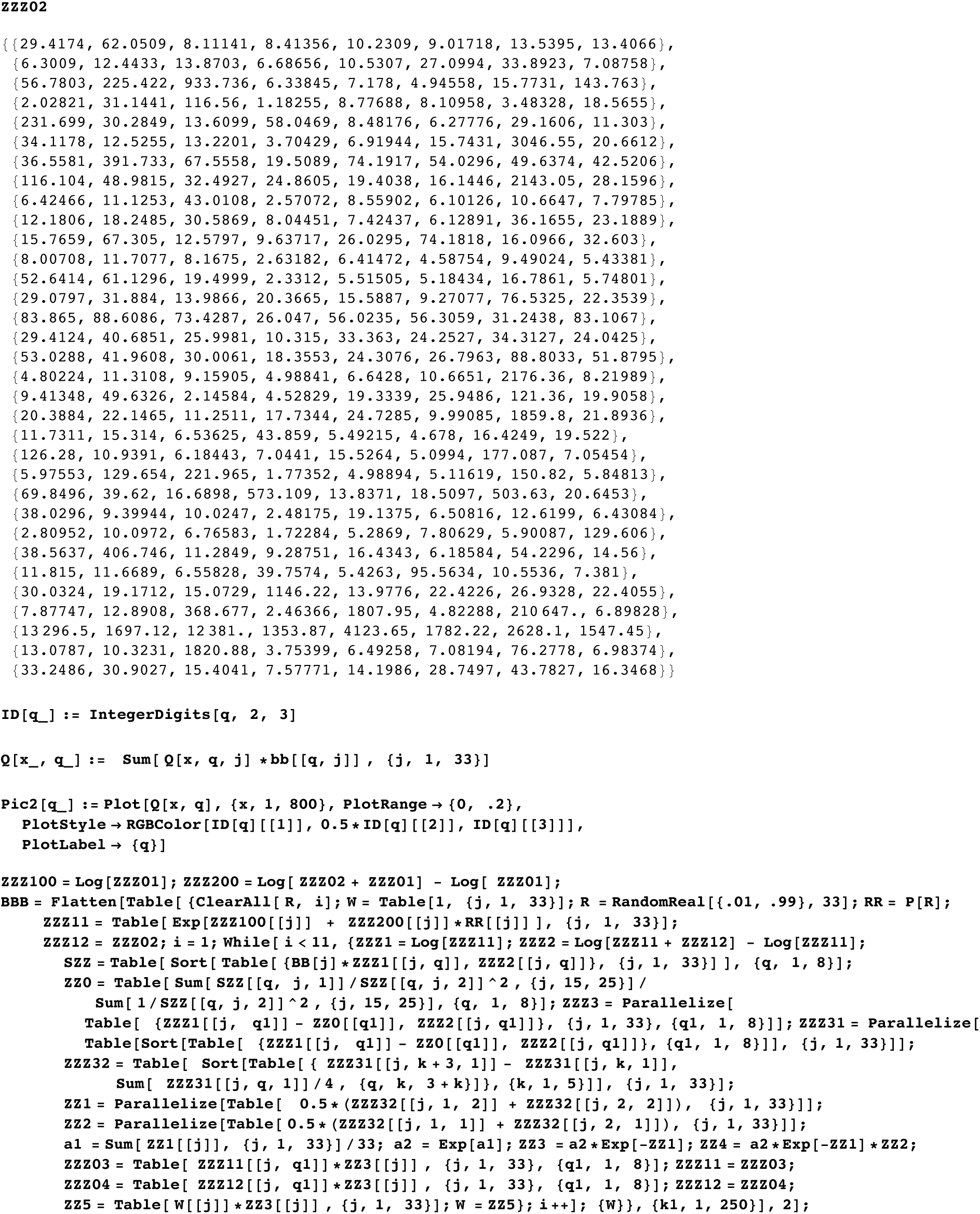

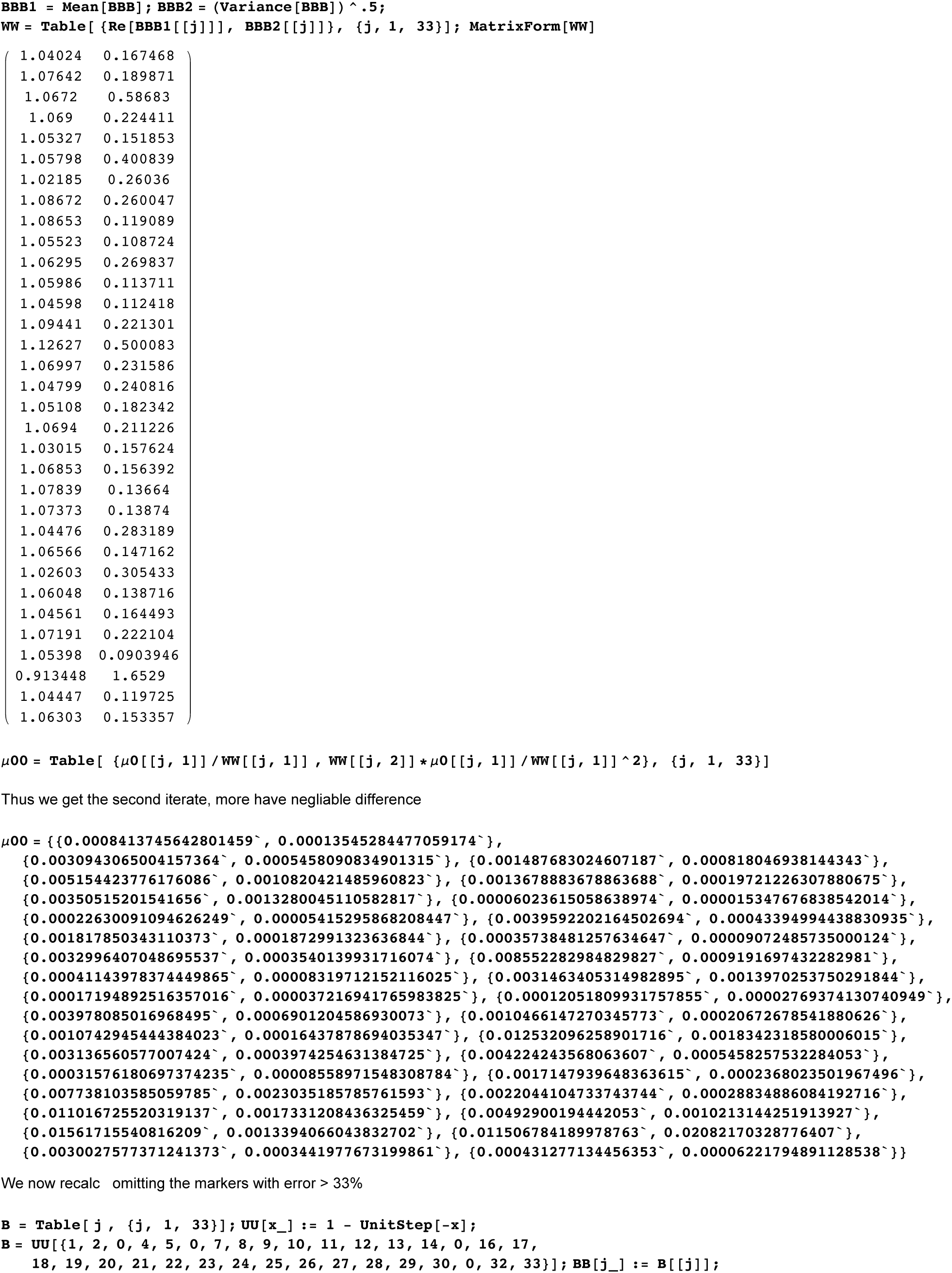

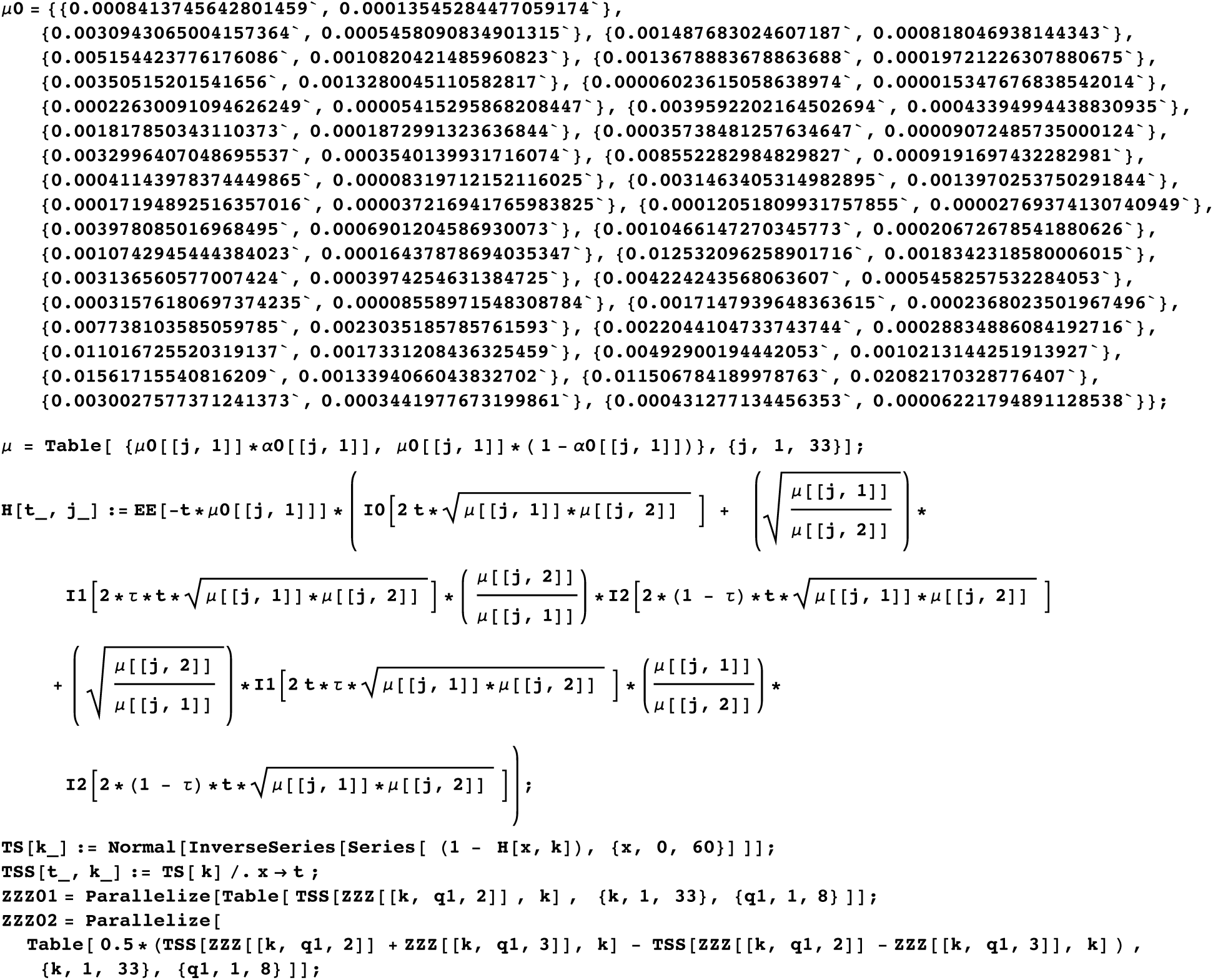

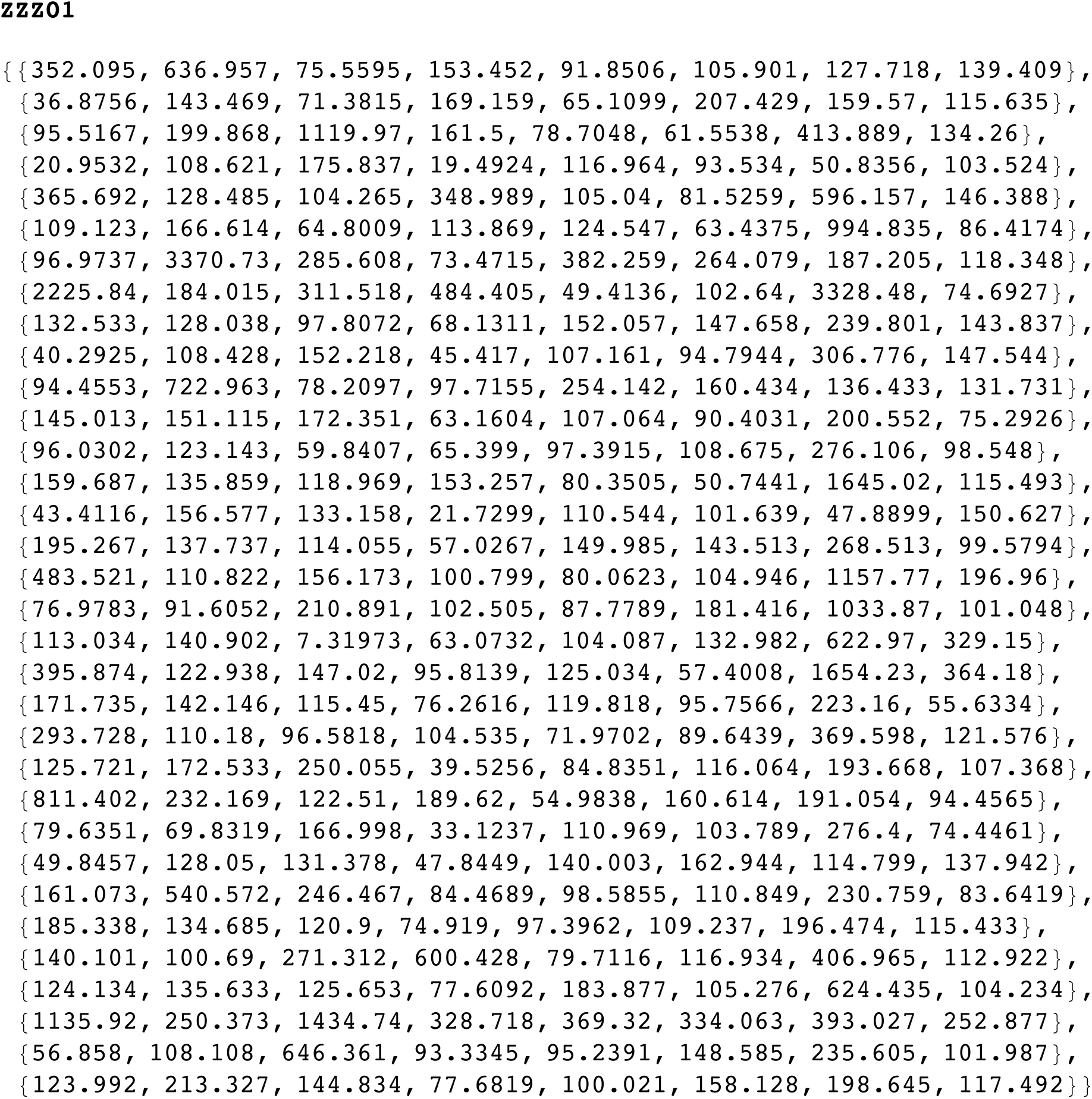

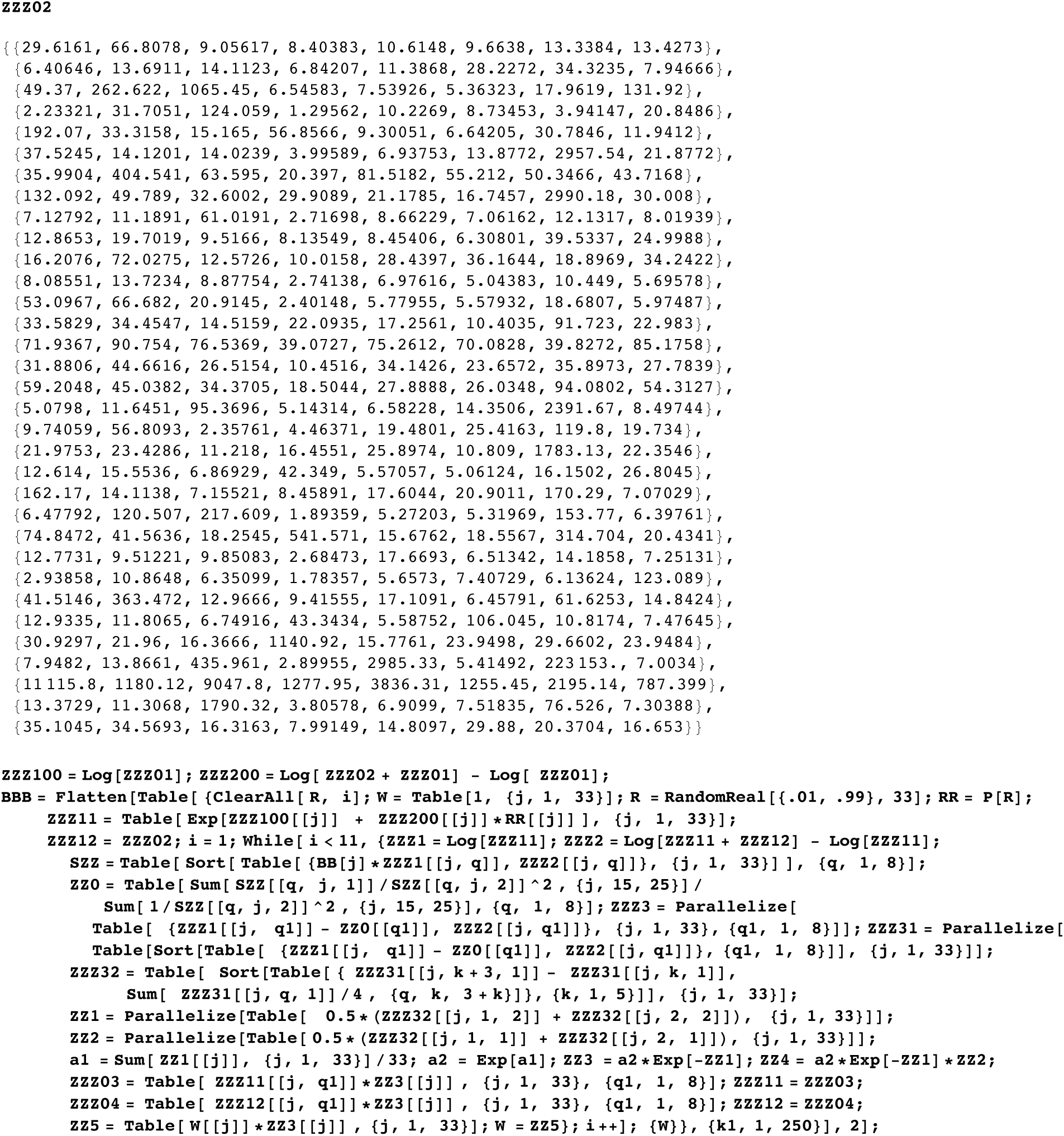

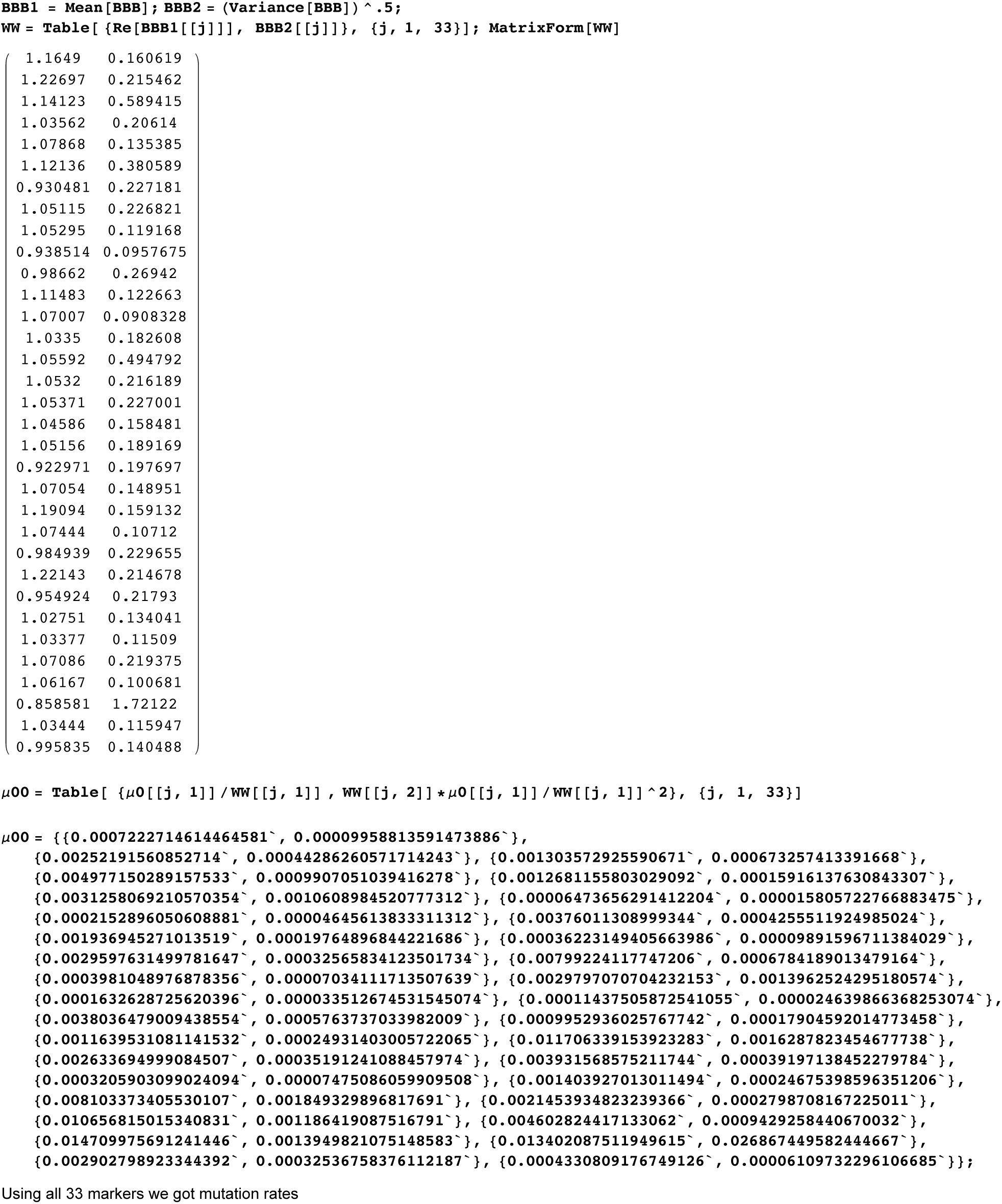

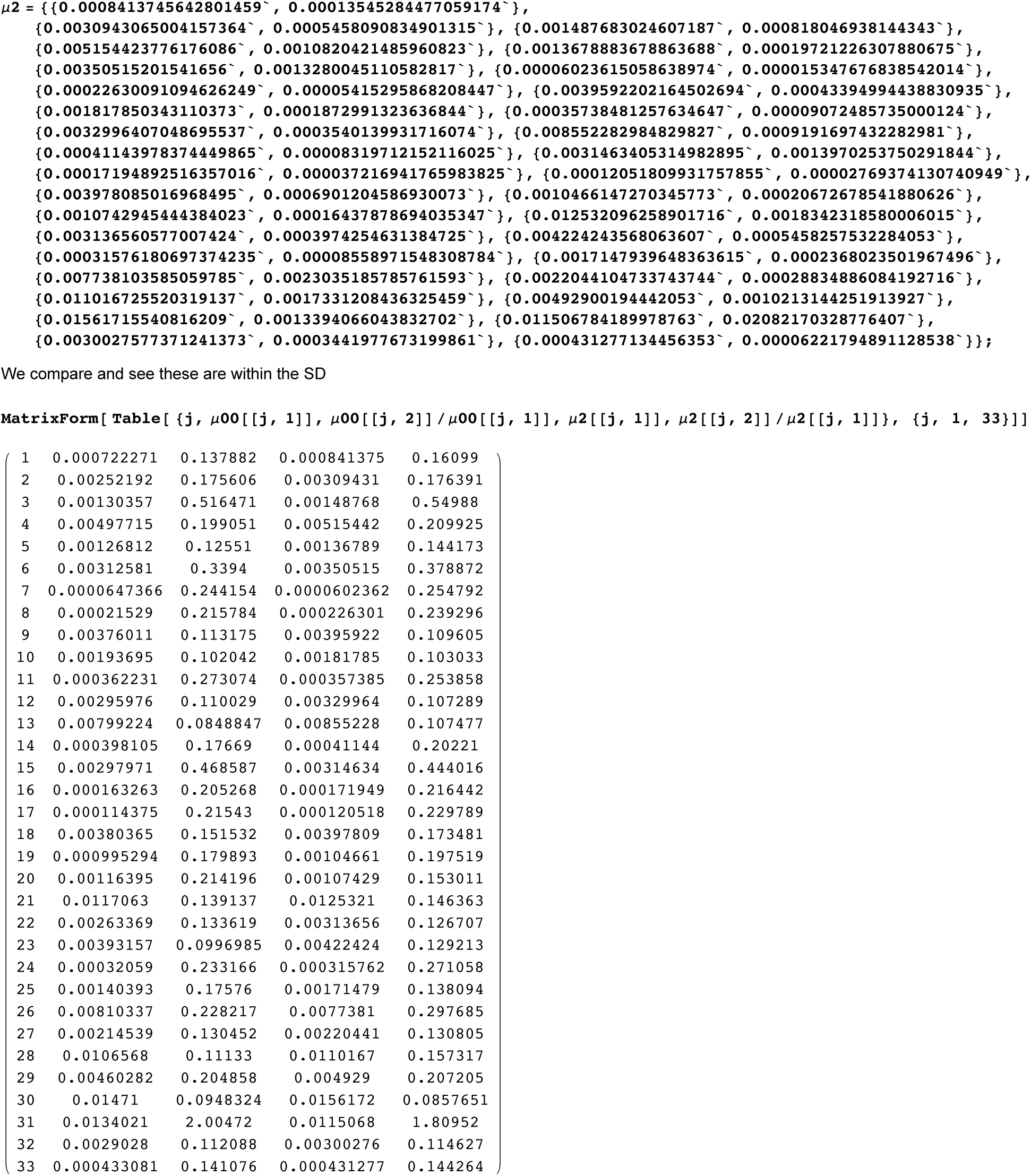

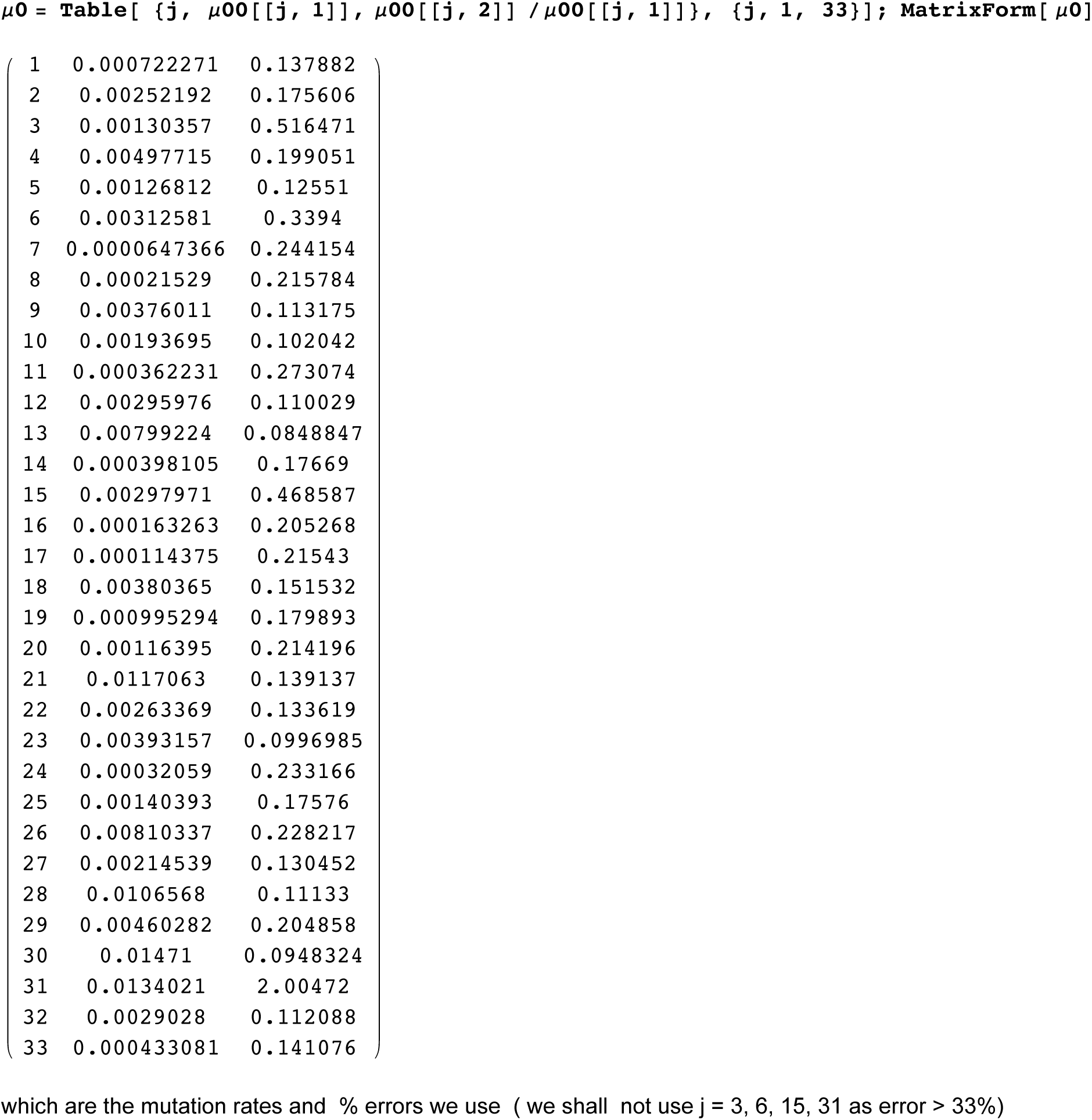

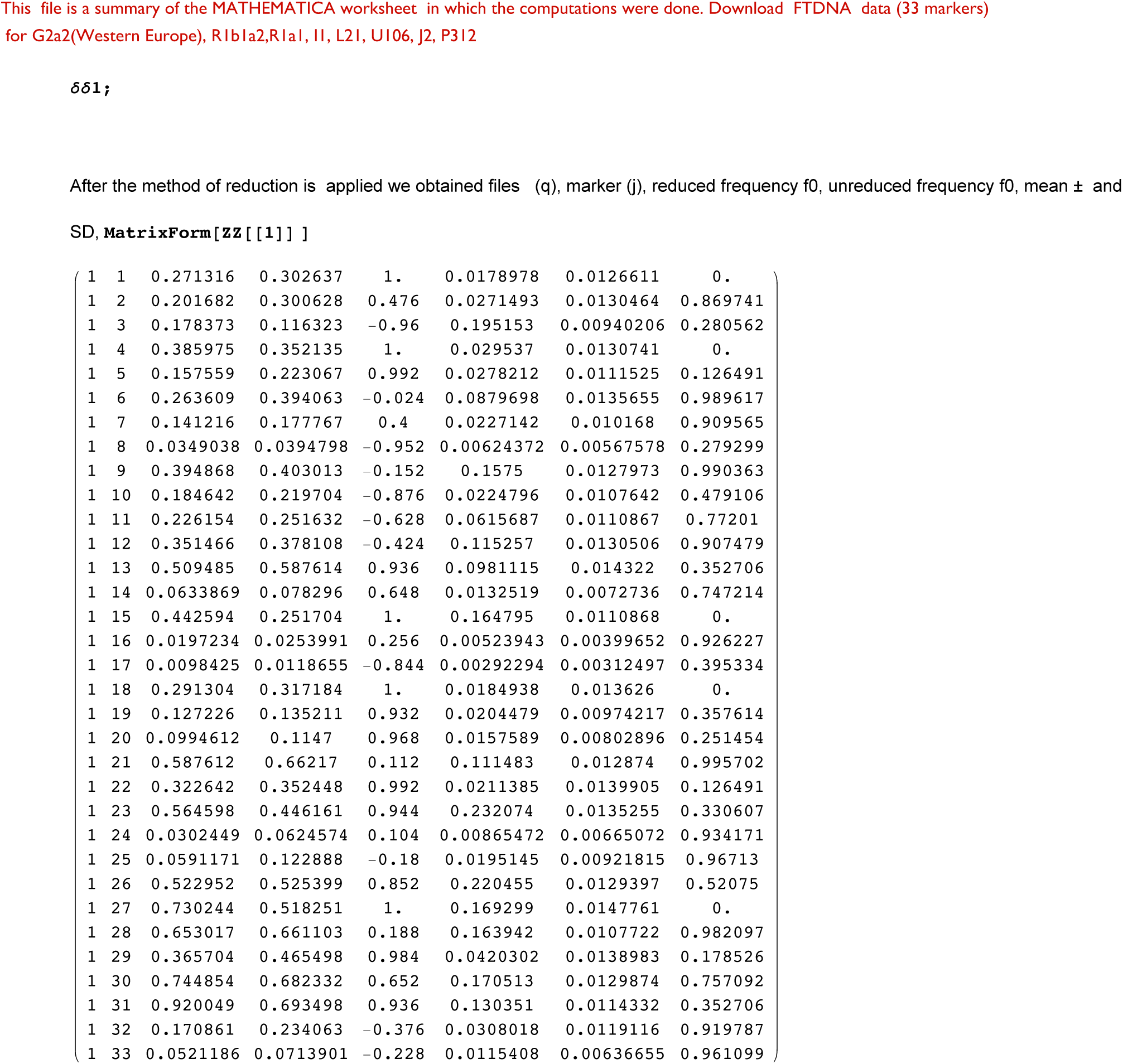

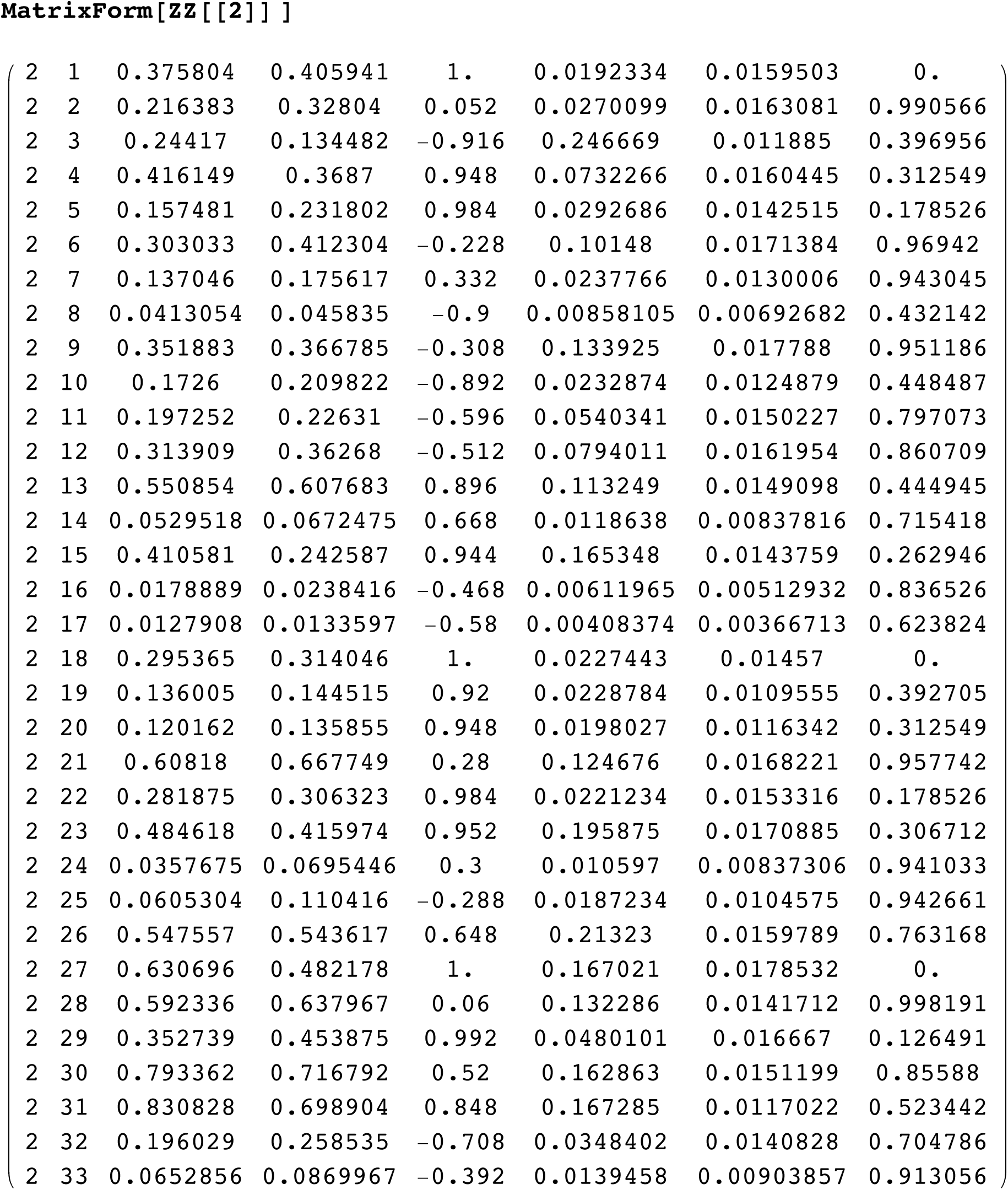

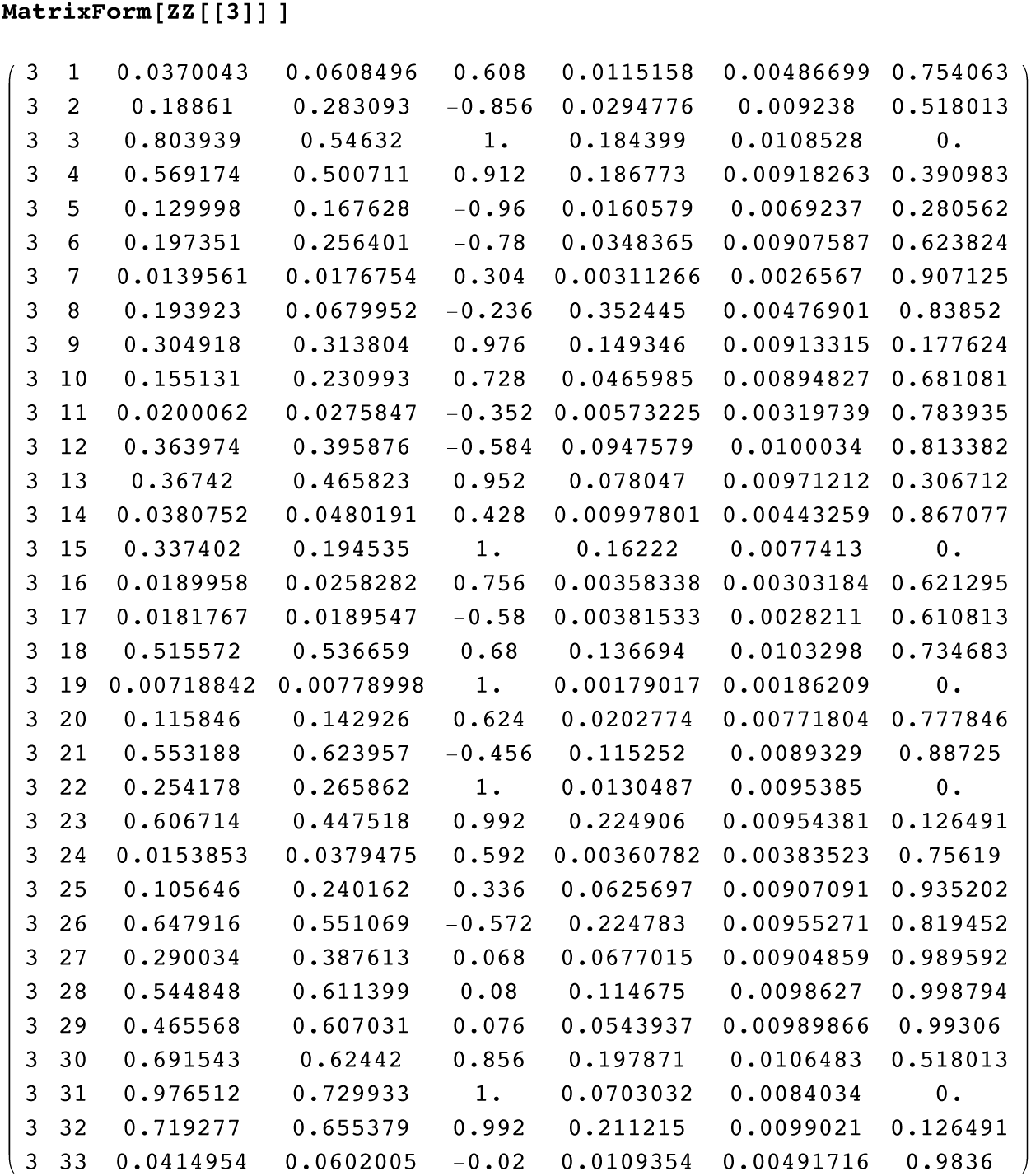

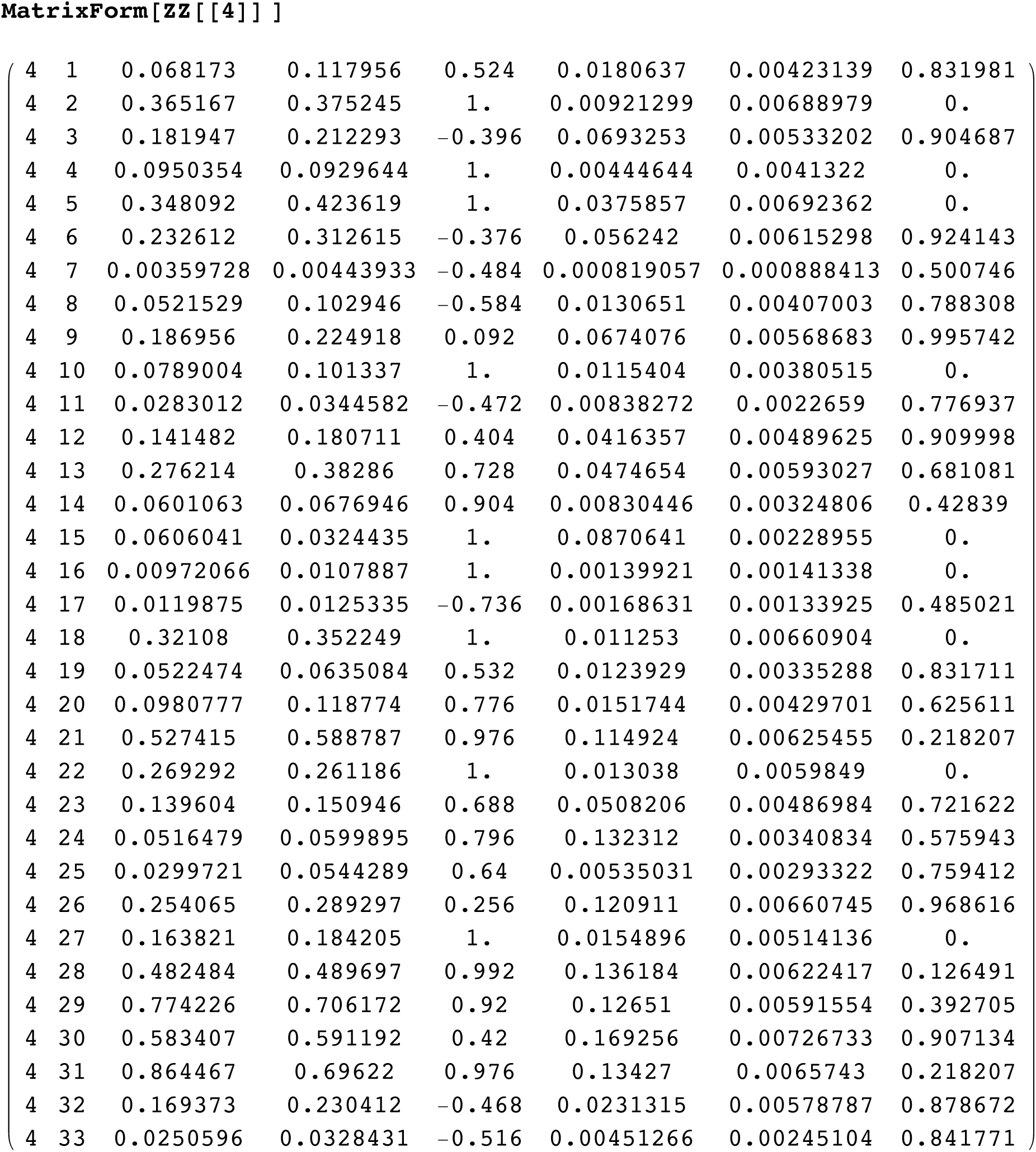

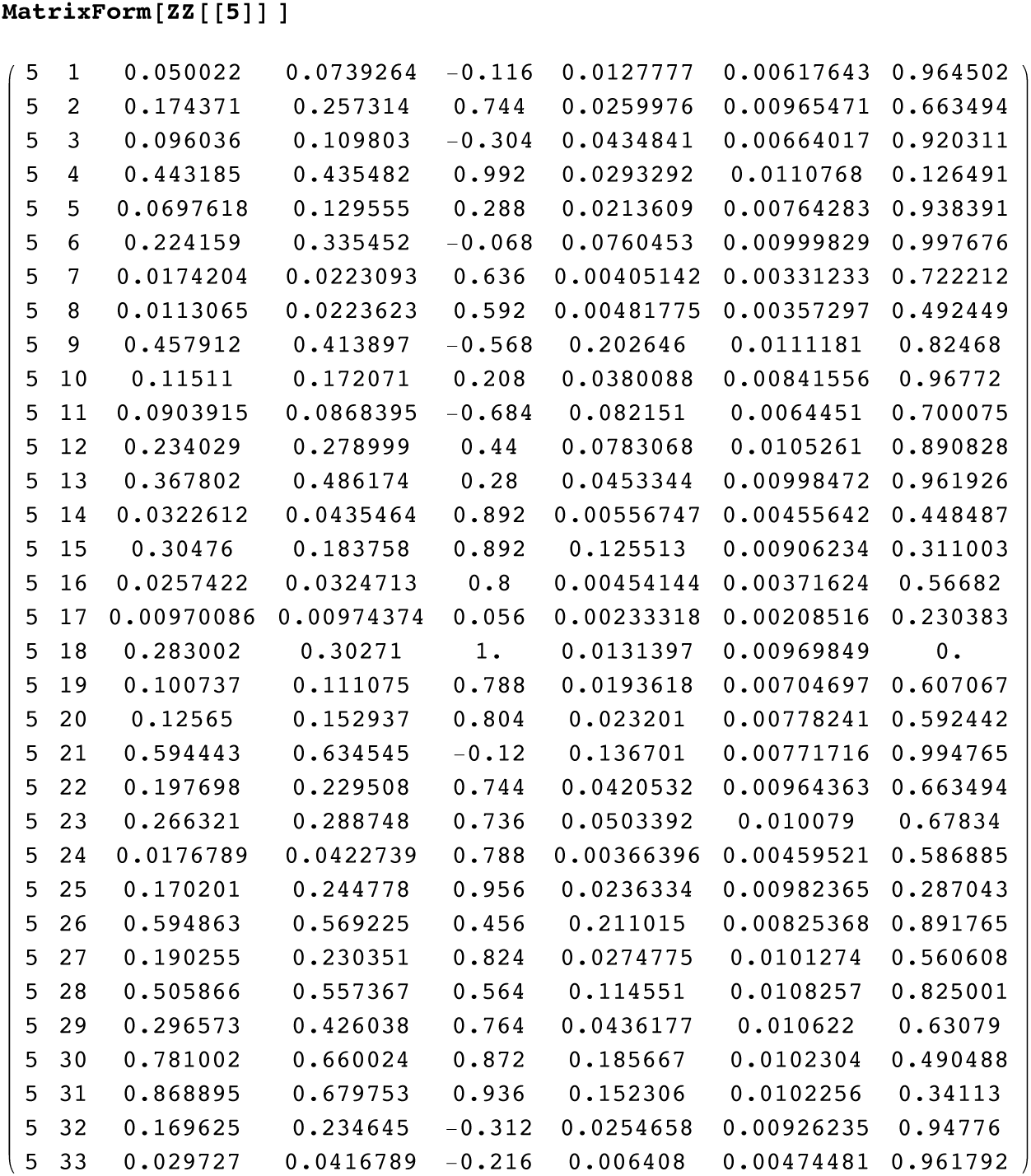

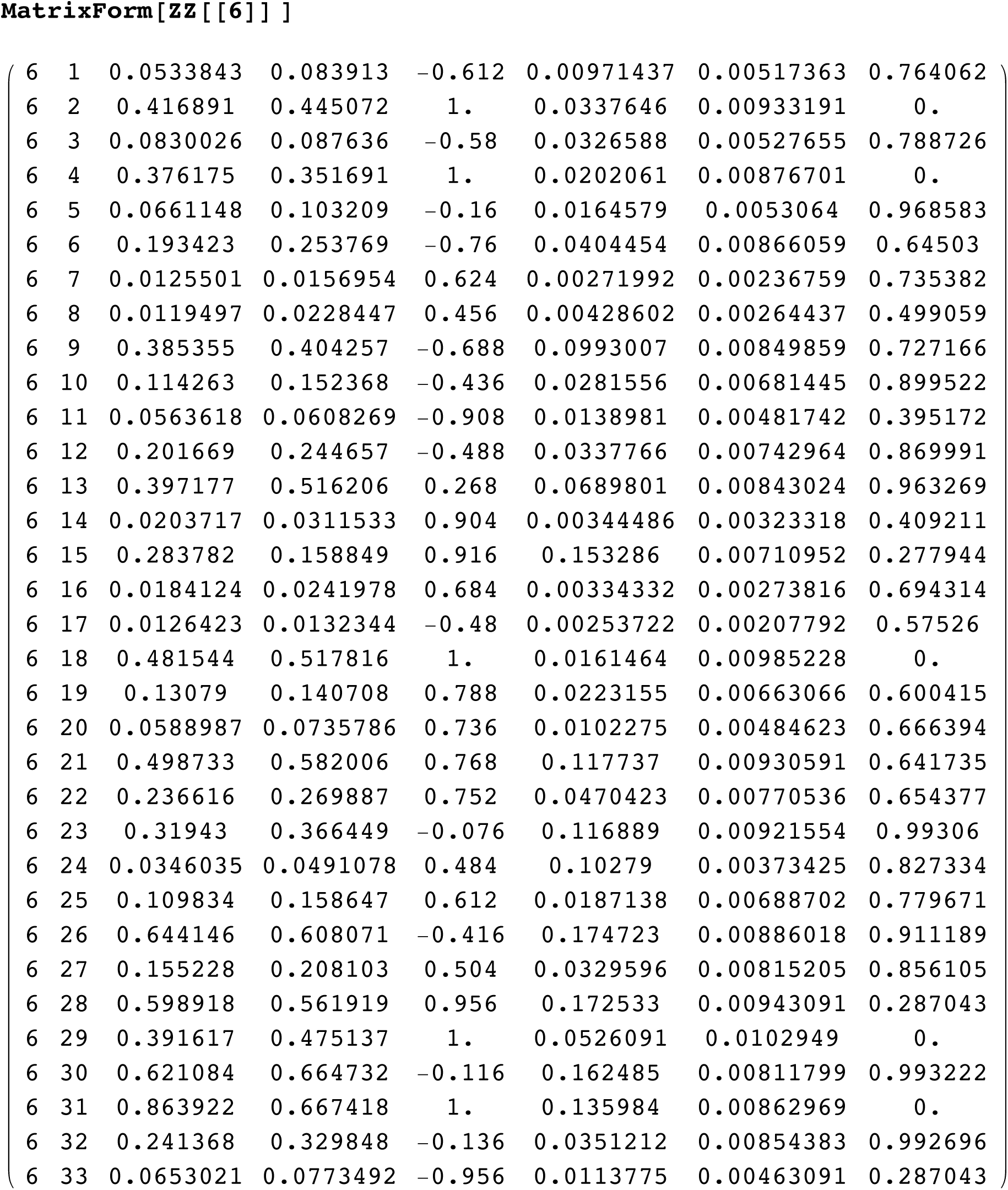

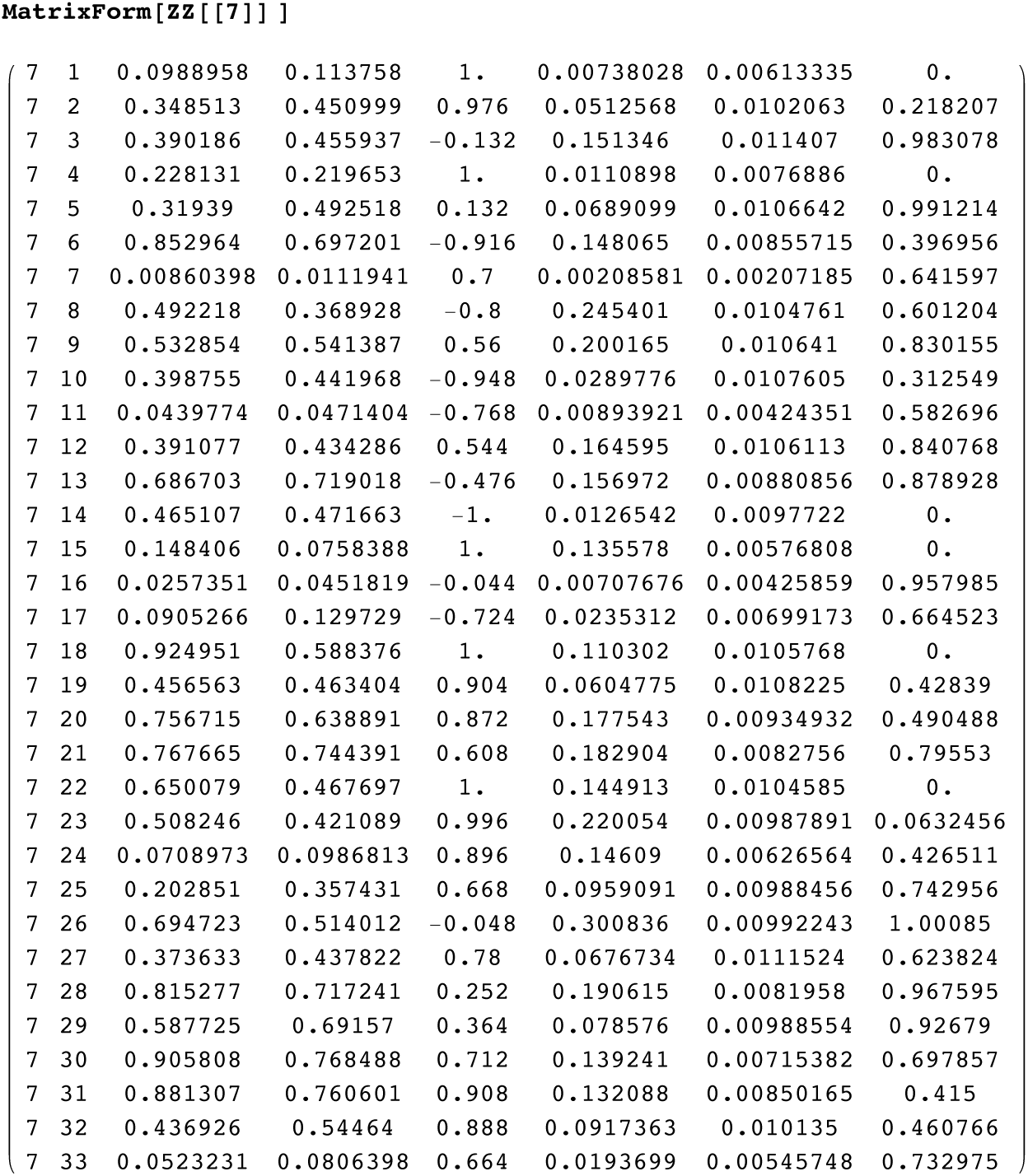

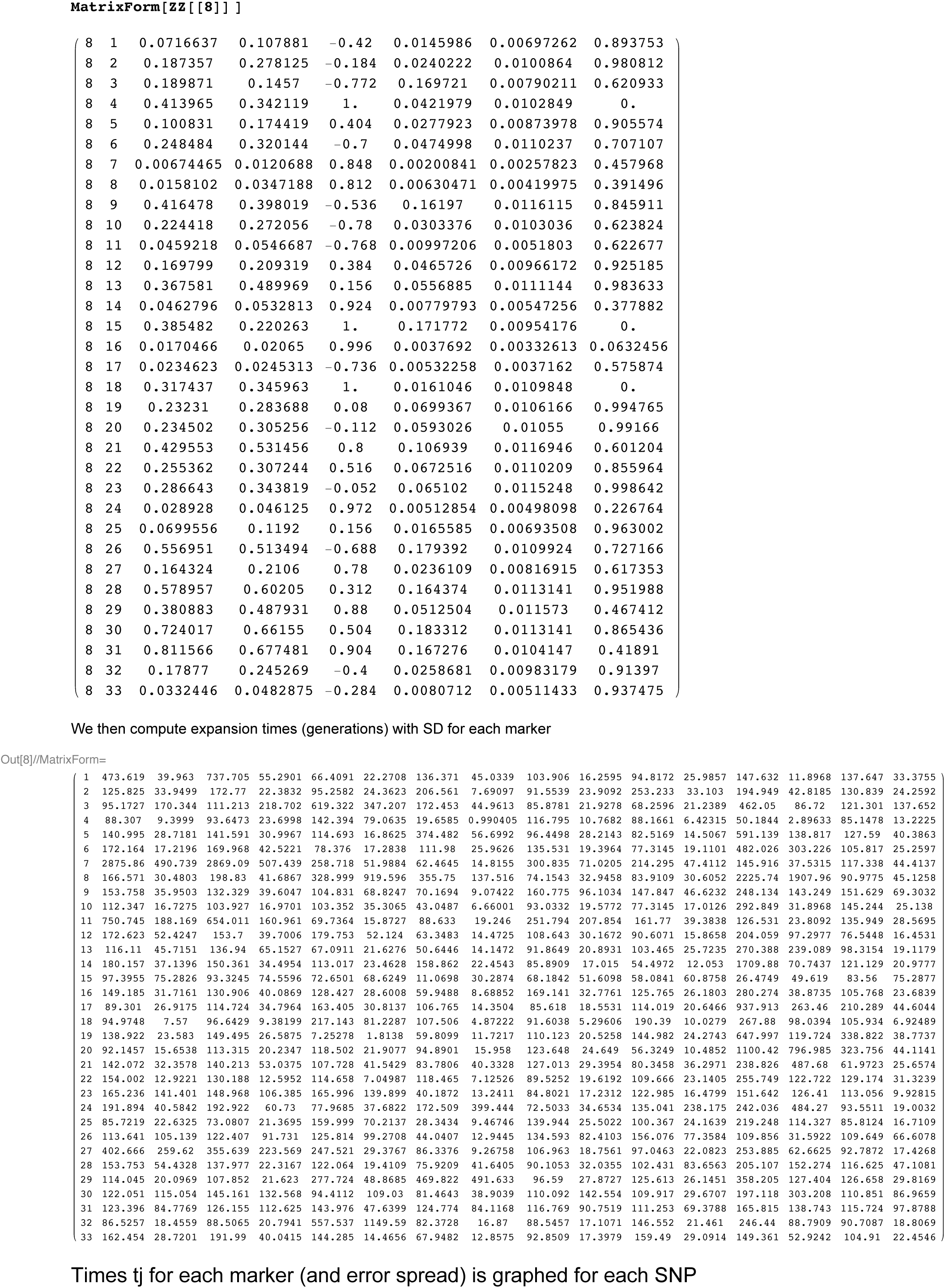

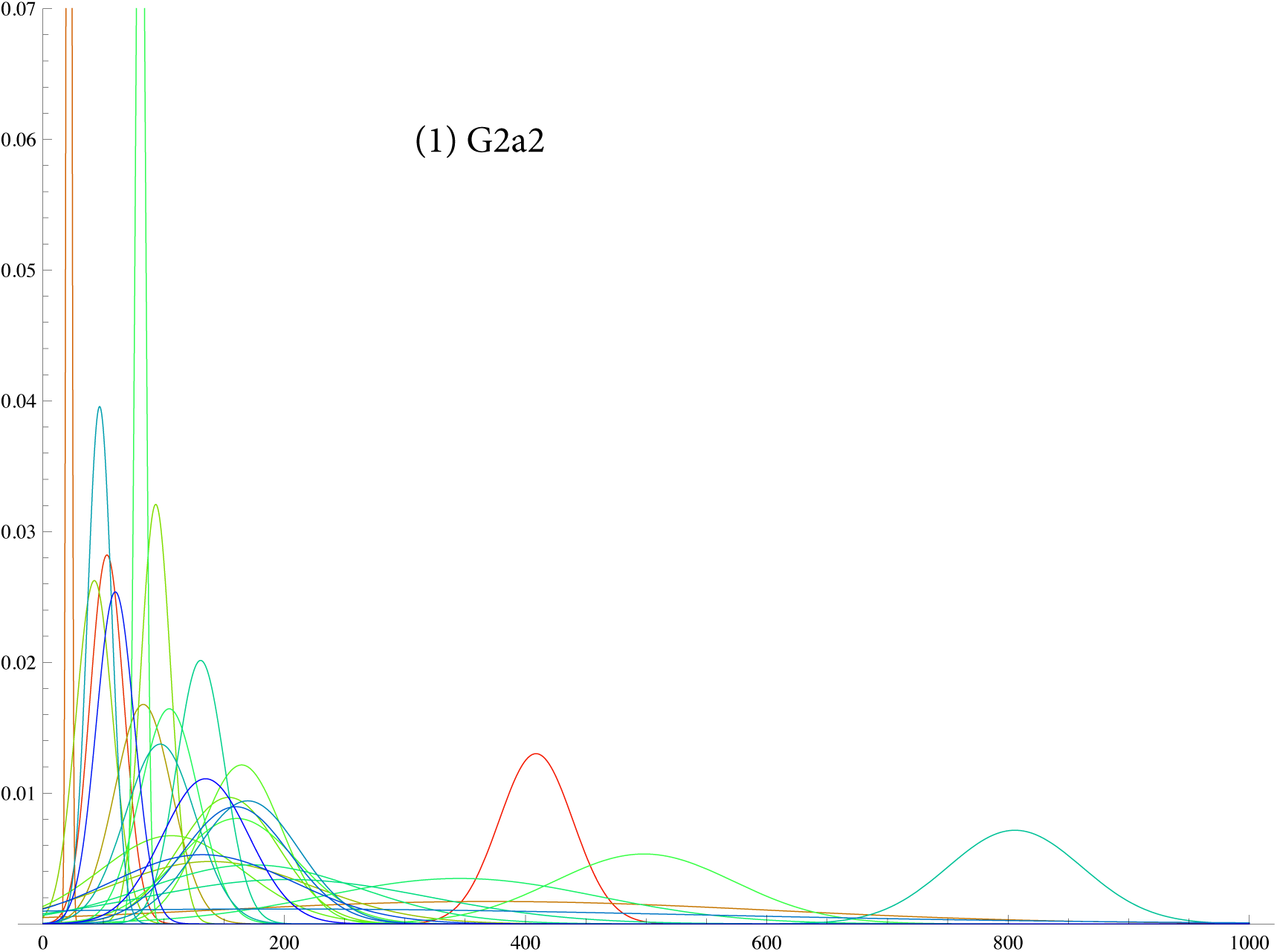

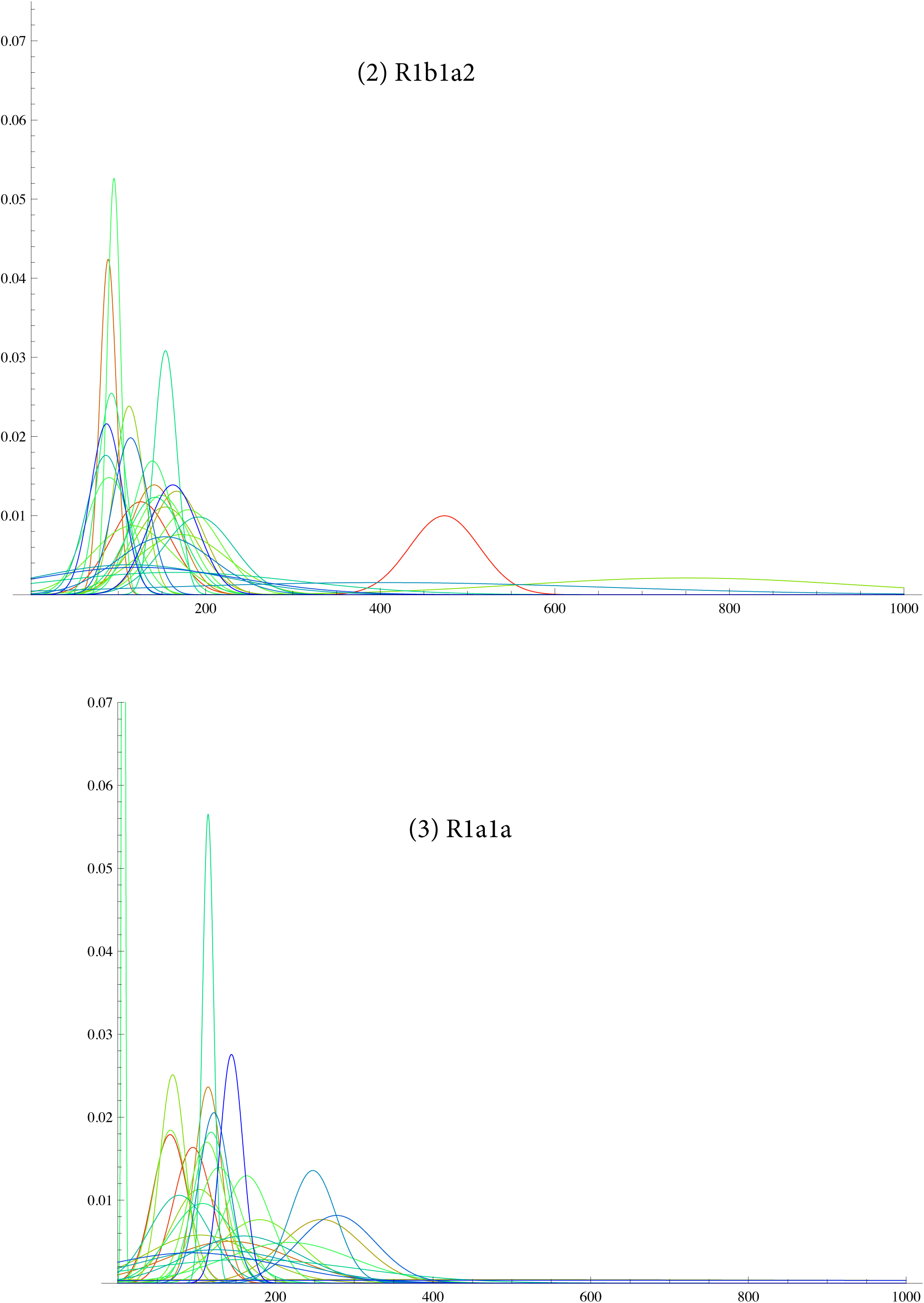

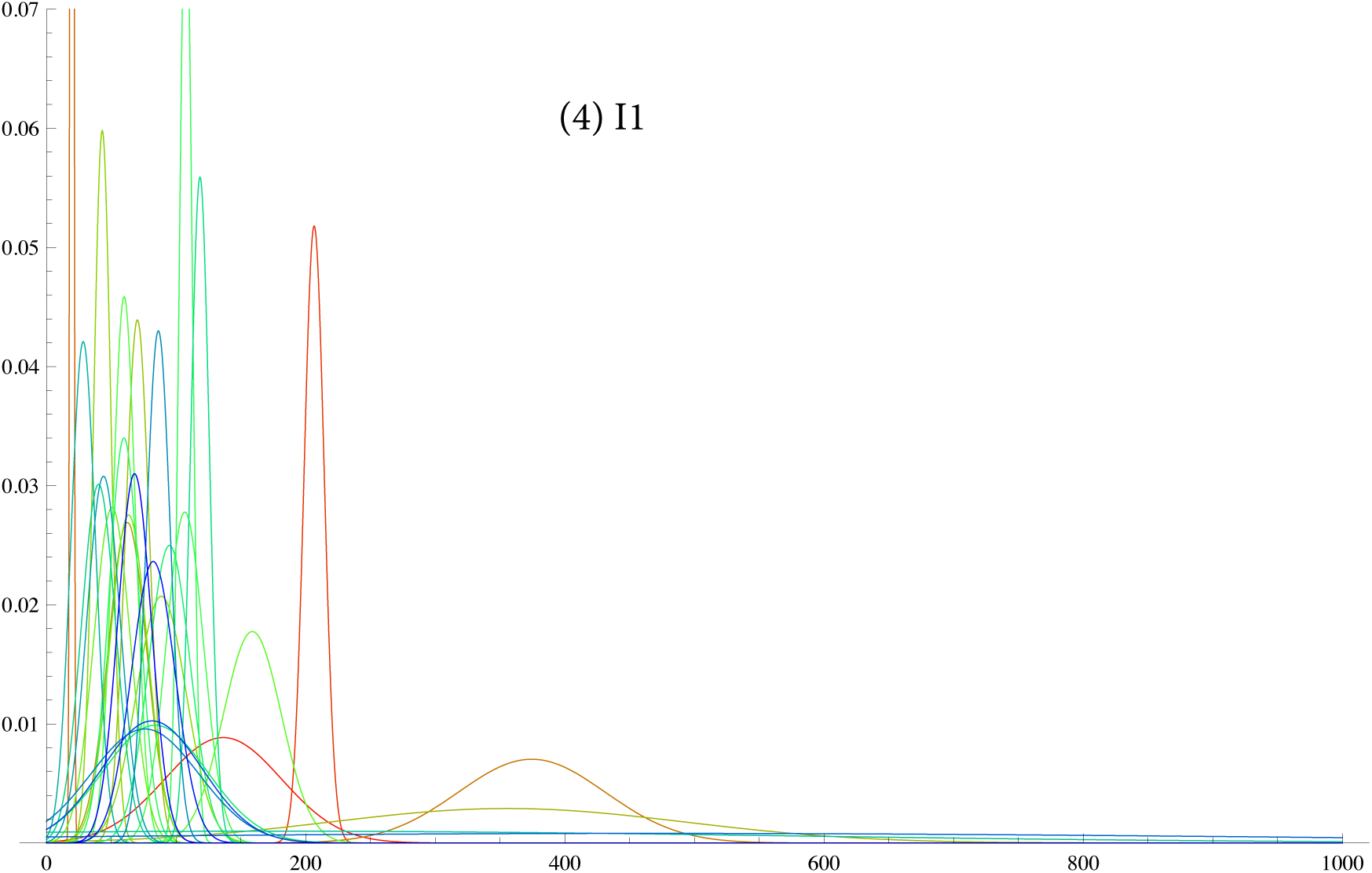

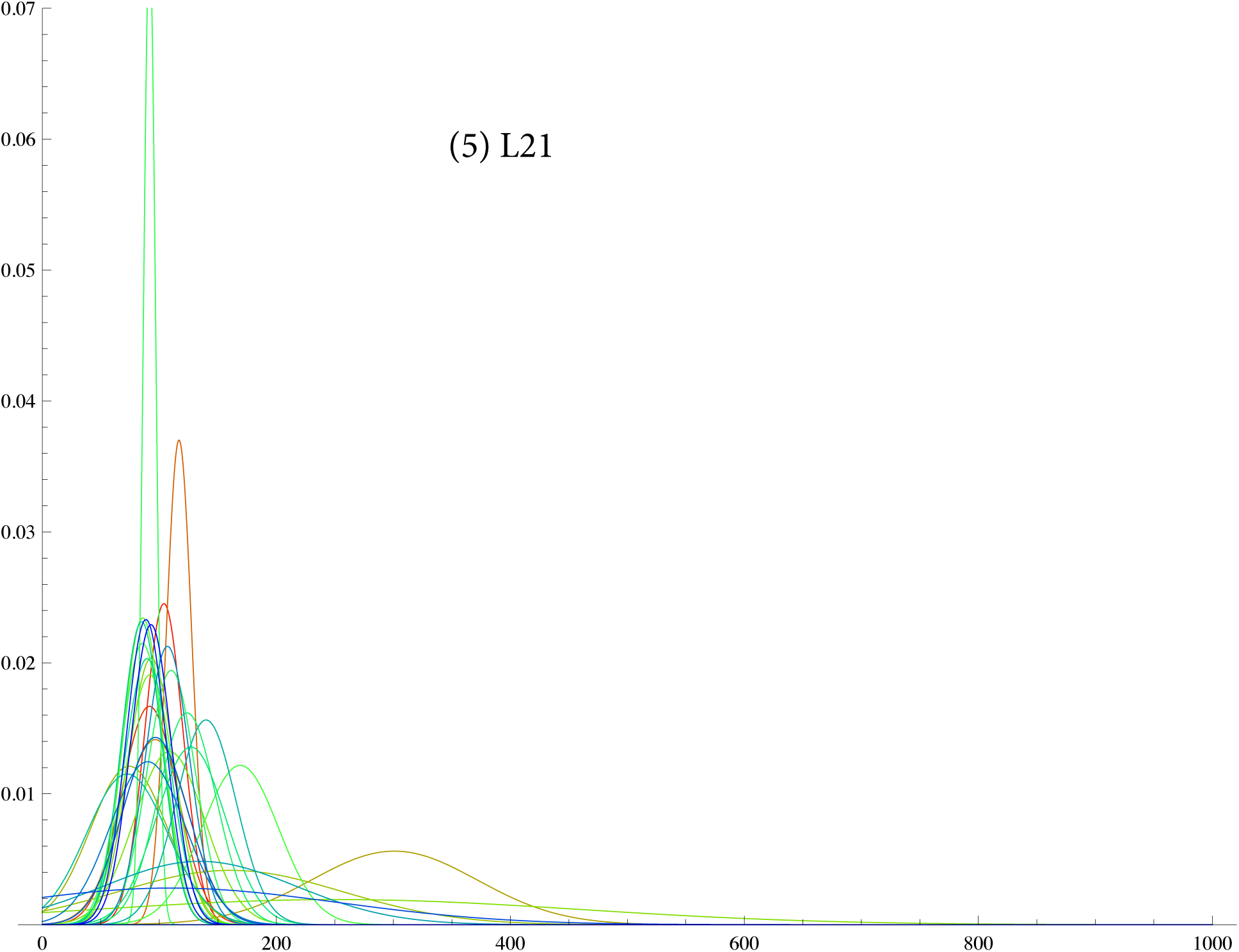

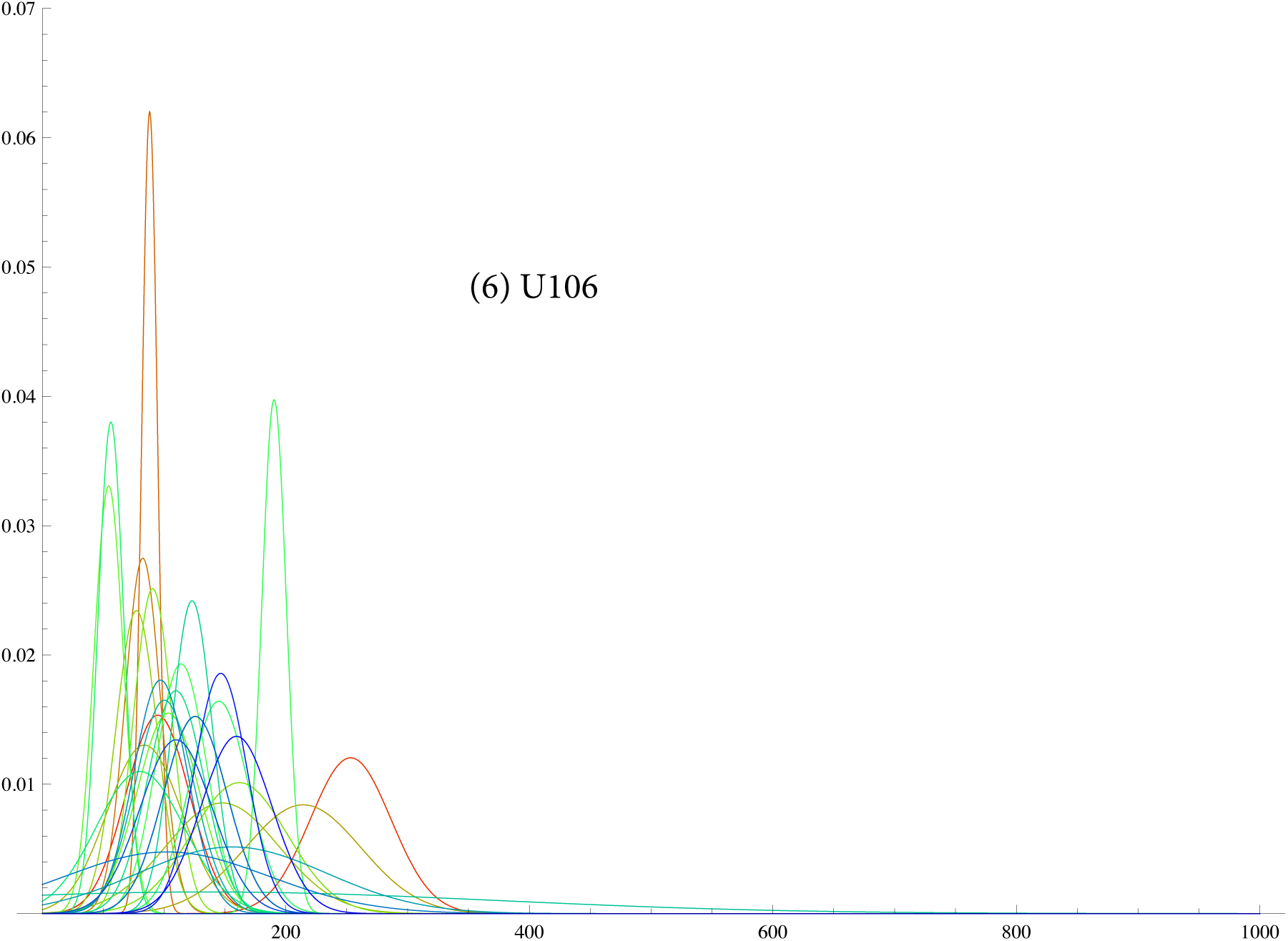

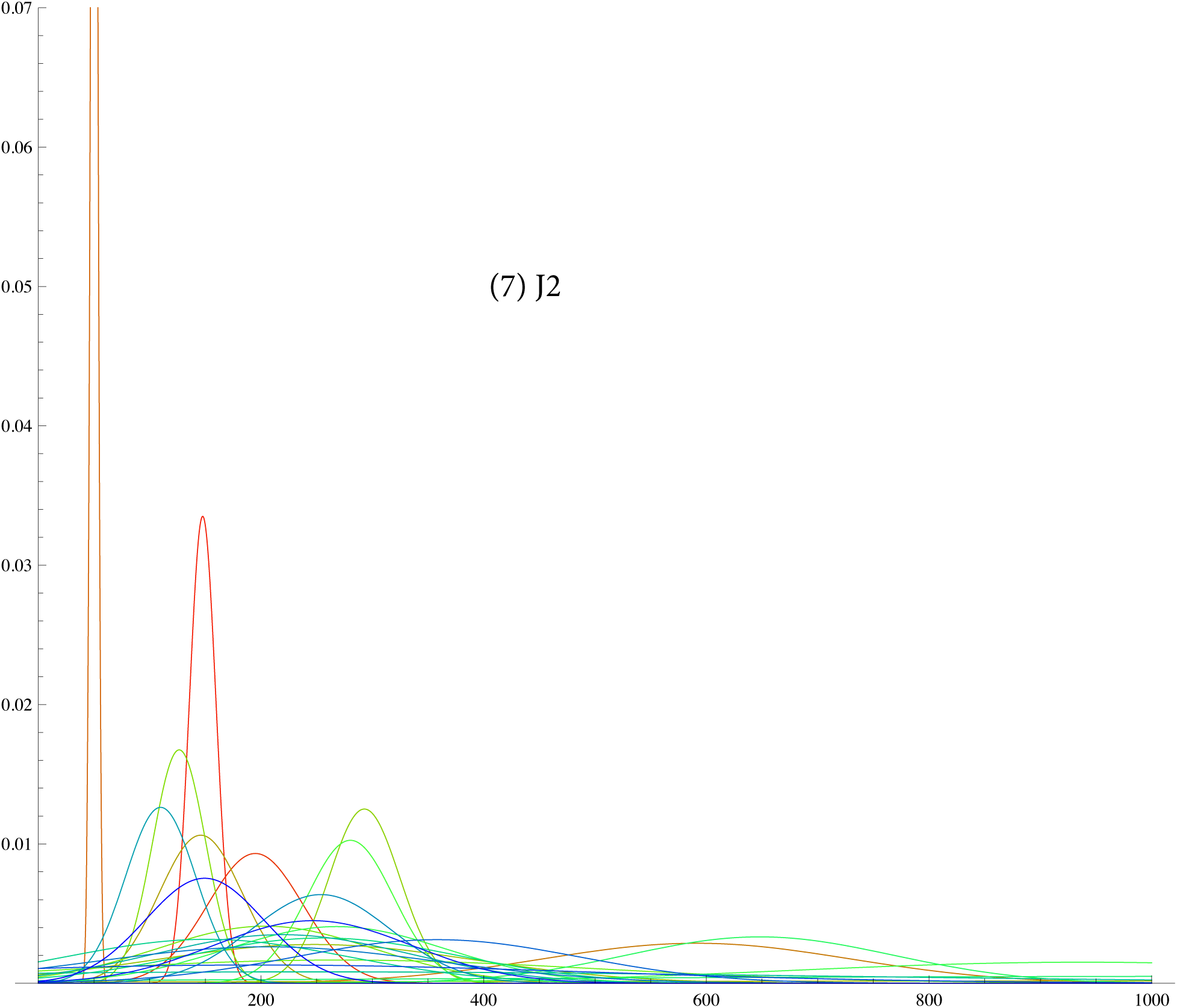

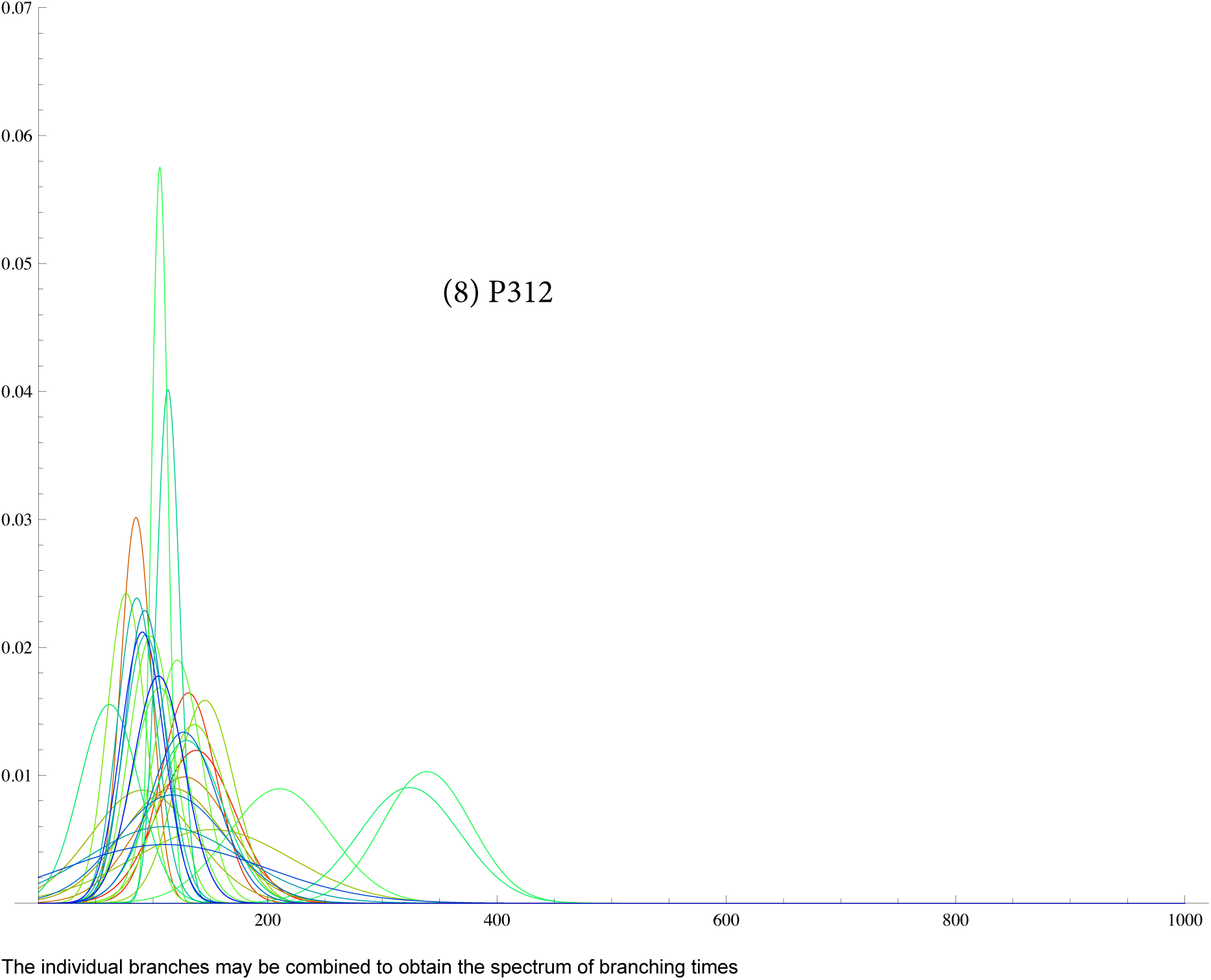

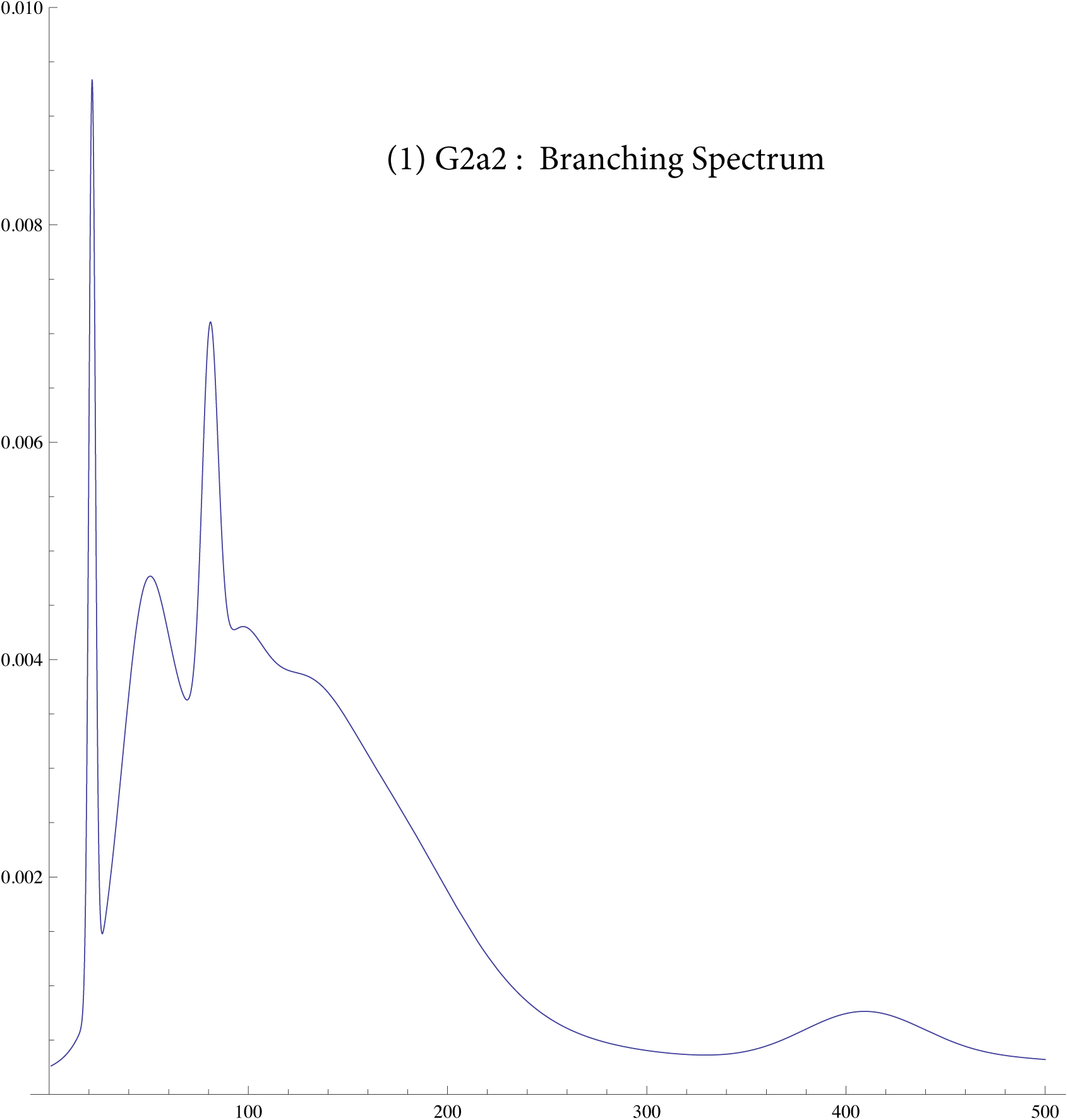

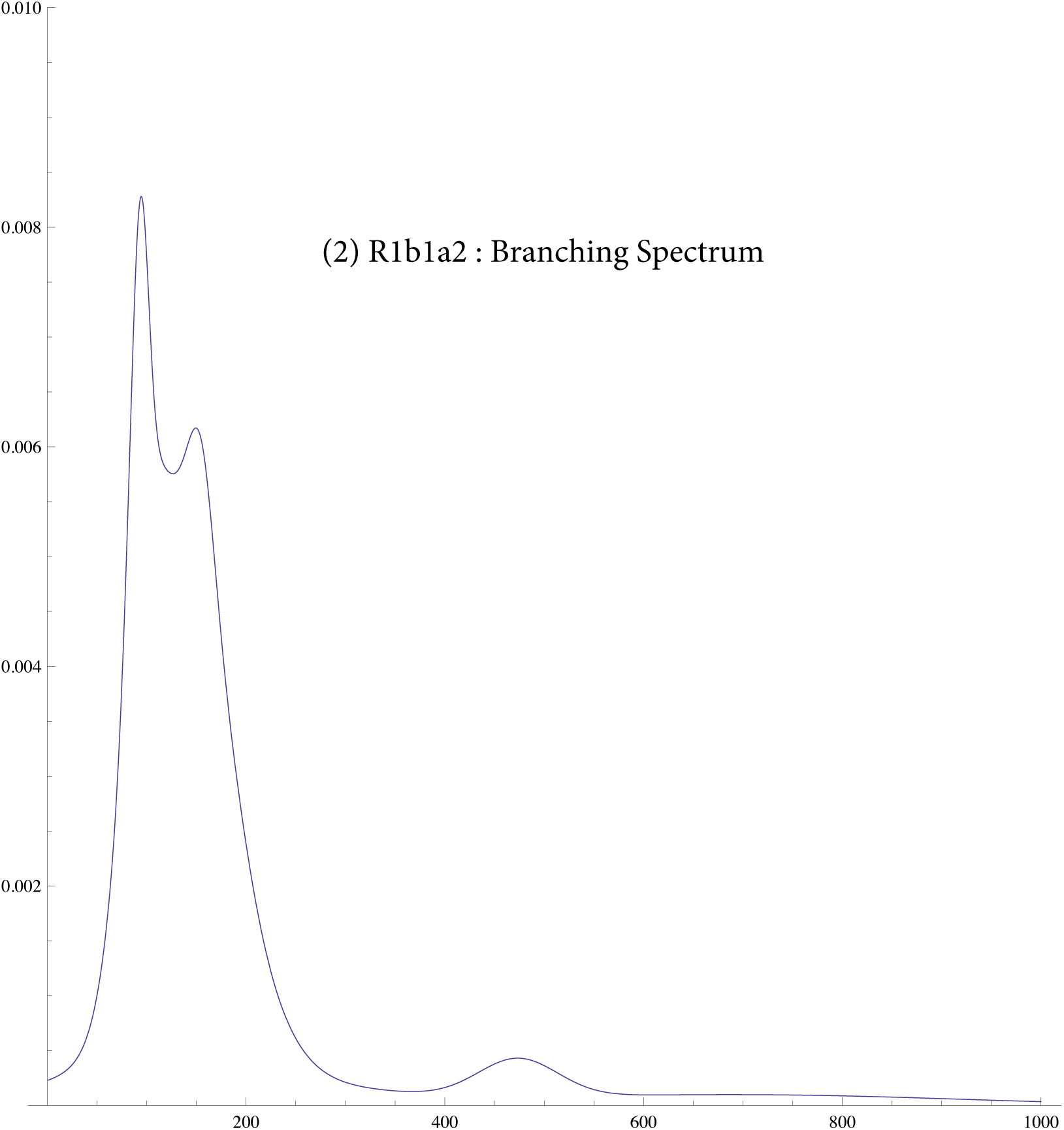

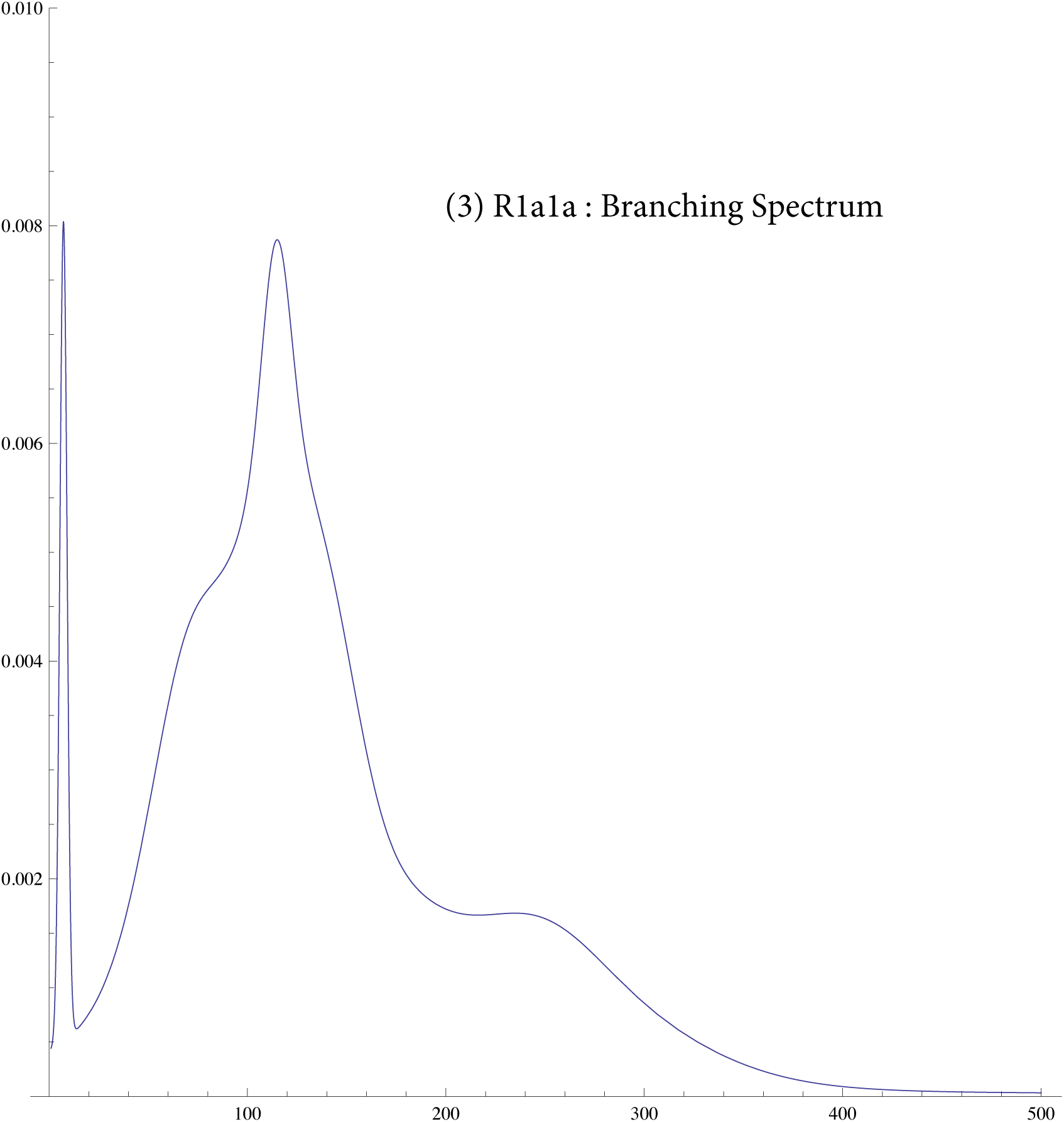

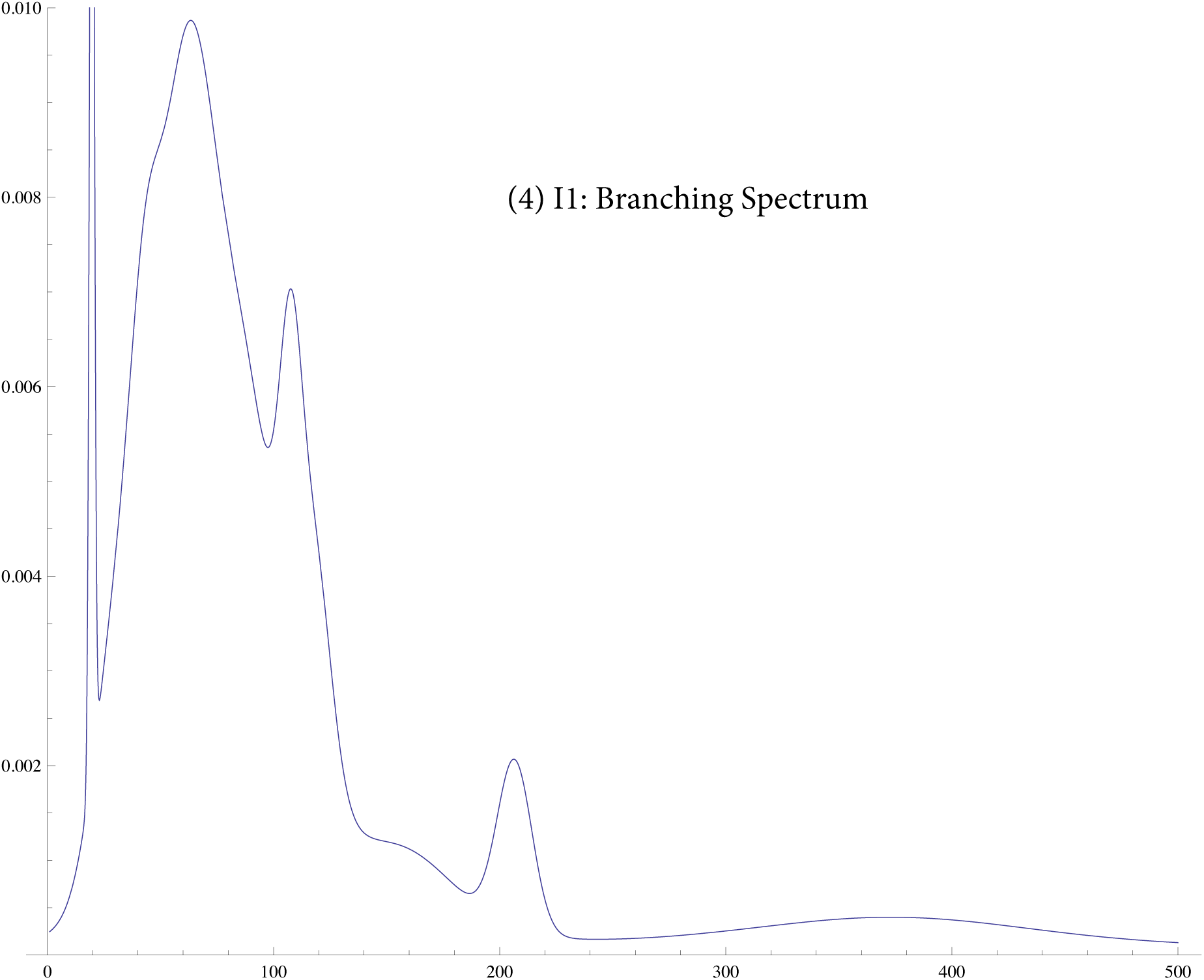

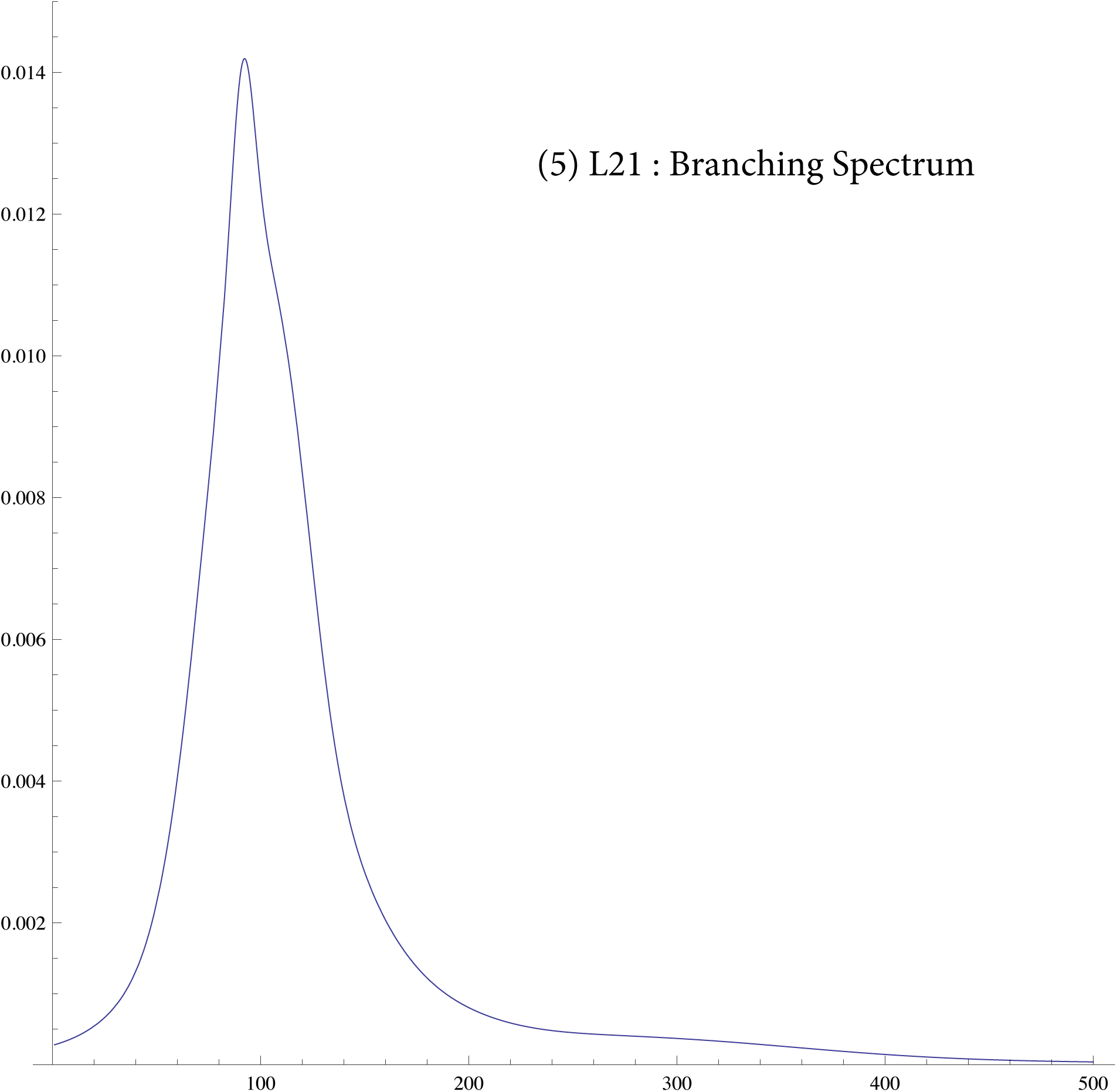

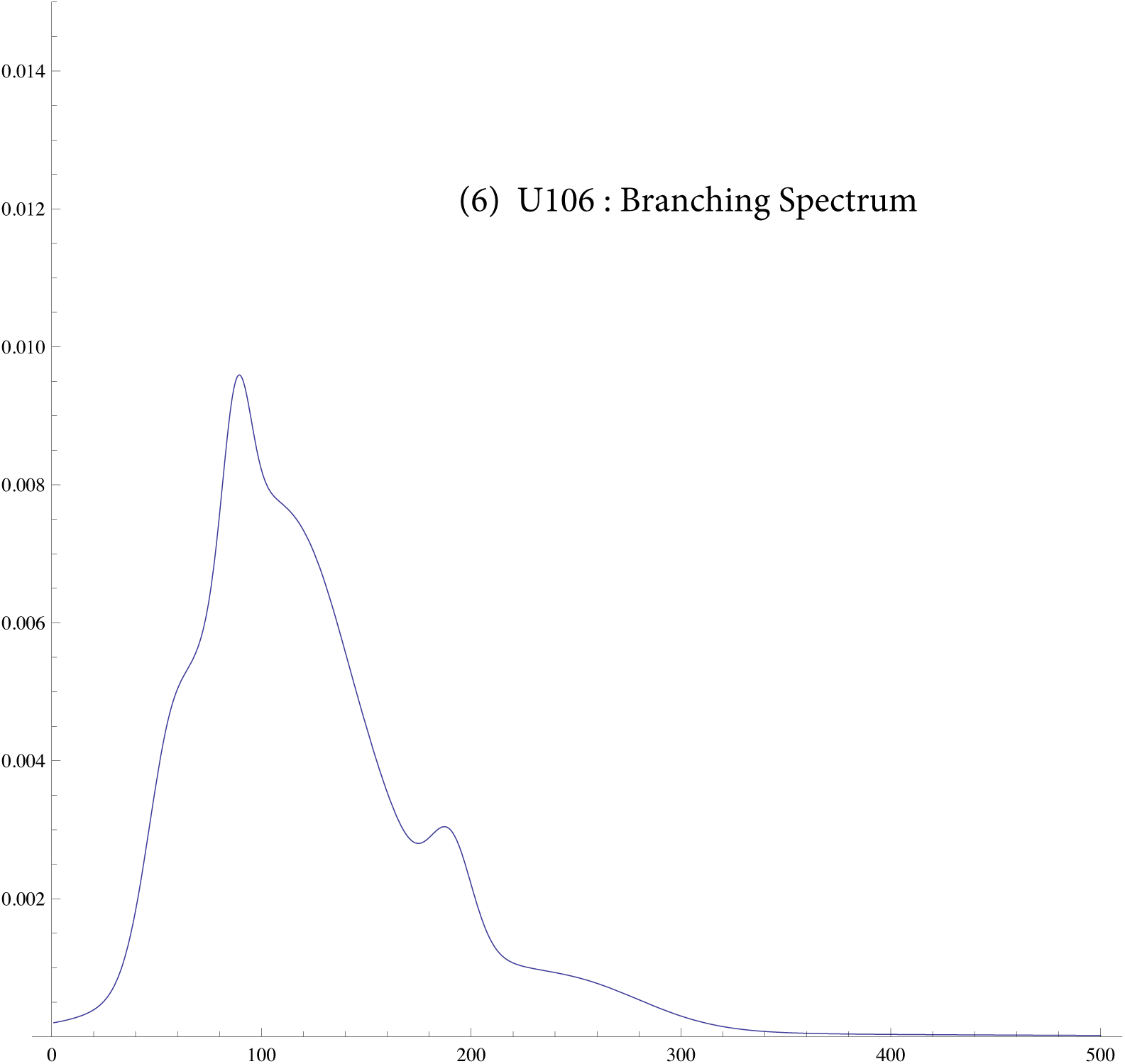

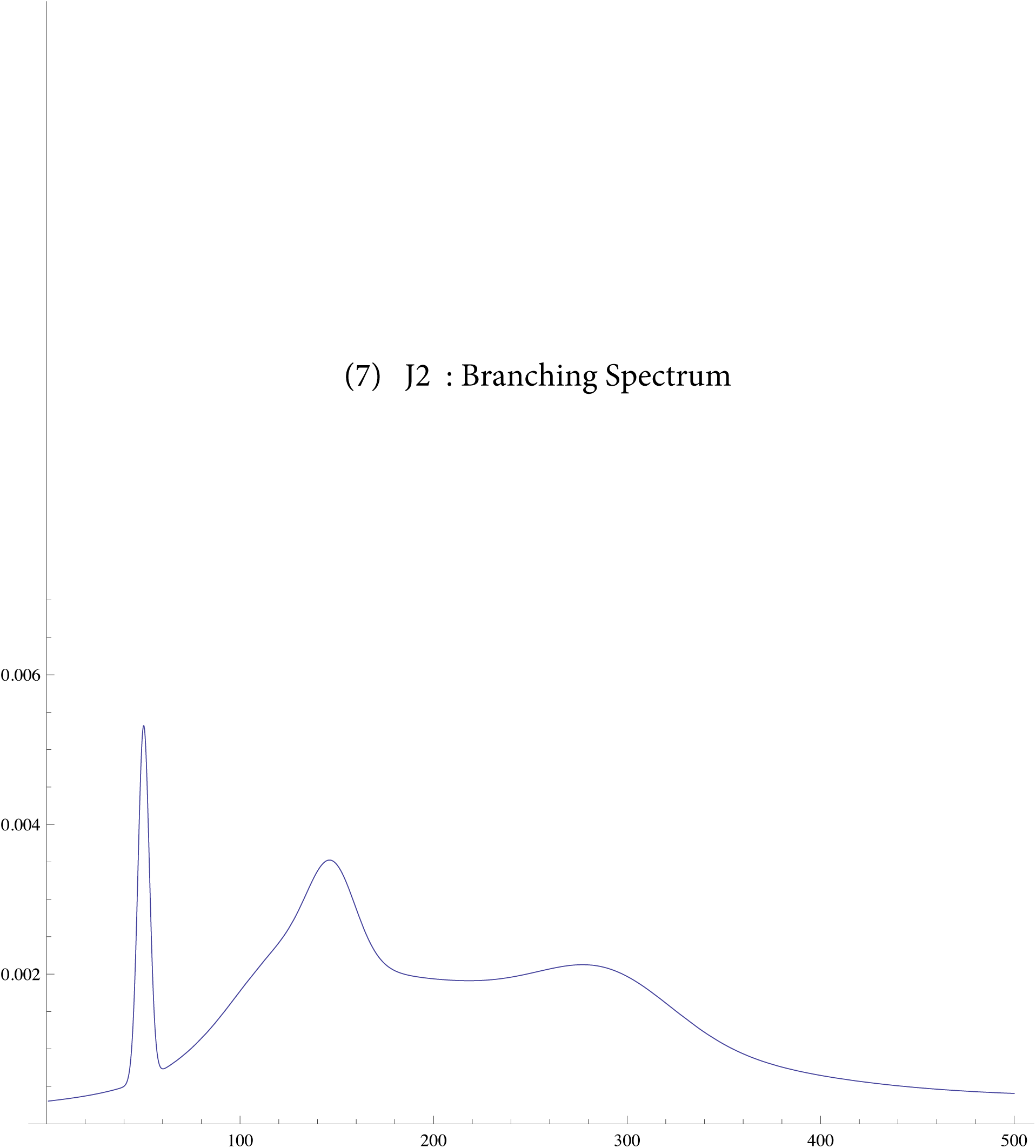

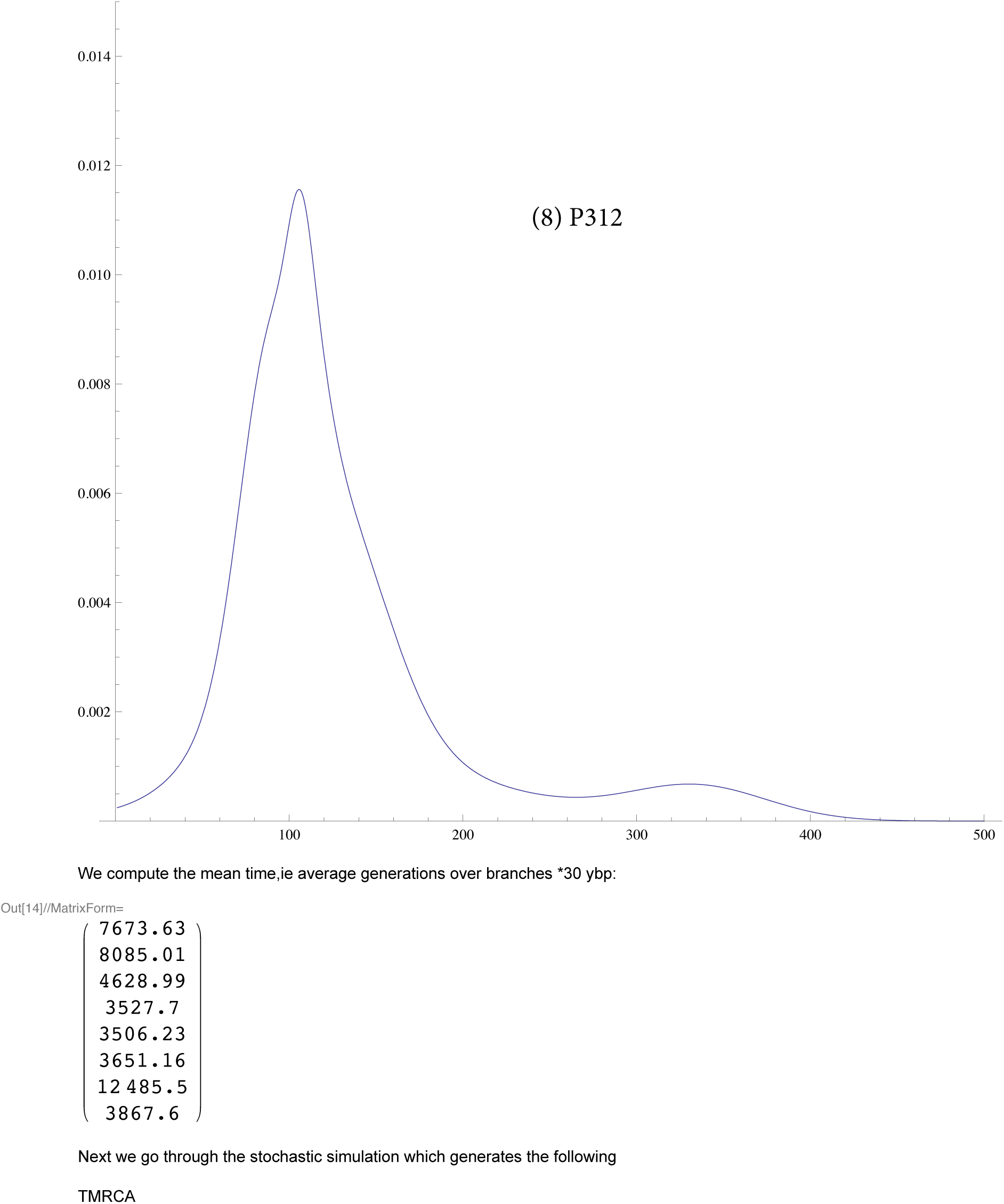

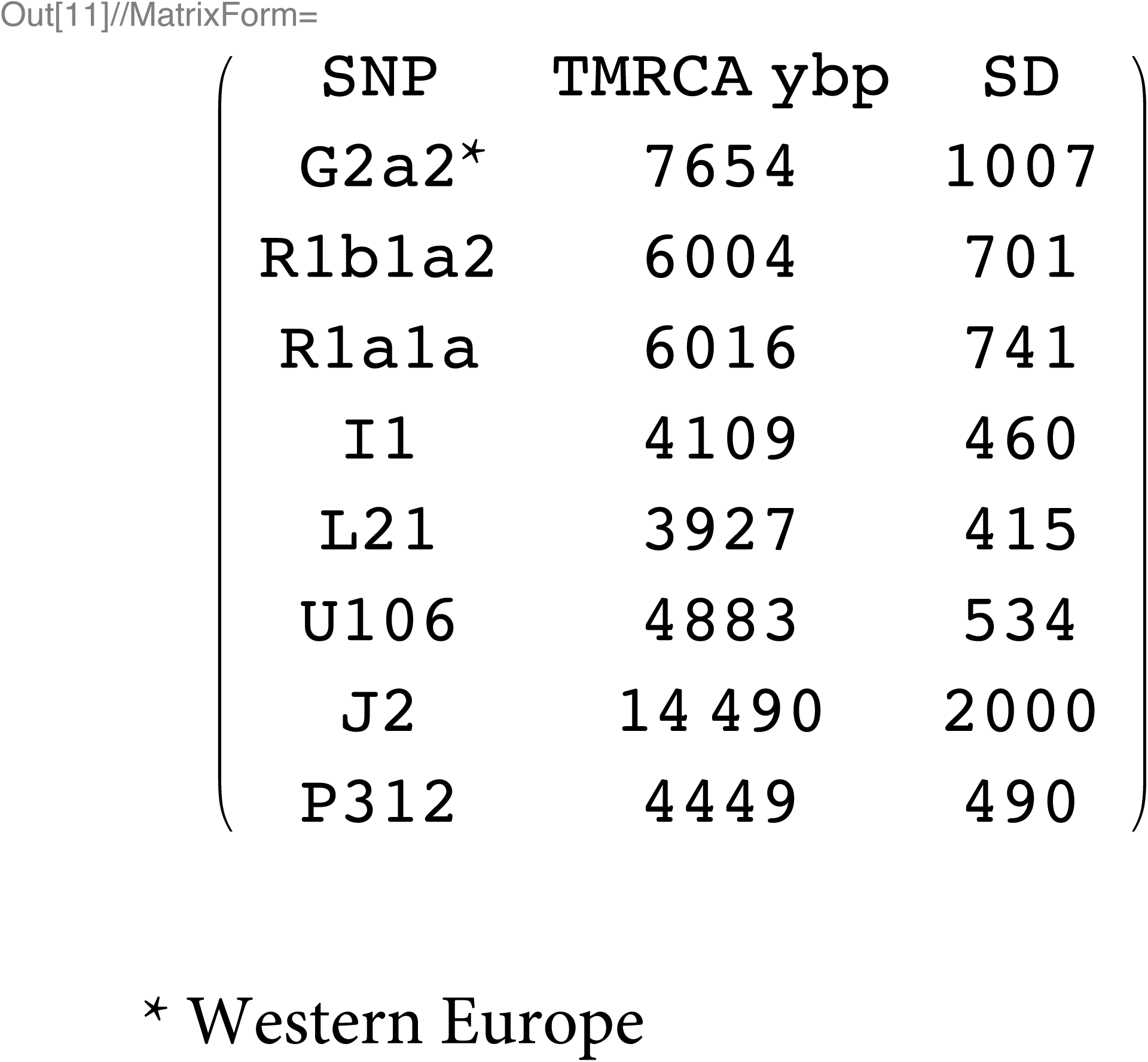

## SI 7: data for R1b1a2 for DYS j = 1, ..33

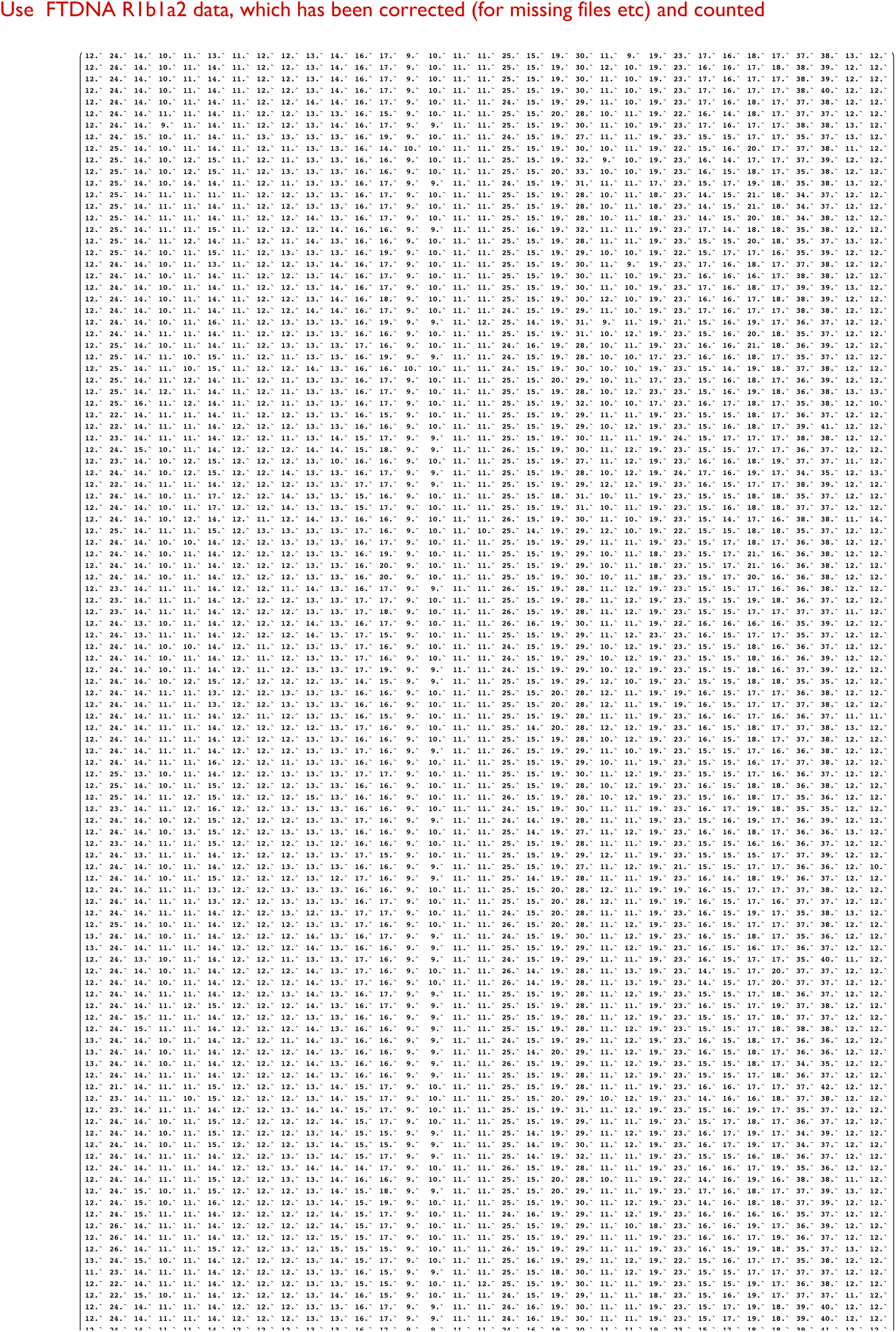

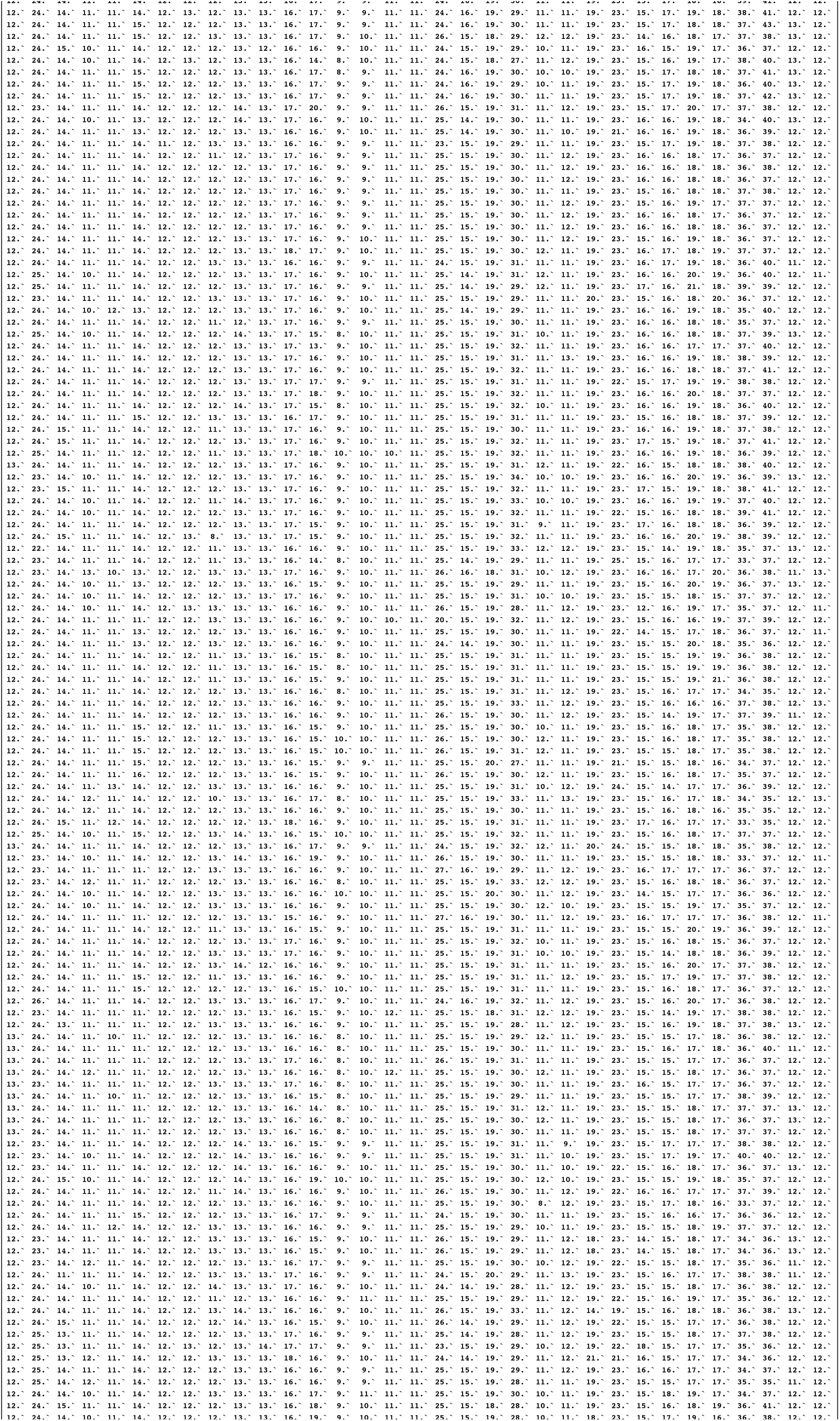

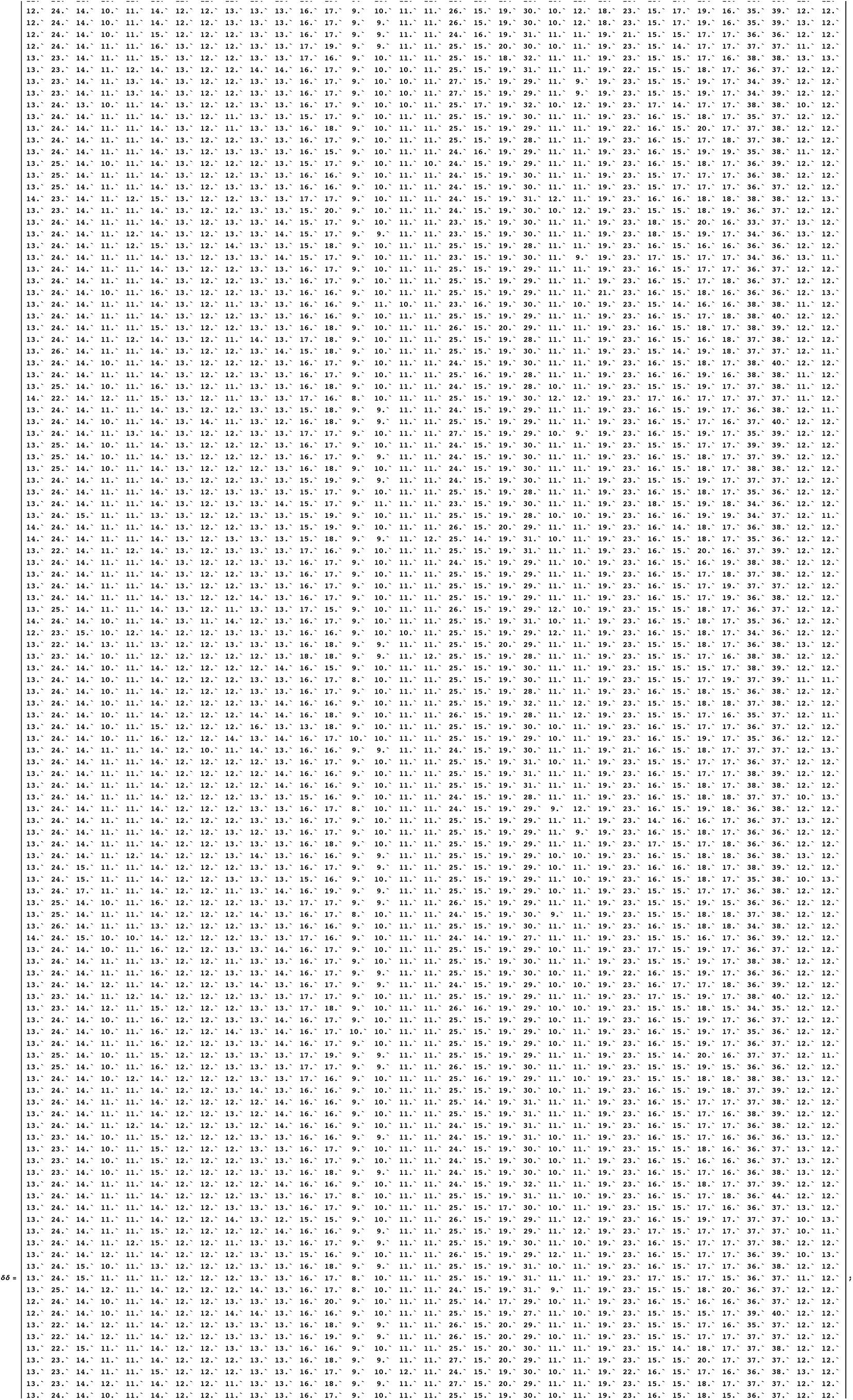

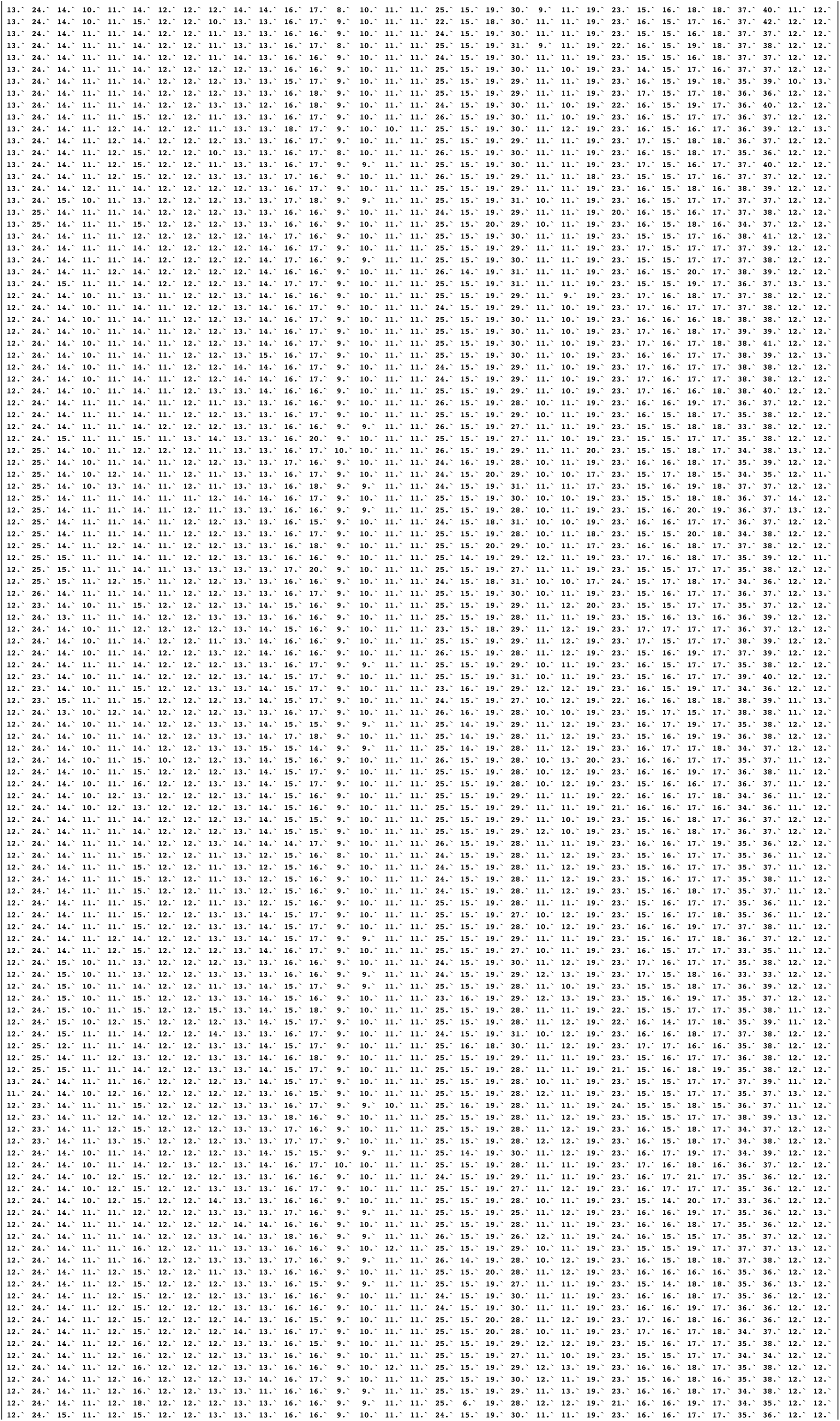

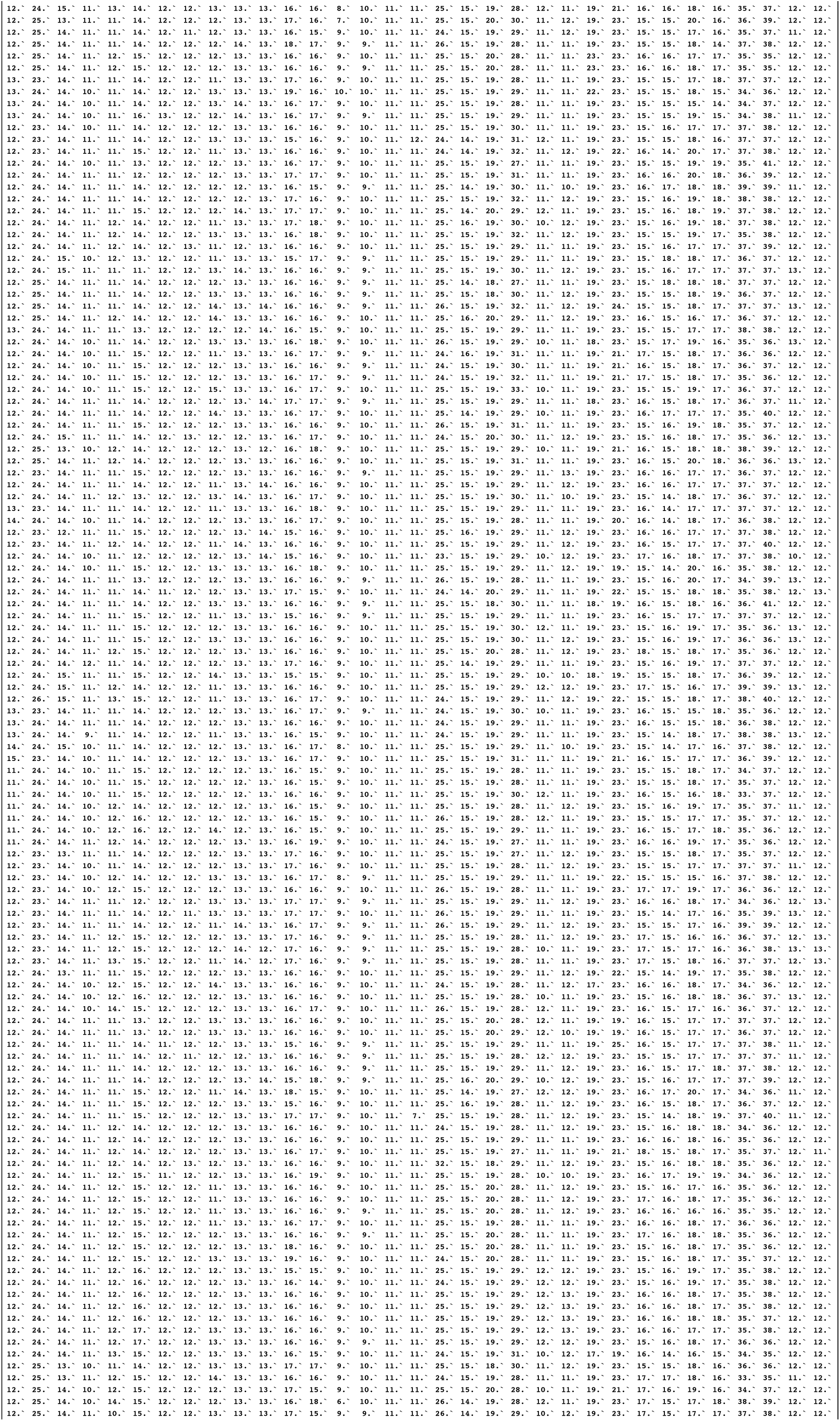

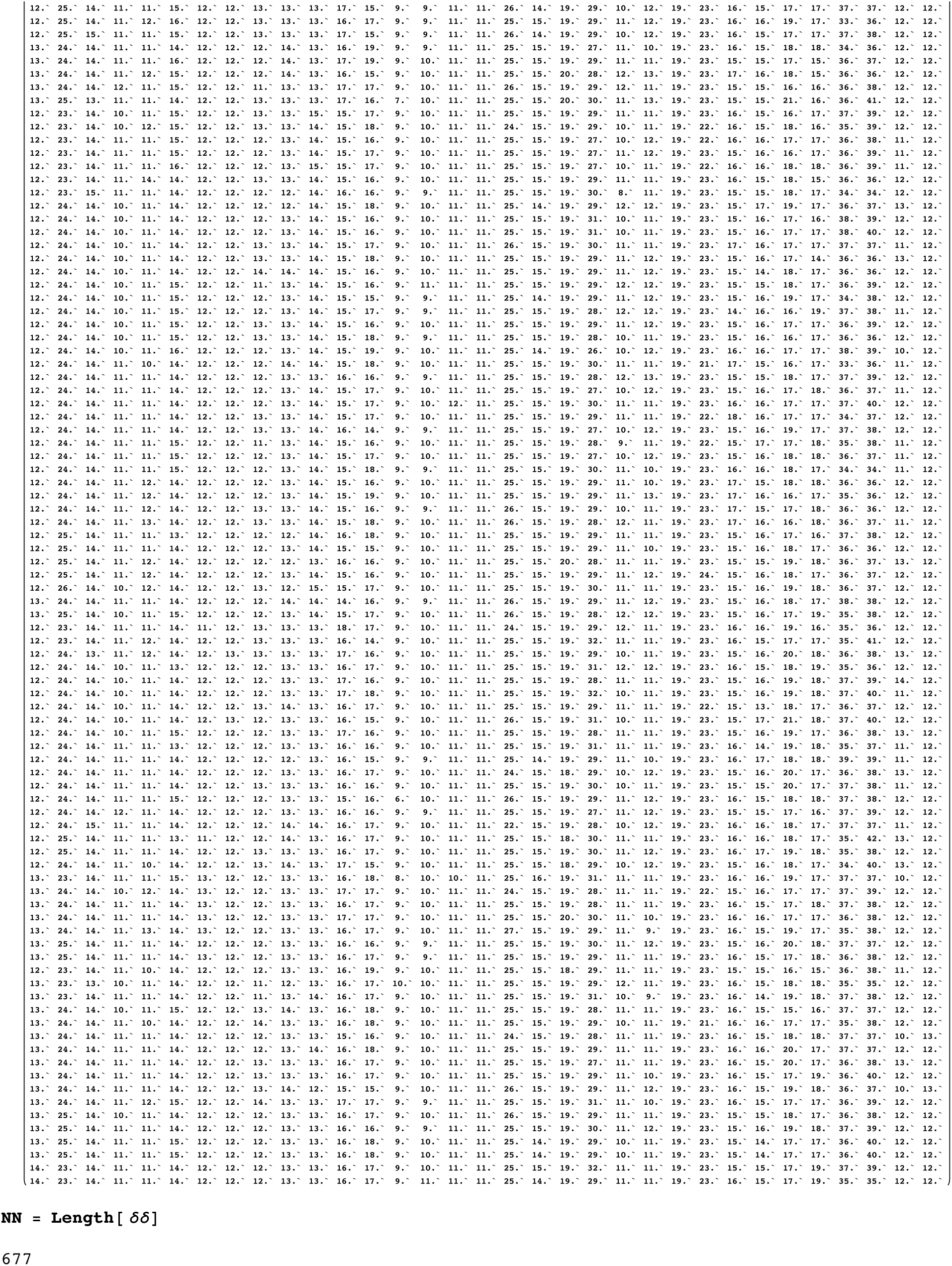

